# Genetically encoded RNA strand exchange circuits for programmable protein expression and information processing in cells

**DOI:** 10.1101/2025.09.24.678369

**Authors:** August O. Staubus, Fernanda Piorino, Olga B. Vasilyeva, Molly E. Wintenberg, John M. Hurley, Nina Y. Alperovich, Samuel W. Schaffter

## Abstract

Nucleic acid circuits based on toehold-mediated strand exchange (TMSE) have enabled sophisticated *in vitro* molecular computers, biosensors, and diagnostics, but translating these successes to living cells for continuous operation has been challenging. Primary hurdles include difficulty producing the multi-stranded circuit components *in situ* and poor understanding of design principles in complex cellular environments. Here we develop genetically encoded toehold exchange riboregulators, which harness TMSE to regulate protein expression, to successfully demonstrate *in situ* production and operation of TMSE circuits in living cells. We find the design principles and parts used for TMSE circuits in test tubes largely transfer to *Escherichia coli*, allowing construction of single-nucleotide-specific sensors, multi-layer cascades, and classifier circuits with programmable logic regimes, all with a single gate design derived from state-of-the-art *in vitro* TMSE components. Further, we illustrate how toehold exchange riboregulators can leverage TMSE to reprogram mRNA 5’ UTRs in *trans*, enabling predictable and modular control of translation strength over two orders of magnitude for diverse proteins without changing mRNA sequences. Our results identify optimal architectures for TMSE circuits in cells and caveats where strand exchange in cells differ substantially from cell-free systems, implicating active cellular processes. This study establishes toehold exchange riboregulators as versatile RNA devices for controlling protein expression and genetically encoded TMSE circuits as a unified platform for molecular programming across environments – bringing the full potential of nucleic acid computing from the test tube to the cell and accelerating advances in synthetic biology, nanobiotechnology, and cellular engineering.

## INTRODUCTION

Programming cellular behavior with the same predictability and flexibility with which we program electronic devices is a major goal of synthetic biology. Such a capability has enormous potential to advance biomanufacturing^1^, agriculture^2^, and medicine^3,4^. Tremendous progress in cellular programming has been made, from automated design of transcription factor-based circuits in bacteria^5,6^ to multi-strain yeast consortia that process information^7^ to programmed T cells that specifically recognize cancerous cells^8^. However, many challenges remain. Engineered genetic circuits remain relatively small (5 to 10 species) with limited processing power, in part due to crosstalk between components and the associated cellular burden of producing and operating the circuits^9^. Further, many biomolecular implementations of genetic circuits are well-suited for specific chassis but lack interoperability, leaving the field without a general molecular programming language that can span a wide range of applications. RNA synthetic biology and RNA nanotechnology have begun to address these challenges by programming cells with RNA:RNA interactions that have a small genetic footprint and are straightforward to design with universal base pairing rules^10–12^. Genetically encoded RNA circuits often rely on riboregulators – RNA devices that regulate gene expression in response to environmental triggers. Currently, engineered riboregulators are typically designed as RNA hairpins with an unstructured, single-stranded domain that serves as a toehold to recruit partially complementary single-stranded RNA (ssRNA) inputs that initiate a process of strand displacement, opening the hairpin to regulate gene expression at the transcriptional^13–15^ or translational level^15–18^ or through guide RNA activation in CRISPR/Cas systems^19–22^. Among other functions, these systems have been programmed to execute multi-input logic^22–24^ and multi-input/multi-output mapping^14,25^ in bacteria. However, scaling up information processing capacity of hairpin-based systems requires increasingly complicated RNA structures^14,23,24^, and distinct designs are often needed for each capability. For example, different riboregulator designs are used for translational activation and repression^16,17^, AND vs OR logic^14,23^, and detection of single nucleotide mismatches^26,27^. Although inputs and outputs of each device are modular, designs differ across devices making integration challenging. Further, logical operations often rely on cell-specific machinery, rather than on base pairing alone^23,28^, which makes porting technologies across environments difficult. A more composable RNA device design that processes information exclusively with base pairing would expand the capabilities of RNA circuits in cells.

Reactions based on toehold mediated strand exchange (TMSE) of nucleic acids, originally developed for dynamic DNA nanotechnology^29^, provide precisely this kind of scalable and versatile platform for building molecular circuits that interface with biological systems^30–32^. In contrast to conventional, hairpin-based systems, TMSE circuits rely on partially double-stranded nucleic acid complexes (‘gates’) composed of at least two strands. Single-stranded nucleic acid inputs react with sequence complementary gates *via* an unpaired toehold domain, releasing single-stranded nucleic acid outputs that in turn initiate downstream strand exchange reactions^29,33^. A key feature of TMSE reactions is toehold exchange, in which the formation of base-pairs on the input toehold on the gate is exchanged for the dissociation of base-pairs on the output toehold when the output strand is released (Figure 1a). Toehold exchange can be designed such that there is no net change in base pairing, making TMSE reactions readily reversible. This thermodynamic balance enforces strict sequence specificity and enables minor changes to toehold lengths to precisely control the kinetics and extent of reactions^34,35^ – capabilities difficult to achieve in hairpin-based systems without careful redesign. Toehold exchange also imparts the modularity and composability of TMSE gates, as the input and output portions of the gate do not impose any sequence constraints on each other. *In vitro* TMSE circuits composed of DNA have demonstrated some of the most sophisticated cell-free molecular computations to date^36,37^, including complex digital calculations^38^ and neural networks^39–42^ with over 100 components.

**Figure 1:**
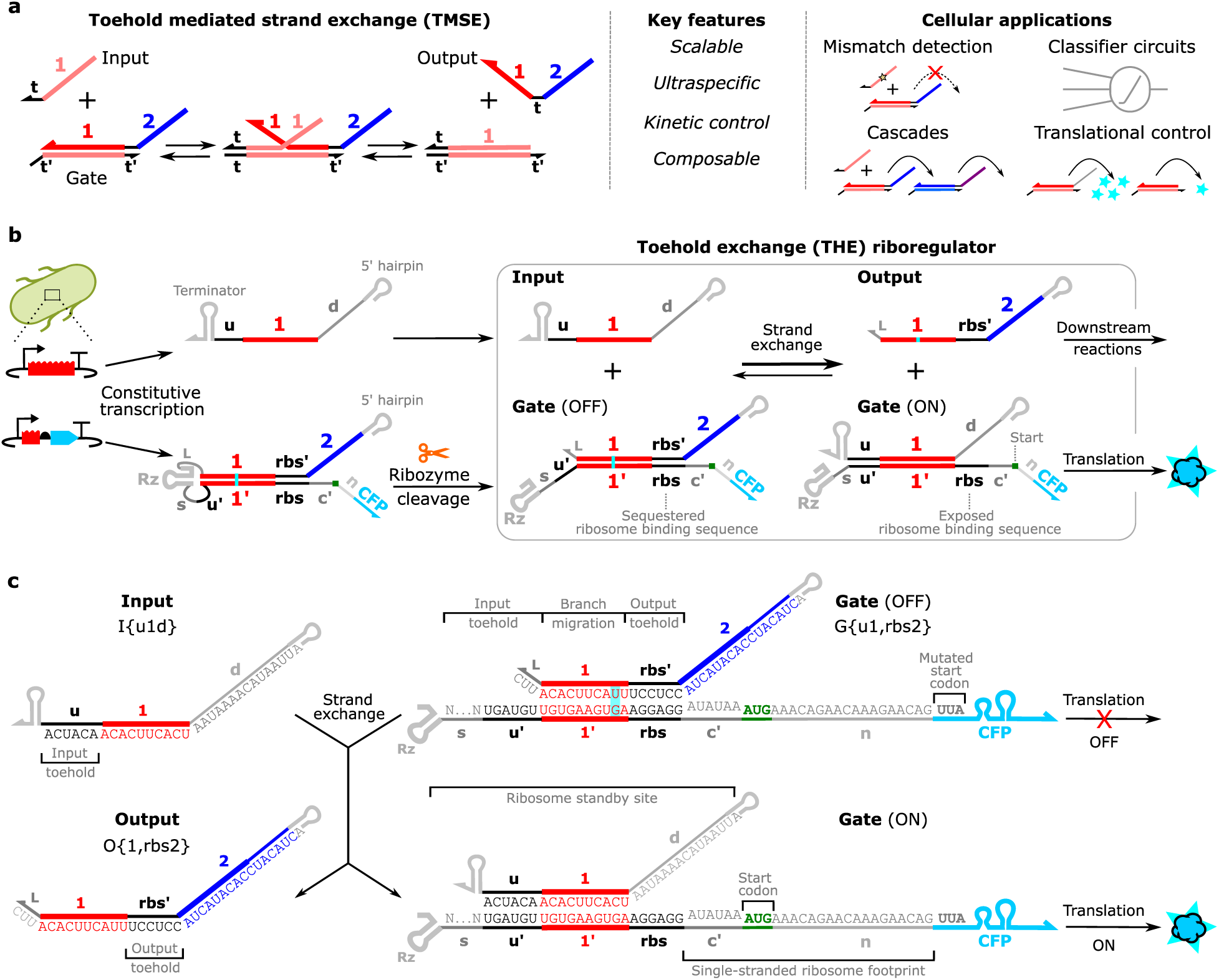
Toehold exchange (THE) riboregulator design. (**a**) Overview of TMSE reactions, features, and applications. (**b**) Schematic of toehold exchange riboregulator production and operation in bacteria. THE riboregulators are designed to couple RNA strand exchange to protein expression (Supplementary Figure 2). Single-strand RNA (ssRNA) inputs and gate hairpins are transcribed from DNA templates. An internal self-cleaving ribozyme (*Rz*) cleaves the gate hairpin to produce a double-stranded (dsRNA) gate complex. The stem of the dsRNA gate contains a sequestered ribosome binding sequence (*rbs*), preventing translation of a downstream protein coding sequence (in this case, cyan fluorescent protein, *CFP*). If an RNA input with complementary toehold (*u*) and branch migration (*1*) domains is present, the THE riboregulator and input undergo TMSE to displace an output strand from the gate and expose the *rbs* domain, initiating translation of the protein coding sequence. This process is referred to as toehold exchange because the base pairing of the input toehold (*u:ú*) is exchanged for the dissociation of the output toehold (*rbs′:rbs*). (**c**) Sequence schematic of an example toehold exchange riboregulator gate and input. The of *rbs*, *d* domain of the ribosome standby site, and the *ć* and *n* domains of single-stranded ribosome footprint of the Gate (ON) complex were designed to maximize translation rate (Supplementary Section 2.1). The *s* domain is shown as Ns as this variable-length domain can serve as a single-stranded spacer or an extended input toehold domain depending on the sequence and the input design. Inputs, gates, and outputs are defined by I{}, G{}, and O{} respectively, with the identities of their input and output subdomains specified within the braces separated by a comma. This nomenclature defines connectivity of components in larger networks. Here, and in subsequent figures, vertical cyan lines in component schematics designate G:U wobble pairings and the generic Rz designation is replaced by Rx, where x indicates ribozyme sequence identity. A detailed description of component nomenclature and domain identities is presented in Supplementary Section 1.

Despite their success in the test tube, the application of TMSE circuits to cellular engineering remains severely limited. While TMSE circuits have been transfected into cultured mammalian cells and used to process information and regulate gene expression^43,44^, the components of these circuits have a limited lifetime in the cellular environment. Moreover, in fast-growing organisms like bacteria, TMSE components quickly dilute out after only a few rounds of cell division (Supplementary Figure 2), rendering operation and measurement of circuit performance impossible. Ideally, TMSE circuits would be genetically encoded such that the constituent components are continuously expressed in cells and circuit instructions replicate across generations, allowing circuits to be continuously replenished to overcome degradation and dilution.

A promising approach for genetically encoding TMSE circuits involves transcribing RNA gates that initially fold into hairpins before being cut *via* internal ribozymes to produce dsRNA gates suitable for strand exchange^45,46^. This method allows multiple partially complementary dsRNA gate complexes to be produced together *in situ* without substantial cross-reaction. In *in vitro* transcription (IVT) reactions these cotranscriptionally encoded RNA strand displacement (ctRSD) circuits maintain the scalability, composability, and predictable and programmable kinetic behavior of DNA-based TMSE systems^45–47^.

Here we develop toehold exchange (THE) riboregulators to demonstrate genetically encoded RNA TMSE circuits in bacteria. THE riboregulators are designed analogously to ctRSD gates, with the process of TMSE exposing a previously occluded ribosome binding site (*rbs*) and permitting translation of a downstream protein coding sequence (Figure 1b). By coupling RNA strand exchange to fluorescent protein expression, we characterize the design parameters of genetically encoded RNA strand exchange circuits in *E. coli*. We find that most designs from a previously characterized *in vitro* ctRSD toolkit^47^ can be directly reused, highlighting their potential for cross-environment operation. However, we also identify a few exceptions where strand exchange trends in cells differ substantially from IVT^45^ and cell-free settings^48^. Using a suite of previously characterized domains, we demonstrate the modularity and specificity of input/output sequences as well as the composability of strand exchange gates by implementing multi-layer cascades.. Additionally, we highlight the versatile capabilities of genetically encoded RNA strand exchange reactions by building a 3-input classifier circuit that can be programmed to execute multiple distinct logic operations. We further demonstrate that THE riboregulators can detect single-nucleotide differences and dynamically reprogram 5ʹ untranslated regions (UTRs) to precisely and predictably tune translation rates of disparate proteins. These results illustrate how a single THE riboregulator design can achieve functions that required over 5 unique designs with hairpin-based systems^23–27,49^. More broadly, this work represents the first demonstration of genetically encoding TMSE circuits in living cells, laying the foundation for implementing existing sophisticated TMSE circuits for applications in cellular engineering and intracellular control of nucleic acid nanostructures^50^, condensates^51^, and nanomachines^52^.

## RESULTS AND DISCUSION

### Toehold exchange riboregulator design

In simple buffered solutions, the medium in which the vast majority of TMSE research has been conducted, strand exchange is measured with molecular beacons – chemically synthesized, fluorescently labeled nucleic acid complexes^45,48^. However, using molecular beacons in bacteria is impracticable, as cell division rapidly dilutes their concentration below levels of detection. To overcome this limitation, we began by developing a genetically encoded reporting scheme to measure TMSE in bacterial cells, allowing the reporter to be maintained in DNA and replicated during cell division. Genetically encoded RNA reporters used in cells are typically linked to either fluorescence-activating RNA aptamers^53–55^ or to riboregulators that control the expression of a fluorescent protein^13,17^. Rather than connect the output of a TMSE reaction to existing riboregulators for reporting^13,17,53,55^, we sought to develop a measurement that relied directly on dissociation of the output toehold of a gate as this explicitly indicates completion of toehold exchange. We focused on developing a TMSE-based riboregulator so that protein expression could readily connect RNA strand exchange reactions to cellular function.

To connect the process of toehold exchange to regulation of protein expression, we used the previously described ctRSD gates and inputs as a model^45,47^. To modify the gates to control protein expression, we changed the output toehold of the gate to act as a ribosome binding site (*rbs*), *i.e.*, a sequence crucial for initiating translation that is complementary to the *E. coli* 16S ribosomal rRNA (also referred to as the Shine-Dalgarno sequence)^56,57^. Downstream of this *rbs* toehold, we placed a start codon and a coding sequence for cyan fluorescent protein (CFP) so that translation of a fluorescent protein would initiate when the *rbs* is exposed after toehold exchange (Figure 1b, Supplementary Figure 2). In the absence of a complementary input RNA, the *rbs* is sequestered in the stem of the dsRNA gate, preventing translation initiation. In addition, we extended the input RNA sequence to include a 5′ single-stranded domain, termed *d*, that we hypothesized would serve as a distal ribosome standby site to increase translation initiation rates of the otherwise highly structured ON gate^58,59^ (Figure 2a, Supplementary Section 2.1).

**Figure 2:**
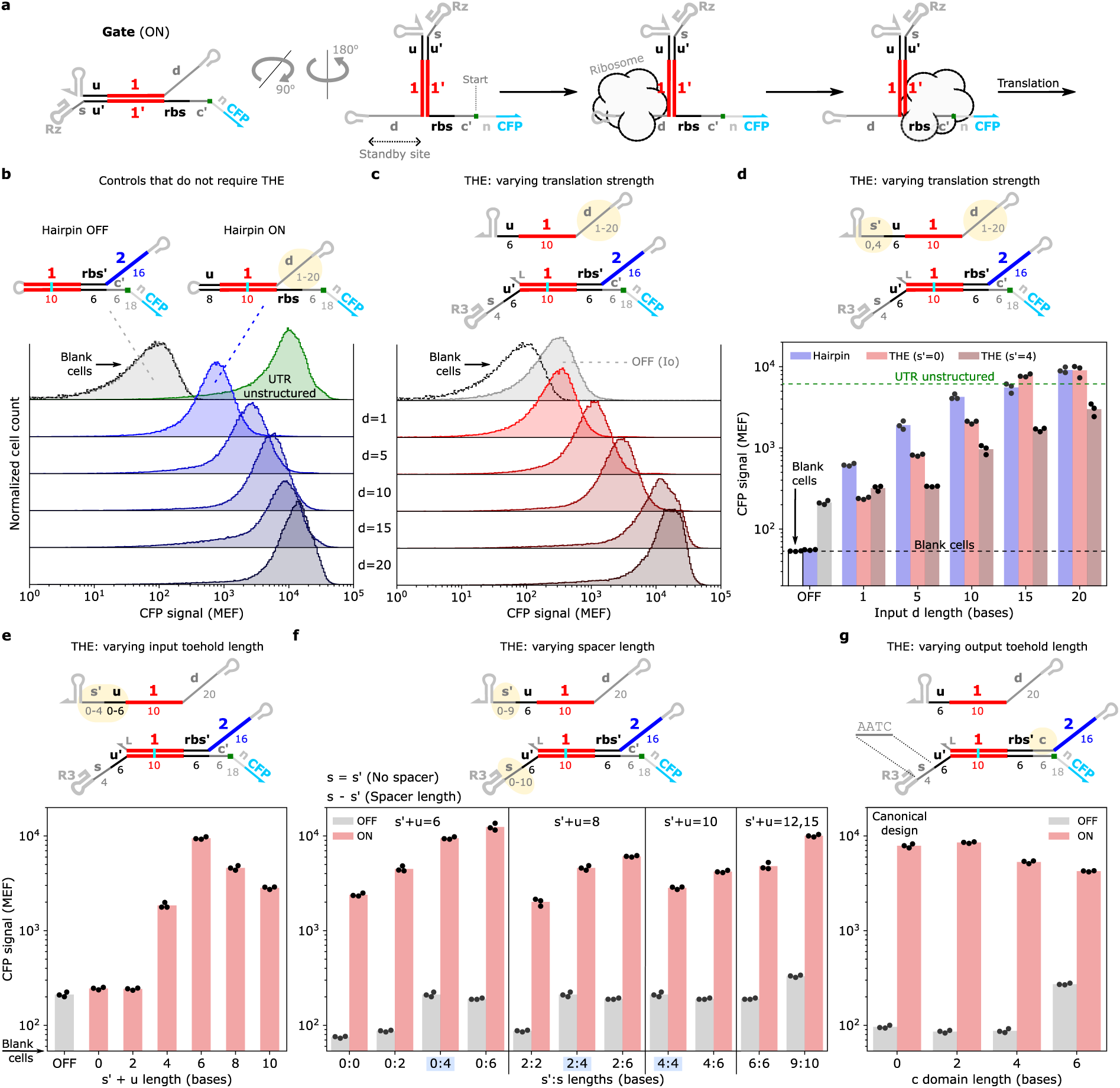
THE riboregulator design space in *E. coli*. (**a**) A single-stranded domain distal to the dsRNA stem of an ON THE riboregulator (*d* domain) influences translation rate by providing a standby site for ribosome binding prior to translation initiation^58,59^. (**b**,**c**) Flow cytometry results of THE riboregulators with different *d* domain lengths for hairpin control components that do not require strand exchange for translation (panel b) or separate input and gate components that require strand exchange for translation (panel c). The green ‘UTR unstructured’ distribution represents a construct lacking secondary structure in the 5ʹ UTR (Supplementary Figure 1). (**d**) Geometric means of flow cytometry distributions in panels b and c, and from experiments with an input possessing a 10-base toehold (*s*′=4). (**e**,**f**,**g**) Geometric means of flow cytometry distributions characterizing different input toehold lengths (panel d), input toehold and spacer (*s*) lengths (panel e), and increasing output toehold lengths *via* a clamping (*c*) domain adjacent to the *rbs* (panel f). The results highlighted in blue on the x-axis of panel f are also shown in panels c and d. The ‘canonical design’ in (panel g) represents the gate design used in rest of the study. In all panels, yellow-highlighted domains on the RNA schematics represent the domains varied in the accompanying results. The numbers below the constructs indicate domain lengths in bases. MEF indicates molecules of equivalent fluorophore. Results labeled ‘OFF’ represent gates coexpressed with a non-complementary input (Io). Replicates represent three independent colonies selected after transformation. See Supplementary Figure 5 for plasmid schematics. See Supplementary Figures 18 and 20 for flow cytometry distributions of results in panels (e-g).

To test the feasibility of our design, we started with simplified hairpins that mimic the OFF and ON states of the gates, bypassing the complexity of ribozyme cleavage and strand exchange (Figure 2b). We cloned these hairpin controls, under control of a constitutive T7 promoter, into plasmids and transformed them into *E. coli* BL21 Star (DE3), which expresses T7 RNA polymerase (RNAP) upon induction with IPTG. Expression of a constitutively OFF hairpin, containing a sequestered *rbs* within its stem, produced minimal signal that was indistinguishable from autofluorescence (Blank cells, Figure 2b). In contrast, expression of a constitutively ON hairpin, which presented an exposed *rbs* and varying *d* domain lengths, produced measurable CFP expression above background. As hypothesized, increasing the length of the *d* domain increased CFP expression, with *d* ≥ 15 bases producing similar CFP signal to a control mRNA with an unstructured 5ʹ UTR (Figure 2b). These results demonstrate that sequestering an *rbs* at the base of a hairpin strongly represses translation and that a long *d* domain enables high translation levels despite the substantial structure near the exposed *rbs*.

We next tested the RNA strand exchange scheme, which requires successful gate folding, ribozyme cleavage, and toehold exchange to initiate translation. We term these gates toehold exchange (THE) riboregulators (Figure 1c). In these experiments the gate and input RNAs, both under control of constitutive T7 promoters, were cloned into separate plasmids that were then co-transformed into *E. coli* BL21 Star (DE3) (Supplementary Section 3). Both the input and riboregulator RNAs were designed with a 6-base input toehold (*u* and *u′*, respectively). The riboregulator was designed with a 4-base spacer (*s*) domain and we tested input RNAs with *d* domains spanning 1 base to 20 bases (Figure 2d). As with the ON hairpin control, increasing *d* domain length on the input RNA resulted in increased CFP expression. Additionally, as negative control, we tested an input RNA designed to be non-complimentary to the THE riboregulator (*Io*, OFF). In contrast to the OFF hairpin control, the THE riboregulator coexpressed with this non-complementary input produced CFP expression that — while still over an order of magnitude below the ON state — was above autofluorescence, indicating leaky expression (Figure 2c,d). For input RNAs with *d* ≥ 15 bases and only 6 bases of complementary with the gate input toehold (*s*′ = 0), protein expression was not limited by RNA strand exchange, as both the ON hairpin and the ON THE riboregulators produced similar signal.

To verify that the THE riboregulator operates *via* the designed ctRSD^45^ mechanism, we tested the ability of inputs to activate gates with either non-complementary toeholds, branch migration domains, or both. We found that if either the toehold (*u:uʹ*) or the branch migration domain (*1:1ʹ*) sequences of the input and THE riboregulator were not complementary, protein expression was not activated, confirming that strand exchange was sequence specific (Supplementary Figure 17). We next tested THE riboregulators with a point mutation in the ribozyme that abolishes cleavage, confirming that ribozyme cleavage was necessary for high protein expression, particularly for input RNAs with 6-base toeholds (Supplementary Figure 17). RT-qPCR measurements of ribozyme cleavage^60^ further indicated > 90 % of the gate transcripts cut in the cellular environment (Supplementary Section 5). Lastly, in a companion study in cell-free expression systems^48^, we found the fluorescent protein measurements from THE riboregulators recapitulated the same trends observed for an exogenously added molecular beacon that reports on the release of the output strand, indicating THE riboregulators provide a valid indicator of RNA strand exchange.

### Characterizing THE riboregulator design space

We next explored how THE riboregulator design features influenced protein expression, focusing on design alterations commonly used to control the rate and extent of TMSE in enzyme-free conditions^35^ and IVT^45^. We began by investigating the influence of input toehold complementarity length by testing a THE riboregulator with a combined spacer and input toehold (*s*+*u′*) length of 10 bases with input RNAs that had 0 bases to 10 bases of complementarity with this region (*s′+u*). We found that the input RNA needed at least 4 bases of complementarity with the gate in this single-stranded region to produce protein expression above the OFF gate (Figure 2e). Surprisingly, increasing *s*+*u′* complementarity beyond 6 bases resulted in decreased expression levels (Figure 2e and Supplementary Figure 18). Input RNAs with 10-base *s*+*u′* had lower CFP expression than input RNAs with 6-base *s*+*u′* complementarity across nearly all *d* domain lengths (Figure 2d). This phenomenon was not sequence dependent as the same results were observed for different branch migration domains, toeholds, ribozymes (Supplementary Figure 19), and input terminators (Supplementary Figure 40). These results contrast with experiments in cell-free expression systems, in which increasing s+*u′* complementarity increased both strand exchange rate and the level of protein expression^48^.

Intrigued by the unexpected trend in protein expression as a function of input RNA toehold length, we next explored how changing the spacer (*s*) length in the THE riboregulator influenced expression levels. We tested THE riboregulators with spacer domains spanning 0 bases to 10 bases along with input RNAs possessing increasing degrees of complementarity with the *s* domains, *i.e.,* increasing *s′* lengths (Figure 2f). In line with IVT experiments^45^, increasing *s* length resulted in higher protein expression across all RNA inputs. Input RNAs with 6-base toeholds (*s′* = 0) showed consistently higher expression than input RNAs with more complementarity (*s′* > 0) across *s* lengths (Figure 2f). Higher protein expression with increasing *s* length is likely due to an increase in the RNA strand exchange rate, which was independently measured in a reconstituted cell-free protein expression system^48^. Increasing *s*+*u′* complementarity of the input RNA to 15 bases brought protein expression up to similar levels seen for a 6-base toehold (*s′* = 0) (Figure 2f), suggesting that long enough toeholds (*s′*+*u:s*+*u′)* can mitigate whatever cellular interactions interfere with strand exchange at intermediate toehold lengths.

Altering gate spacer lengths and input toehold complementarity provided fine-tuned control of protein expression levels, so we next explored the influence of output toehold length. Specifically, we extended the length of the gate duplex between the *1:1ʹ* and *rbs:rbs′* domains by up to 6 bases (*e* domain, Supplementary Figures 20 and 21). Surprisingly, input RNAs with 6 bases of complementarity to the input toehold of the gate (*u:u′*) were able to trigger nearly the same level of protein expression when either 6 bases (*rbs:rbs′*) or 12 bases (*e+rbs′:e′+rbs*) needed to dissociate on the output toehold of the gate, which should not be possible with purely base pairing thermodynamics^35^. These results manifested for multiple input domain sequences and appeared to be unique to RNA strand exchange in the cellular environment, as increasing the length of the output toehold relative to the input toehold proportionally lowered protein expression and RNA strand exchange rates in a reconstituted cell-free protein expression system^48^. Similar results were observed when increasing the output toehold length by adding a clamp domain (*c*) domain downstream of *rbs′*, which we hypothesized would reduce leak by burying the *rbs* deeper in a dsRNA stem (Supplementary Section 2.1). Including *c* domains of up to 6 bases did not reduce leaky expression in the OFF state and only mildly decreased protein expression in the ON state using an input RNA with a 6-base toehold (Figure 2g and Supplementary Figure 20).

Based on the results of this characterization, we selected a THE riboregulator design with an input toehold length of 6 bases, an *s* domain of 4 bases, an output *rbs* of 6 bases, and no *c* domain as the canonical design to move forward with for further characterization and integration into TMSE circuits (Figure 1c). Because the RNA input with a 6-base toehold showed the highest expression, an *s* domain that could serve as a longer input toehold on the gate was not needed. So, we changed the *s* sequence to have low complementarity with input and output sequences of the ctRSD toolkit. This minor change also reduced leak expression from the OFF gate (Figure 2g).

### Characterizing modularity and specificity of THE riboregulators

Using the canonical THE riboregulator design, we next investigated how robust performance was to swapping in different sequence domains from the IVT-characterized ctRSD toolkit^47^. The ctRSD toolkit contains a suite of largely modular gate domains required to build multi-component circuits with more sophisticated behaviors. For example, using different input/output toehold sequences within a gate mitigates misfolding^47^, diverse ribozyme sequences are necessary to reduce homology across circuits with multiple gates, and orthogonal input/output sequences are needed to connect components together.

To assess how well the modularity of the ctRSD toolkit demonstrated in IVT transferred to *E. coli*, we tested THE riboregulators with two different input toehold sequences (*u* and *v*) and four distinct ribozyme sequences^47^. THE riboregulators with *u* or *v* input toeholds resulted in ON/OFF ratios > 20 for most of the tested ribozymes (Supplementary Figure 22). One exception was the *CPEB-3* ribozyme^61^ (*Rh*), which resulted in low signal when used with the *v* toehold, likely due to poor cleavage in this context (Supplementary Section 5). We also tested the performance and specificity of THE riboregulators with 6 different input branch migration domains that had demonstrated similar performance in IVT^47^ (Figure 3a). All inputs produced similar ON expression levels and ON/OFF ratios > 20 (Figure 3b). The inputs were also orthogonal (Figure 3c and Supplementary Figure 23), despite having the same toehold sequence. Together, these results show that ctRSD domains initially characterized *in vitro* largely act as modular units when used in cells, supporting development of more complex circuits.

**Figure 3:**
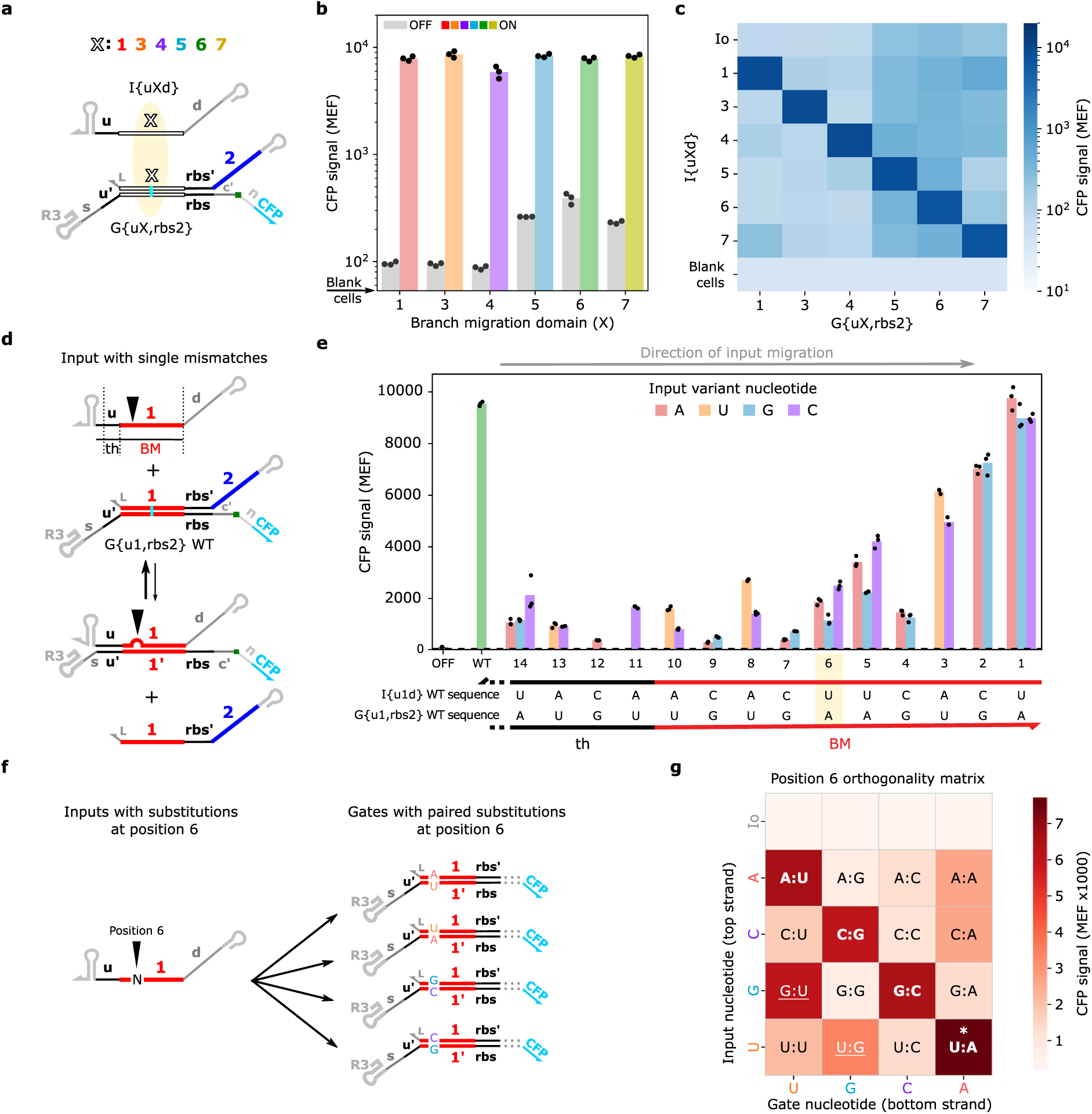
Specificity of THE riboregulators. (**a**) Schematic of THE riboregulators with different input branch migration (BM) domains, denoted by *X*. The branch migration sequences (*1*, *3*, *4*, *5*, *6*, *7*) were taken from the IVT-characterized ctRSD toolkit^47^. (**b**,**c**) Geometric means of flow cytometry distributions (panel b) and orthogonality matrix (panel c) for THE riboregulators with the different branch migration domains (see Supplementary Figure 23 for flow cytometry distributions). (**d**,**e**) Schematic (panel d) and flow cytometry results (panel e) of I{u1d} substitution variants that introduce a single mismatch at different positions along the input migration path, which starts with toehold (*th*) binding on the left and proceeds to the right through the branch migration (BM) domain. Substitutions resulting in G:U wobble pairs between the input and gate were avoided, as these are expected to promote efficient strand exchange. Data points represent three independent colonies, and bars represent their mean. (**f**,**g**) Schematic (panel f) and flow cytometry results (panel g) for single nucleotide mismatch classification by leveraging the specificity of THE riboregulators. The heatmap shows the mean CFP expression from three independent colonies. Complementary base pairs trigger CFP expression along the diagonal of the heatmap. The underlined G:U and U:G wobble pairs also exhibit appreciable CFP expression, as expected (see Supplementary Figures 25 and 26 for additional details and measurements at different substitution positions). The * indicates the WT sequence.

When testing the orthogonality of different inputs, we observed that inputs I{u1d} and I{u7d} were orthogonal despite only differing by only 3 bases, suggesting TMSE could be very specific in *E. coli*. To test the limits of strand exchange specificity, we evaluated a collection of 10 I{u1d} variants with two consecutive substitutions (Supplementary Figure 24) and a collection of 30 I{u1d} variants with only a single substitution (Figure 3d) in either the toehold (*u*) or branch migration domain (*1*) against G{u1,rbs2}. The latter collection included multiple different substitutions at most positions to assess nucleotide-specific effects. Across the single substitution inputs, mismatches introduced near the 3ʹ end of the input – in either the toehold domain or first half of branch migration domain – reduced CFP expression > 3-fold. In contrast, single mismatches introduced to the 5ʹ end of the input (in the last three bases of the dsRNA branch migration region of the gate) were well-tolerated and produced a signal within a factor of two of ‘wild type’ I{u1d} (Figure 3e). This trend was qualitatively consistent with the results from the doubly substituted inputs (Supplementary Figure 24). Additionally, these findings align with theoretical and experimental studies of DNA TMSE in the presence of mismatches in simple buffered solutions^62–64^, suggesting that *in vitro* characterizations of strand exchange remain applicable in cellular contexts.

We subsequently built 19 G{u1,rbs2} variants with compensatory substitutions that restored canonical base-pairing with a substituted input. When co-expressed, 17 of the 19 variant input-gate pairs exhibited activity levels comparable to the wild type pair (Supplementary Figure 26). The gates for the remaining two pairs had substitutions in their input toehold sequence and were predicted to misfold (Supplementary Figure 27), suggesting that their failure to activate is a consequence of gate malfunction rather than an effect of the input. Together, these experiments highlight that THE riboregulators possess exquisite sequence specificity likely arising from altered strand exchange kinetics and completion probabilities produced by even a single mismatch at sensitive positions.

The sharp drop in activity resulting from single mismatches near the 3ʹ end of THE riboregulators suggested that a single mismatch may be sufficient to prevent crosstalk between ctRSD circuit components. To determine whether single-base differences enable mutual orthogonality, we constructed a complete combinatorial cross-reactivity screen at positions sensitive to mismatches. Focusing on positions 5 and 6 in the branch migration domain, we tested all 16 combinations of input and gate single nucleotide variants at both positions (Figure 3f and Supplementary Figures 25 and 26). As expected, fully complementary input and gate combinations produced strong CFP expression signal, and all combinations with a single mismatch had lower CFP expression (Figure 3g and Supplementary Figures 25 and 26). The two combinations off the diagonal with appreciable CFP expression represented wobble base pairs (G:U and U:G) and their relative expression levels correlated with predictions from the nearest neighbor thermodynamic model^65^ (Supplementary Figure 26). These results suggest that a single mismatch can confer mutual orthogonality, making the potential sequence space for cellular ctRSD circuits enormous. There are > 1 million sequences that differ by at least one nucleotide across input positions 6 to 14 if all four bases are used and nearly 60 000 sequences if the three-base restriction for input sequences is maintained.

### Testing composability of genetically encoded TMSE gates in bacteria

Having characterized a suite of orthogonal input, output, and ribozyme sequences with individual THE riboregulators, we next explored composing multiple genetically encoded TMSE gates – designed identically to ctRSD gates characterized by IVT^45,47^ – into multi-layer cascades using a THE riboregulator as the reporter. Signaling cascades are essential for coordinating timing of molecular events^66^, executing logical operations^38^, and multi-input classification^39^. However, cascades that rely entirely on RNA to propagate signal across multiple layers are not well characterized in bacteria. Signal decay between layers or cross-reaction between multiple partially complementary, coexpressed gates could limit such RNA-only cascades in cells.

Starting with 2-layer cascades, we designed upstream gates with output domains that trigger a downstream THE riboregulator (Figure 4a). We designed upstream gates for three different input RNAs (I{v3}, I{v4}, I{v5}). As in previous IVT studies^47^, we designed cascades with input and output toeholds alternating between *u* and *v* across circuit layers to avoid repeating toeholds within any gate. Additionally, we tested two variations of each upstream gate, one variation with the canonical 3-base linker (*L*) between the output strand and the ribozyme – as previously demonstrated in IVT^45,47^ – and another variation with a 3′ hairpin^67^ in place of this short linker. Upstream gates without the 3′ hairpin on their output strand produced negligible signal while those with the 3′ hairpin produced ON/OFF ratios around 5 (Figure 4b). These results were not sequence specific, as we found outputs with 3′ hairpins were necessary for propagating signal for upstream gates with a different ribozyme sequence (Supplementary Figure 28) and a different output sequence (Supplementary Figure 29). Further, we confirmed four additional RNA stabilizing hairpins^67,68^ could be used with similar performance (Supplementary Figure 28). The 3′ hairpin may protect the output RNA from rapid degradation after it is released, giving it enough time to initiate a downstream reaction. However, the same 3′ hairpin dependence was observed for cascades in a reconstituted cell-free expression system advertised to have minimal nuclease activity^48^, suggesting more than stability alone could be at play.

**Figure 4:**
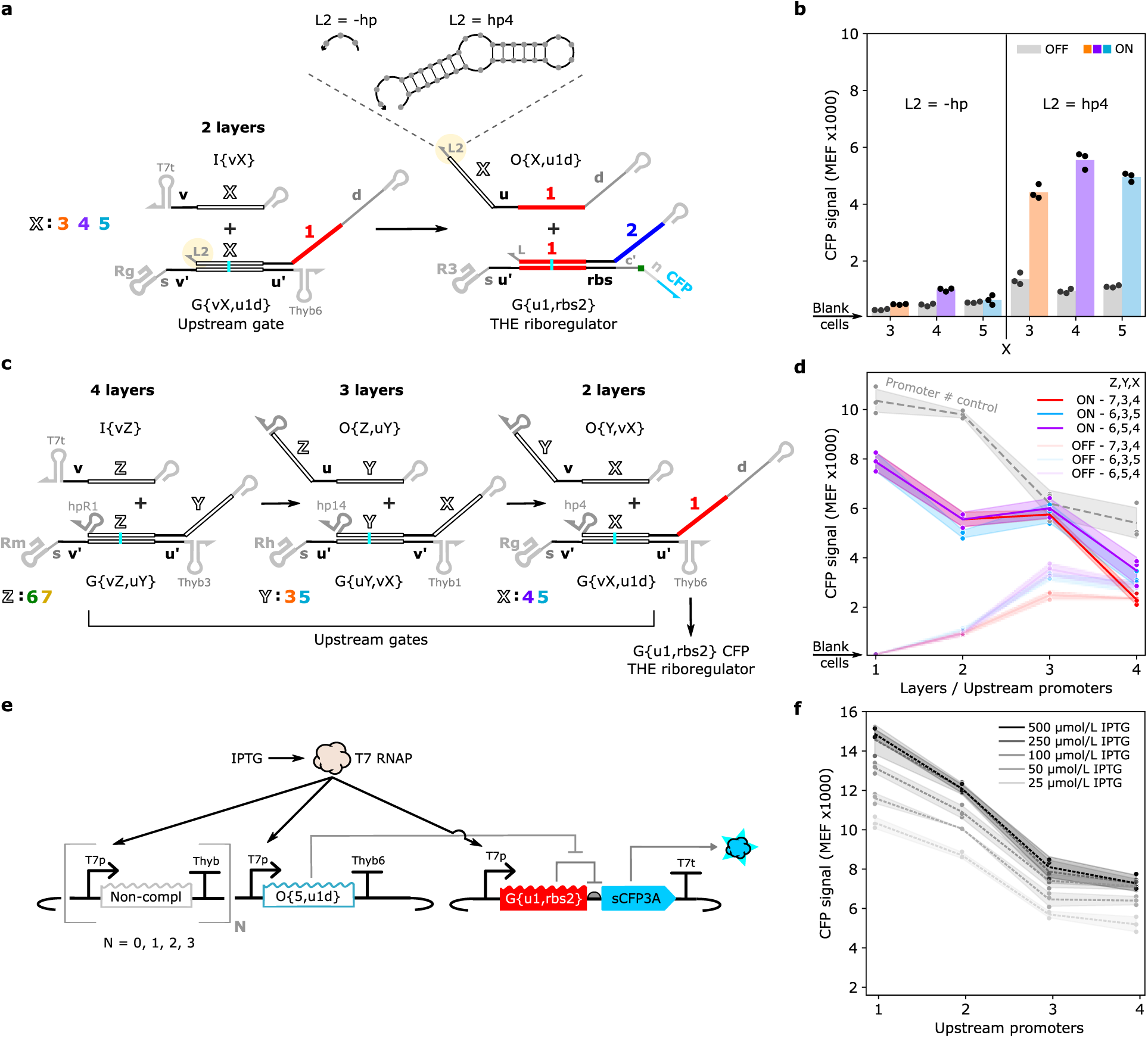
Multi-layer cascades of genetically encoded TMSE gates. (**a**,**b**) RNA circuit schematic (panel a) and flow cytometry results (panel b) of 2-layer RNA strand exchange cascades. Three different input branch migration domains were tested (*X*), each with a short 3-base linker (*L*=-hp) or a 37-base hairpin forming linker (*L*=hp4) at the 3′ end of the output strand. (**c**,**d**) RNA circuit schematic (panel c) and flow cytometry results (panel d) of multi-layer RNA strand exchange cascades with different combinations of input and output domains (*X*,*Y*,*Z*). For 2- and 3-layer cascades, upstream gates were not encoded and instead the complementary input to the gate in the highest layer was expressed. See Supplementary Figure 6 for plasmid schematics. See Supplementary Figure 29 for multi-layer cascades with additional input and output combinations. The dashed gray line represents a single layer cascade with T7 promoters regulating expression of non-complementary (decoy) input RNAs in lieu of the gates used in the cascades (panel e). (**e**,**f**) Plasmid schematic (panel e) and flow cytometry results (panel f) of the promoter number controls tested at different IPTG concentrations (100 µmol/L is used in most experiments). See Supplementary Figure 7 for expanded plasmid schematics. Data points represent three independent colonies, solid lines and bars represent the mean, and shaded regions represent one standard deviation from the mean.

We next expanded to test four different sequence combinations of 3-layer cascades and six different sequence combinations of 4-layer cascades (Supplementary Figure 29). Because all upstream gates were encoded on the same plasmid (Supplementary Figure 6), each gate was designed with a different ribozyme sequence and a different 3′ output hairpin to reduce sequence homology (Figure 4c). Overall, increasing the number of layers resulted in lower ON signal and higher OFF signal, with ON and OFF becoming indistinguishable at 4 layers (Figure 4d). Similar to the 2-layer cascades, performance of the 3-layer cascades was consistent across different gate sequences (Figure 4d) and only 1 of the 8 upstream gates we tested was non-functional (Supplementary Figure 29). Decreasing ON signal with increasing layers cannot be attributed entirely to TMSE, as a single layer circuit that controlled for the total number of promoters in each cascade (Figure 4e) displayed decreasing signal as the number of promoters was increased (Figure 4f), likely a consequence of increasing resource competition as more circuit components are added. Increasing T7 RNAP expression did not change this trend (Supplementary Figure 30). The observed increase in OFF signal with increasing layers could be the result of misfolded gates that react without input as seen in IVT^45^ or a fraction of gates reacting cotranscriptionally. In an effort to improve the ON/OFF ratios of the multi-layer cascades, we included a fuel strand that recycles the input strand^38^ (Supplementary Figure 31) and tested gates with an inverted transcription order^47^ (Supplementary Figure 29) but neither strategy improved cascade performance.

### Cellular applications of genetically encoded TMSE circuits and THE riboregulators

Our characterization of genetically encoded TMSE components provides proof of principle for adopting this biotechnology in a variety of cellular applications. So, we next highlight how these design principles can support application development.

The specificity and composability of TMSE reactions support larger circuit development, but our multi-layer cascade results indicate larger circuits in bacteria should have a wide architecture, in which multiple inputs are processed in a single layer, rather than a deep architecture that processes inputs across many layers. To illustrate how such larger circuits can be built we implemented a thresholding-based multi-input classifier. This circuit architecture serves as a base unit in many TMSE-based logic circuits and neural networks^38,41^ and can classify different combinations of inputs in a single layer. The circuit relies on controlling the relative rates of reaction of inputs with a THE riboregulator and a threshold gate (TG{u1d}), such that inputs react with TG much faster than with the reporting THE riboregulator (Figure 5a). In this way, the circuit only produces an output if there is more input than TG (Figure 5b). When deployed as a logic element^38^, the same circuit can be reprogrammed to execute different operations. For example, a circuit with two inputs can exhibit OR logic if no, or very little, TG is produced, *i.e.,* the presence of either input is sufficient to produce an output. Alternatively, if TG is expressed at a level higher than either input alone, the system will exhibit AND logic because both inputs are required to produce an output. When deployed as a node in a neural network^41^, the circuit functions as a linear classifier such that the node activates when the sum of the inputs exceeds a set threshold.

**Figure 5:**
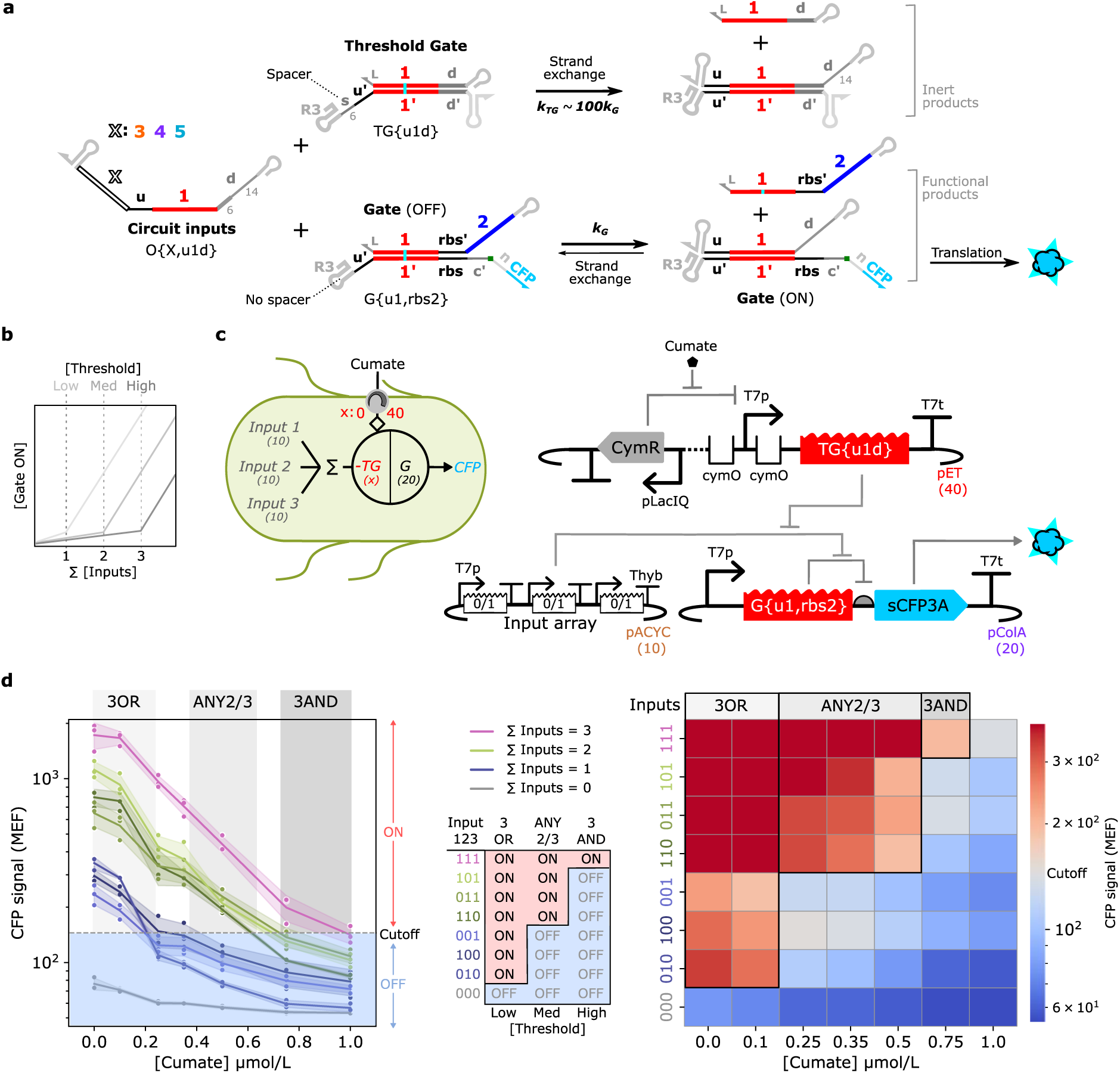
A field programmable, multi-input classifier circuit. (**a**) RNA strand exchange classifier implementation. Input RNAs with different 3′ trailing domains (*X*) mimic the outputs of upstream ctRSD gates. Inputs react with either a THE riboregulator to initiate protein translation or with a threshold gate (TG). TG is designed to react with the input faster than the THE riboregulator to sequester inputs before they can initiate protein expression. Substantial protein expression is achieved when the sum of input transcription is greater than TG transcription. A 6-base spacer (*s*) on TG compared to no spacer on the THE riboregulator provided the kinetic disparity between the two components. (**b**) For the circuit in (panel a), the concentration of TG dictates the concentration of input needed to activate the THE riboregulator. (**c**) The cell-level (left) and DNA-level (right) implementation of the classifier circuit. TG subtracts circuit inputs, and cumate concentration dictates TG transcription *via* an inducible promoter. For the eight input combinations, a 1 indicates expression of a complementary input and a 0 indicates expression of a non-complementary (decoy) input. Numbers in parentheses indicate nominal plasmid copy numbers, which bound relative expression of components. See Supplementary Figure 10 for detailed plasmid schematics. (**d**) Flow cytometry results of the classifier circuit across cumate concentrations, illustrating analog behavior (left) and different Boolean-like classification regimes (right). Data points represent three independent replicates, lines represent the mean, and shaded regions represent one standard deviation from the mean. The gray dashed line in the left plot indicates the ON vs OFF cutoff, which is the center value of the divergent heatmap. Controls confirming the circuit mechanism are in Supplementary Figure 32 and flow cytometry distributions are in Supplementary Figure 33.

To implement a multi-input classifier, we designed 3 different RNAs to serve as circuit inputs (O{3,u1d}, O{4,u1d}), and O{5,u1d}) that converge on a single layer with a THE riboregulator and TG. To design a suitable TG we used the principles uncovered in Figure 2, introducing a long gate spacer (*s*) domain to achieve a predicted ∼100-fold faster strand exchange rate (from Refs^45,48^ and Figure 2f) for TG with the inputs compared to the reporting riboregulator (Figure 5a). We used gate spacer length over increasing input toehold complementarity for kinetic control because we observed decreased protein expression for input RNAs with toeholds > 6 bases (Figure 2e). To control the expression level of TG, we placed it under control of an inducible CymR promoter (Figure 5c). Based on the nominal copy numbers of the plasmids we used, this circuit should be able to produce 3 different classifications depending on the concentration of cumate used: 3 input OR, ANY 2 OUT OF 3, and 3 input AND (Figure 5).

The THE riboregulator and TG plasmids were transformed with each of the 8 plasmids containing different circuit input combinations. Cells encoding each of these 8 combinations were then tested across a range of cumate concentrations to experimentally verify the 3 different classification regimes (Figure 5d). Additional control experiments were consistent with this circuit operating *via* the kinetic thresholding mechanism compared to resource competition or retroactivity (Supplementary Figure 32). These results illustrate a field-programmable^42^ RNA circuit design for executing multi-input information processing in a single layer. By varying the inducer concentration in the growth media, the same genetic constructs can be reprogrammed to exhibit different classifications on the same set of inputs, an ability that typically requires different circuit and input designs with existing riboregulators^14,23^.

Beyond circuits, the precise control of protein expression offered by THE riboregulators could support a wide range of biotechnology applications. A particularly unique and useful mechanism of control is changing the length of the *d* domain on short, non-coding RNA inputs to trigger translation-level changes in protein expression of an mRNA under control of a THE riboregulator. Current riboregulators control translation at the post-transcriptional level, using RNA triggers to activate protein expression at a fixed translation strength^14,17^. Tuning translation strength is typically achieved by changing the *rbs* of the mRNA, a hard-coded genetic change that can introduce CDS-specific base pairings that result in unexpected expression (Figure 6a, left). As the *rbs* and CDS are sequence constrained, redesigning to remove such an effect is challenging. With a THE riboregulator, translation strength can be precisely tuned without making genetic changes to the mRNA sequence, as strand exchange allows an mRNA’s 5ʹ UTR to be reprogrammed with standby sites of differing translation strengths – dictated by the *d* domain length on the exchanging input RNA^58^ (Figure 2a). The designed secondary structure of a THE riboregulator reduces the propensity for confounding base pairing interactions, which should result in predictable changes in expression across translation strengths (Figure 6a). If undesired interactions are introduced, the sequence of the *d* domain can be changed independently of the *rbs* or CDS to recover desired expression.

**Figure 6:**
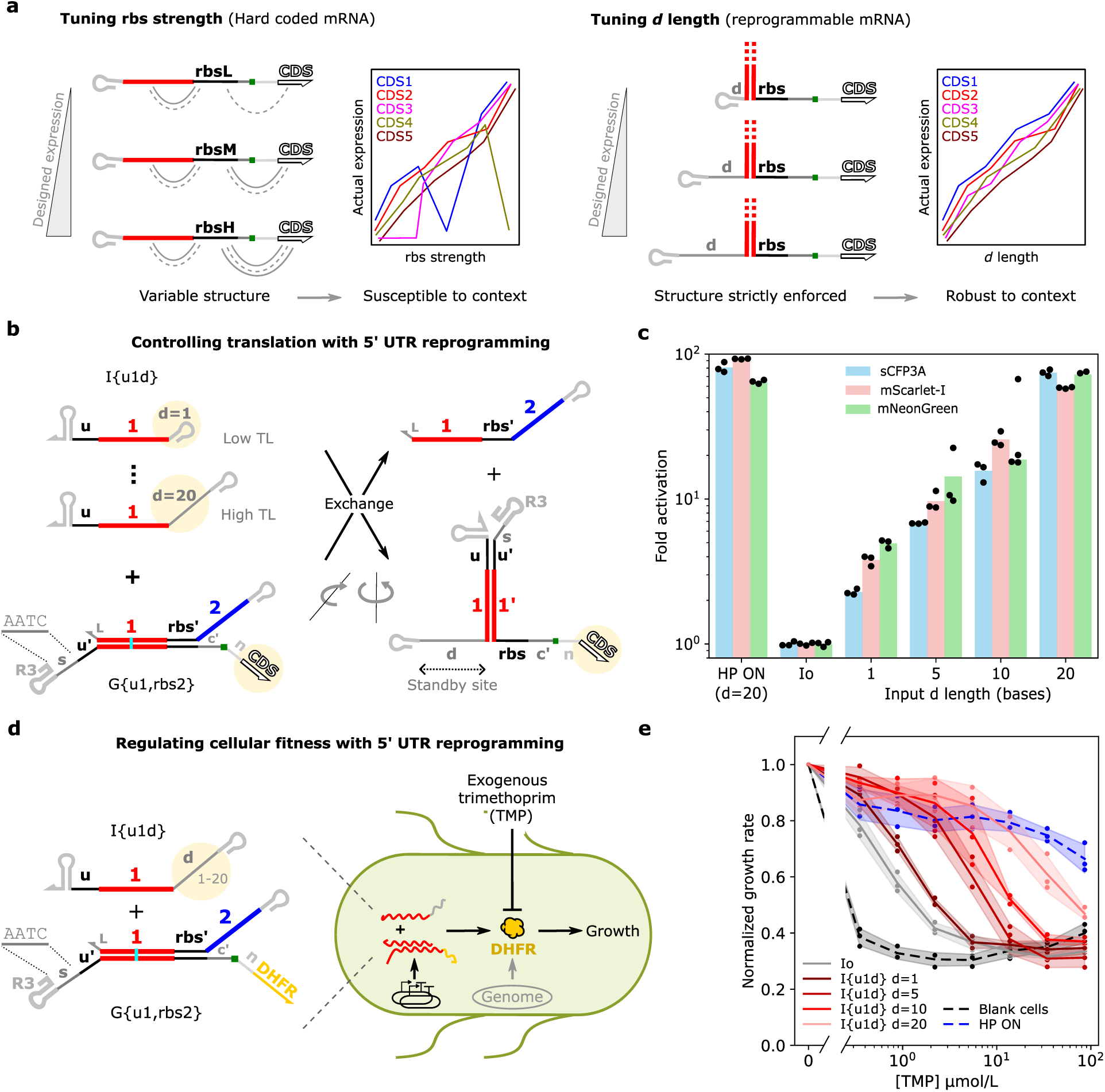
THE riboregulators enable precise translational control of proteins with different sequences and functions. (**a**) Left: Tuning mRNA translation strength by changing the ribosome binding sequence (*rbs*). Gray parabolas below the mRNAs indicate strong (solid lines) or weak (dashed lines) base pairing interactions, which can alter anticipated expression levels. Right: Changing *d* domain length on a THE riboregulator input RNA allows translation strength tuning independent of mRNA sequence. (**b**,**c**) Schematic (panel b) and flow cytometry results (panel c) for control of translation strength by leveraging strand exchange to reprogram the 5ʹ UTR of THE riboregulator mRNA. Bar plot shows the fold activation for three CDSs, with each data point representing expression from an independent colony normalized by the mean value of the decoy (Io) strain for that CDS. (**d**,**e**) Schematic (panel d) and plate reader data (panel e) for regulation of cellular fitness *via* control of *E. coli* dihydrofolate reductase (DHFR) with a THE riboregulator. DHFR is an essential gene for bacterial growth that is inhibited by the antibiotic trimethoprim (TMP). The line plot shows growth rate from OD_600_ measurements after TMP addition, normalized by the growth rate of each strain without TMP. Dashed lines indicate controls, either blank cells without a plasmid copy of DHFR (black) or cells with hairpin ON components (*d*=20, Figure 2b) that constitutively express DHFR (blue). See Supplementary Section 10 for additional experimental details. Data points represent three independent colonies, solid lines represent the mean, and shaded regions represent one standard deviation from the mean.

To demonstrate that 5ʹ UTR reprogramming could tune expression across *d* domain lengths for different protein CDSs, we designed THE riboregulators of 6 other fluorescent proteins spanning the green to red spectrum (Figure 6b). Changes in the fold activation of protein expression as a function of *d* domain length were remarkably consistent across these different CDSs (Supplementary Figure 34), even for sCFP3a, mScarlet-I, and mNeonGreen (Figure 6c), which only share approximately 30 % sequence identity and < 50 % sequence similarity^69^, and proteins with C-terminal degradation tags (Supplementary Figures 34 and 35). Beyond fluorescent proteins, we designed a THE riboregulator to control dihydrofolate reductase (DHFR) – an enzyme necessary for bacterial growth in minimal media (Figure 6d) – and used 5ʹ UTR reprogramming to control bacterial fitness levels across different concentrations of a DHFR-inhibiting antibiotic trimethoprim (TMP) (Figure 6e). Control of protein expression levels with *d* domain length also transferred to riboregulators with different 3’ UTRs (Supplementary Figure 39). These results highlight 5ʹ UTR reprogramming *via* TMSE as a simple but powerful tool for controlling protein expression that is readily transferable to many protein CDSs.

## CONCLUSION

Here we developed toehold exchange riboregulators as a genetically encoded tool for measuring TMSE reactions and controlling protein expression in living cells. These results represent the first cellular demonstration of genetically encoded TMSE circuits – merging key capabilities of DNA computing and cellular riboregulators. Intriguingly, we found circuit components characterized in IVT^47^ could be directly adopted for use in bacteria (Figure 2) with specificity, scalability, and composability largely preserved in the complex cellular environment (Figure 3). These properties allowed us to directly port multi-component circuits from the test tube to the cell (Figures 4 and 5). Further, we extend TMSE circuit capabilities by introducing a new, context-robust mechanism for regulating protein translation strength without changing mRNA sequence – 5’ UTR reprogramming (Figure 6). Together these results present THE riboregulators as versatile RNA devices with a highly adaptable design and genetically encoded TMSE circuits as a unified platform for molecular programming across environments^45,48^.

THE riboregulators allowed us to identify TMSE properties in *E. coli* that deviate from expected trends based on the thermodynamics of base pairing alone and contrast with previous IVT or cell-free transcription-translation results^45,48^. For example, increasing input toehold lengths beyond 6 bases decreased expression (Figure 2e and Supplementary Figures 18 and 19), inputs with 6-base toeholds facilitated displacement of outputs with 12-base toeholds, and increasing branch migration length increased leaky expression – even when clamping the *rbs* (Figure 2g and Supplementary Figures 20 and 21). Further, input and output strands required 3′ secondary structures for signal propagation, which is also required in cell-free expression systems^48^ but not in IVT^45^. These results could be due to circuit components interacting with RNA binding proteins, RNA chaperones, or endogenous RNAs that either help^70,71^ or hinder strand exchange^72^. RNases could also be involved; short, unstructured circuit input/output RNAs^73,74^ and long dsRNA strand exchange intermediates^75,76^ may be particularly susceptible to degradation. Encouragingly these trends are not strongly sequence dependent, resulting in consistent and predictable behavior and potentially allowing these bugs to be turned into features. For example, relatively short input toeholds driving displacement of long output toeholds enables flexible integration with other devices^25^ while maintaining sequence specificity.

Our results lay the foundation for directly implementing much larger and more sophisticated genetically encoded TMSE circuits in cells. The high specificity of TMSE supports a large number of orthogonal sequences but other scale-up challenges arise when operating in cells. For circuits that run entirely with TMSE, shallow and wide architectures, such as those used to build TMSE winner-take-all neural networks^39,40,42^, are well suited for cellular implementation. Building deep, multi-layered architectures, like those used for digital logic^38,41^, will likely require signal amplification between layers, which could be achieved by linking RNA strand exchange to transcription of inputs for the next layer^25,77,78^. For any architecture, resource competition, retroactivity, and cell burden^9^ will become issues as circuit size increases; controlling the timing of when circuit elements are expressed could help circumvent this challenge. Directly porting *in vitro* TMSE circuit designs to genetically encoded versions will also require predictable control of circuit component stoichiometries in cells. The thresholding-based classifier we developed in Figure 5 shows a relatively crude implementation for controlling relative concentrations, but predictably mapping DNA sequence to RNA concentration in cells will be crucial. THE riboregulators provide a powerful measurement tool to advance these capabilities.

While THE riboregulators serve as a powerful tool for measuring TMSE reactions in cells, they offer many advantages as a standalone biotechnology. We found THE riboregulators were sensitive to single-nucleotide mismatches introduced early in branch migration but tolerant of mismatches later in branch migration. These properties could support the development of either ultraspecific or highly flexible biosensors for single-nucleotide variants of disease or infectious organisms^79,80^. For example, the ancestral strain of SARS-CoV-2 can be distinguished from two other variants by different point mutations at the same genomic position^80^. A variant-specific THE riboregulator could be designed by placing the target base early in the branch migration domain, while a generalist THE riboregulator activated by all three variants could be designed by placing the degenerate position at the end of the branch migration domain. Developing these distinct biosensors requires only a single-base change in the canonical THE riboregulator design, rather than a different device design^26,27^, making it straightforward to integrate these biosensors into larger information processing circuits.

We also illustrated multiple routes to modulate protein expression with THE riboregulators without changing connectivity with other TMSE gates. For example, the *s* domain of a gate can tune expression nearly 5-fold for the same input, and the *d* domain of an input can tune expression nearly 100-fold for the same gate. Cell-free expression results^48^ support that these two methods act at different levels of gene regulation – the *s* domain operating post-transcriptionally by changing TMSE rate and the *d* domain operating translationally by changing protein expression strength. This level of versatility is not easily achieved with other riboregulators and could support applications in biomanufacturing^49,81^, biosensing^77,82^, and supramolecular assembly^83–86^. For example, the ratios of multiple proteins could be controlled using orthogonal THE riboregulators, each with input RNAs that have unique *d* domain lengths. TMSE cascades could control the order of protein expression with *s* domain and *d* domain lengths controlling the flux and expression strength at each layer. Further, inducible expression of input RNAs with different *d* domain lengths allows transcription and translation to be controlled independently and more predictably than existing methods^49^.

This work represents a substantial advance that promises to expand the capabilities of TMSE. For example, the continuous component turnover enabled by genetically encoded RNA circuits allows them to dynamically respond to changing environmental stimuli, enabling sophisticated information processing tasks difficult to achieve with *in vitro* TMSE circuits, such as unsupervised learning^42^. Genetically encoded TMSE circuits thus go beyond just bringing the powerful capabilities of TMSE circuits to living cells^30,31^, they expand what is possible with this molecular programming paradigm.

## MATERIALS AND METHODS

### DNA, bacterial strains, and chemicals

#### DNA

DNA oligonucleotide primers were ordered from Integrated DNA Technologies (IDT) with standard desalting. DNA sequences encoding RNA inputs and RNA gates were ordered as eBlock gene fragments from IDT. The Duet vectors used as plasmid backbones (pColADuet-1, pETDuet-1, pACYCDuet-1) were ordered from Novagen. The CymR^AM^ cassette used in the thresholding-based logic circuit was taken from pAJM.657 of the Marionette Sensor Collection^81^, a gift from Christopher Voigt (Addgene Kit #1000000137).

#### Strains

Electrocompetent DH5α cells for transformations were prepared in house (derived from ThermoFisher, cat. no. 18265017). Electrocompetent BL21 Star (DE3) cells for transformations were prepared in house (derived from Invitrogen, cat. no. C601003).

#### Chemicals

1000x stocks of the antibiotics kanamycin (Kan, Sigma, cat. no. K4000), carbenicillin (Carb, Invitrogen, cat. no. 10177-012), chloramphenicol (Cam, Acros, cat. no. 227921000), spectinomycin (Spec, Gold Biotechnology, cat. no. S-140-5), and trimethoprim (TMP, cat. no.) were prepared at 50 mg/mL in water (Kan), 100 mg/mL in water (Carb), 34 mg/mL in 75 % volume fraction ethanol (Cam), 50 mg/mL in water (Spec), and 25 mg/mL in DMSO (TMP). Isopropyl ß-D-1-thiogalactopyranoside (IPTG) stocks were prepared in water. Cumate stocks (4-isobpropylbenzoic acid from Tokyo Chemical Industry, cat. no. I0169) were prepared in 100 % ethanol.

### PCR, plasmid assembly, and cloning

#### PCR

PCRs were conducted with Phusion Flash Master Mix (ThermoFisher, cat. no. F-548) and 0.5 µmol/L of each DNA primer. THE riboregulators were assembled *via* overlap PCR of 0.02 ng/µL of eBlocks encoding a gate design and the protein coding sequence. The 27-base *c′* and *n* domains were used as the complementary domain for overlap PCR. PCRs of DNA inserts encoding input RNAs or upstream ctRSD gates were also conducted with 0.02 ng/µL (Supplementary Figure 4). For inserts, the following thermocycler protocol was executed: 30 cycles of a denaturing step (30 s, 98 °C), a primer annealing step (30 s, 60 °C), and an extension step (30 s, 72 °C). A 3 min extension at 72 °C was executed at the end of the program. Linear plasmid backbones for cloning were prepared *via* PCR of 0.2 ng/µL plasmid DNA and the following thermocycler protocol: 30 cycles of a denaturing step (30 s, 98 °C), a primer annealing step (30 s, 62 °C), and an extension step (120 s, 72 °C). A 5 min extension at 72 °C was executed at the end of the program. Unpurified backbone PCRs were then digested with a volume fraction of 2.5 % FastDigest DPNI (ThermoFisher, FD1703) for 1 h at 37 °C. PCR products and DPNI digested plasmid backbones were purified with a PureLink PCR purification kit (Invitrogen, cat. no. K310001). For overlap extension and backbone PCR products, a wash step with buffer B3 after binding with buffer B2 was included to remove non-specific products < 300 base pairs. The uncleavable ribozyme variants used in RT-qPCR experiments and the input variants with base substitutions were prepared similarly to linear backbones but with inverse PCRs of the plasmids encoding wild type sequences using primers that introduced the desired mutations. Inverse PCR products were then DPNI digested, purified, and ligated with 10 U/µL T4 ligase (New England Biolabs, cat. no. M0202).

#### Plasmid assembly and cloning

Purified DNA inserts were cloned into plasmid backbones with Gibson assembly^87^ using 30-base or 40-base homology domains derived from^88^ (Supplementary Section 3). Insert and backbone DNA were then mixed at a 3:1 molar ratio (backbone concentration of approximately 15 ng/µL) with Gibson assembly mix (New England Biolabs, cat. no. E2611L) and incubated at 50 °C for at least 1 hour prior to transformation into electrocompetent DH5α cells. Inserts from resulting colonies were sequence verified and 5 mL liquid cultures in LB with appropriate antibiotic were grown overnight for glycerol stocks and plasmid extraction. Plasmid extractions were conducted with a Miqron DNA purification robot (Galenvs Sciences) and Galenvs Sciences plasmid purification kits (cat. no. QPM0016) or Qiagen Spin Miniprep kits (cat. no. 27104). Extracted plasmids were then transformed into electrocompetent BL21 Star (DE3) cells for testing. Electrotransformations were conducted with MicroPulser Electroporation Cuvettes, 0.2 cm gap (BioRad, 1652086) using an Eporator (Eppendorf, 4309000027).

Additional details and schematics of inserts are shown in Supplementary Section 3. Primer and homology domain sequences are in Supplementary File 1. GenBank files of plasmids are in Supplementary File 2. Annotated plasmids can also be accessed on Benchling: https://benchling.com/sschafft1/f_/DkQ1GTHono-the-riboreg-ecoli-2025/

### Cell culture and RNA expression

#### Cell culture

Three independent colonies from plasmid transformations into BL21 Star (DE3) cells were selected to inoculate 600 µL of Luria broth medium (LB, ThermoFisher, cat. no. BP1426) with appropriate antibiotics in 96-well, deep round bottom plates (ThermoFisher, cat. no. 278743). Plates were sealed with a gas permeable membrane (4titude, cat. no. 4ti-0597/ST) and incubated overnight (approximately 16 h) at 37 °C, 1000 rpm in an Eppendorf ThermoMixer C with an Eppendorf ThermoTop lid. The next day cells were diluted 50-fold into 300 µL of fresh LB with appropriate antibiotics in a custom 96-well microbial growth plates (Agilent, mat. no. S7898A). Cells were grown for a total of 6 h in the plates at 37 °C in a Neo Synergy2 (Agilent BioTek) with continuous double orbital shaking (559 cycles per minute). Measurements for optical density (600 nm and 700 nm), CFP (ex: 430 nm, em: 475 nm, bandwidth: 10 nm, gain: 85), GFP (ex: 490 nm, em: 515 nm, bandwidth: 10 nm, gain: 75), RFP (ex: 560 nm, em: 601 nm, bandwidth: 20 nm, gain: 100) taken every 10 min. Fluorescent measurements were taken from the bottom of the plate using the Neo Synergy2 monochromator.

#### RNA expression

Unless otherwise stated, experiments with a single input and a single gate induced T7 RNAP expression with 100 µmol/L of IPTG at the same time as the 50-fold dilution of overnight cultures. For multi-layer cascades and logic circuits, 100 µmol/L of IPTG was added 3 hours after dilution of the overnight cultures. For the logic circuit in Figure 5, cumate was added at the same time as the dilution of the overnight cultures. See Supplementary Section 4 for a schematic of the cell culture workflow and representative growth curves with different induction conditions.

### Flow cytometry

After 6 h of growth in a Neo Synergy2, cells were diluted 40-fold into 200 µL of phosphate-buffered saline (PBS, Invitrogen, cat. no. AM9625) supplemented with 170 µg/mL of Cam in 96 well U-bottom plates (Corning, cat. no. 351177). For assays of the thresholding-based logic circuit, which included a plasmid with Cam resistance, PBS supplemented with Spec was used. Flow cytometry was conducted in an Attune NxT acoustic focusing cytometer equipped with violet (ex: 405 nm), blue (ex: 488 nm), and yellow (ex: 561 nm) lasers and a CytKick Max plate auto sampler (Life Technologies). 25 µL of cell samples, typically corresponding to > 100 000 single cells, were measured. Biological replicates were analyzed sequentially followed by a 150 µL PBS wash before measuring the next set of replicates (Supplementary Figure 13).

Automated singlet gating was performed with FlowGateNIST^89^. Fluorescent calibration beads (Spherotech, cat. no. RCP-30-5A) were used to convert CFP fluorescence signal from the VL2 channel (em: 512 nm ± 25 nm) to molecules of equivalent fluorophore (MEF) as described in Ref^89^. The BV510 fluorophore on the beads was used for calibrating the CFP signal, and cells without a CFP expressing plasmid were used as the background for calibration. Distributions of fluorescent cells in figures represent unmodified histograms after calibration to MEF normalized by the total singlet count after automated gating. Bar plots represent geometric means of these distributions (Supplementary Figure 13). Cells expressing GFPs were analyzed on the BL1 channel (em: 530 nm ± 30 nm) and cells expressing RFPs were analyzed on the YL2 channel (em: 620 nm ± 15 nm).

### DHFR growth-based assay

Dihydrofolate reductase (DHFR) growth assays were conducted in M9 medium (1x Difco M9 minimal salts (Fischer Scientific, cat. no. 248510) supplemented with 0.1 mmol/L CaCl_2_, 2 mmol/L MgSO_4_, 20 g/L casamino acids, and a volume fraction of 0.4 % glycerol). See Supplementary Figure 36 for plate layout. We found this assay was not compatible with rich LB medium (Supplementary Figure 37).

Individual colonies from glycerol stock streaks were picked to inoculate 5 mL of M9 medium supplemented with antibiotics (Kan: 50 µg/mL and Carb: 100 µg/mL). Liquid cultures were grown overnight at 37 °C, 300 rpm. In a microbial assay plate, overnight cultures were diluted 50-fold into M9 medium supplemented with antibiotics and 100 µmol/L of IPTG to induce expression of the RNA constructs. Plates were sealed with a gas permeable membrane (Sigma, cat. no. Z380059) and incubated in an Epoch 2 microplate reader (Agilent BioTek) at 37 °C with continuous double orbital shaking (807 cycles per minute). Optical density measurements at 600 nm were taken every 10 min. After 240 min, TMP was added to the media to the final concentrations specified in Figure 4. The optical density measurements over the next 90 min (240 min to 330 min) were used to fit a growth rate at each TMP concentration. For replicates, the assay was repeated on three separate days with individual colonies picked from the same glycerol stock streaks. Additional details are in Supplementary Figure 38.

### RT-qPCR measurements of ribozyme cleavage

RNA extraction, ribozyme protection, and RT-qPCR were conducted primarily as previously described^60^ with notable differences discussed below.

#### RNA production and extraction for RT-qPCR

Plasmids encoding a THE riboregulator or a corresponding uncleavable control variant with a point mutation rendering the ribozyme inactive were individually transformed into BL21 Star (DE3) cells. Single colonies were used to inoculate 2 mL overnight liquid cultures in LB with appropriate antibiotic. Overnight cultures were then diluted 100-fold into 10 mL of LB with appropriate antibiotic and 100 µmol/L of IPTG. Cells were then grown at 37 °C, 300 rpm to an optical density at 600 nm of approximately 0.6 prior to harvesting for RNA extraction. 2 mL of culture was mixed with 4 mL of RNAprotect reagent (Qiagen, cat. no. 1018380) and incubated at room temperature for 5 min. The sample was then centrifuged for 10 min at 5000 g, 4 °C. The cell pellet was resuspended in 100 µL of 50 mmol/L EDTA. Resuspended cells were then lysed with 1 mL of TRI reagent (ZymoResearch, cat. no. R2050) for 5 min at room temperature. If not used immediately, lysed cells were stored at -80 °C.

The RNA extraction protocol was modified from previously described^60^ to remove the DNase digestion. Total RNA was extracted using 1-Bromo-3-chloropropane (Sigma, cat. no. B9673) separation and subsequently purified using Zymo-Spin IIICG columns (ZymoResearch, cat. no. C1006-50-G) and wash buffer (ZymoResearch, cat. no. C1001-50). Purified RNA was eluted in nuclease free water (Ambion, cat. no. AM9938), mixed with an equal volume of RNA storage solution (ThermoFisher, cat. no. AM7001), and stored at -80 °C for later use.

To prevent ribozyme cleavage during reverse transcription, the blocking oligo procedure described previously^60^ was followed exactly. Briefly, 17.1 ng/µL of extracted total RNA was mixed with 157 µmol/L of blocking oligonucleotide. Samples were then heated to 90 °C for 5 min and subsequently cooled to 20 °C at a rate of -1 °C per minute. This solution was then diluted 1250-fold in RNA storage solution. This dilution corresponds to a final RNA concentration of approximately 14 pg/µL based on the concentration of RNA prior to the blocking oligonucleotide addition. We found the high concentration of blocking oligonucleotide interfered with Qubit measurements of RNA concentration, preventing remeasurement of concentrations after blocking.

#### RT-qPCR

The DNA primers for measuring ribozyme cleavage with RT-qPCR were designed as previously described (Supplementary Figure 14). RT-qPCR experiments were conducted with a SuperScript III Platinum SYBR Green One-Step Kit (ThermoFisher, cat. no. 11736059) in an Applied Biosystems ViiA 7 Real-Time PCR System using 96 well PCR plates (Applied Biosciences, cat. no. N8010560). RT-qPCR reactions contained a final concentration of 3 mmol/L of MgSO_4_, 0.2 mmol/L of each dNTP, 50 nmol/L of ROX reference dye, and 0.2 μmol/L of forward and reverse primers. A final concentration of approximately 5 pg/µL of total RNA with blocking oligonucleotide was used (based on the RNA concentration before blocking oligonucleotide addition). RT-qPCR experiments were initiated with a 10 min reverse transcription step at 50 °C. The following thermocycler protocol was then executed: an initial denaturation step (5 min, 95 °C) followed by 40 cycles of a denaturing step at (15 s, 95 °C) and an annealing/extension step (60 s, 60 °C). Fluorescence readings were taken during the annealing/extension step of each cycle. After 40 cycles, samples were heated from 60 °C to 95 °C for melt curve analysis.

Schematics illustrating primer binding sites, primer efficiency experiments, and RT-qPCR control experiments are in Supplementary Section 5. Primer sequences are in Supplementary File S1.

## Supporting information

DNA sequences

Plasmid GenBank Files

Sequence compiler script

## AUTHOR CONTRIBUTIONS (CRediT)

**AOS**: Conceptualization (THE riboregulator applications), Methodology and Investigation (All mismatched input experiments, Plasmid construction and transformations, Cell culture and flow cytometry), Writing – original draft

**FP:** Methodology and Investigation (Experiments with *d* domains across protein CDSs, Plasmid construction and transformations, Cell culture and flow cytometry), Writing – review and editing

**OBV**: Methodology and Investigation (Plasmid construction and transformations, Cell culture and flow cytometry)

**MEW**: Methodology and Investigation (DHFR growth assay), Writing – review and editing

**JMH**: Methodology and Investigation (Plasmid construction and transformations, RT-qPCR) **NYA**: Methodology and Investigation (RT-qPCR)

**SWS**: Conceptualization, Data curation, Formal analysis, Investigation, Methodology, Project administration, Software, Supervision, Visualization, Writing – original draft

## DATA AND CODE AVAILABILITY

Flow cytometry data for this study will be available on Zenodo.

Data analysis and visualization was conducted in Python (3.8.18) in the Spyder IDE (5.4.3). Example analysis and plotting code is available in Supplementary File S3. Additionally, the ctRSD simulator 2.1 package (https://ctrsd-simulator.readthedocs.io/en/latest/SeqCompiler.html) contains a sequence compiling function to stitch together sequences for any combination of sequence domains explored in this study. Scripts for generating all sequences in this study are available in Supplementary File 3. A Google CoLab for running the sequence compiler is available at: <u>CoLab link</u>

## CONFLICTS OF INTEREST

SWS is an inventor on two patent applications pertaining to ctRSD circuits (Application number PCT/US2022/053229) and THE riboregulators (Application number PCT/US25/13610). The authors declare no other conflicts.

## ACKNOWLEGDMENTS

The authors thank Thomas Gorochowski, Shivang Joshi, Eugenia Romantseva, and Elizabeth Strychalski for insightful discussions throughout the project and feedback on the manuscript. In addition, the authors thank Svetlana Ikonomova and David Ross for helpful discussions about the DHFR growth assay. SWS also thanks Bright Eyes, whose album *Digital Ash in a Digital Urn* was an inspiration throughout the study.

## Disclaimer

Certain commercial entities, equipment, or materials may be identified in this document to describe an experimental procedure or concept adequately. Such identification is not intended to imply recommendation or endorsement by the National Institute of Standards and Technology, nor is it intended to imply that the entities, materials, or equipment are necessarily the best available for the purpose. Official contribution of the National Institute of Standards and Technology; not subject to copyright in the United States.

## Notes

### Summary of Updates

Additional experiments characterizing toehold exchange riboregulator specificity (Figure 3). Additional experiments demonstrating how toehold mediated strand exchange can be leveraged to predictably regulate translation strength for sequence and function diverse proteins without changing mRNA sequences (Figure 6). Improvements to clarity and motivation throughout - particularly for the multi-input classifier circuit (Figure 5).

