## Supplemental Information for "Genetically encoded RNA strand exchange circuits for programmable protein expression and information processing in cells"

### Supplementary Information

*Disclaimer:* Certain commercial entities, equipment, or materials may be identified in this document to describe an experimental procedure or concept adequately. Such identification is not intended to imply recommendation or endorsement by the National Institute of Standards and Technology, nor is it intended to imply that the entities, materials, or equipment are necessarily the best available for the purpose. Official contribution of the National Institute of Standards and Technology; not subject to copyright in the United States.

#### 1 Component nomenclature and generating sequences

Supplementary Figure 1 provides an overview of the full nomenclature for all components in this study. This nomenclature, albeit complicated, uniquely describes the sequence identity of every component. The sequences of each domain, separated in domain specific tabs are provided in `ctRSD_domains_list_210.xls` and at: https://github.com/usnistgov/ctRSD-simulator/blob/main/ctRSD-simulator-2.0/Sequence%20Compiler/ctRSD_domains_list_210.xls

The full nomenclature can also be used to generate component sequences for any combination of domains using sequence compiler software we developed in Python. The full documentation for the sequence compiler can be found at:
https://ctrsd-simulator.readthedocs.io/en/latest/SeqCompiler.html

A Google CoLab for running the sequence compiler is available at: CoLab link

All gene fragment sequences ordered for this manuscript are in Supplementary File 1.

GenBank files of all plasmids used in this study are in Supplementary File 2.

Annotated plasmids can also be accessed on Benchling:
https://benchling.com/sschafft1/f_/DkQ1GTHono-the-riboreg-ecoli-2025/

**a**

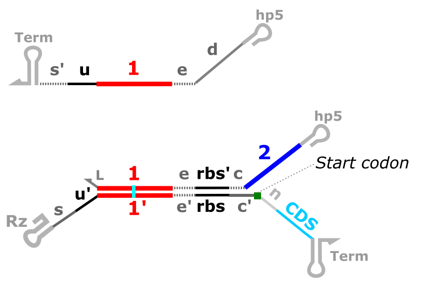

**b**

| Domain name | Length | Description |
|---|---|---|
| *hp5* | *Variable* | *5' hairpin:* *Promote uniform transcription rates and stability* |
| *d* | *1-20* | *Distal domain:* *Provide ssRNA standby site for ribosome binding* |
| * *e* | *0-6* | *Extended branch migration domain:* *Extend dsRNA duplex or output toehold* |
| *1,2 (3,4,N)* | *10* | *Branch migration domain:* *Promote strand exchange specificity* |
| *u (v,w,rbs')* | *6* | *Toehold:* *Facilitate strand exchange* |
| *rbs* | *6* | *Ribosome binding sequence:* *Interacts with 16S rRNA of ribosome to facilitate translation* |
| *rbs'* | *6* | *rbs complementary domain:* *Sequester rbs to prevent translation* |
| *Term* | *Variable* | *Terminator:* *Terminate transcription* |
| *s* | *0-10* | *Spacer:* *Reduce steric hindrance for input binding* |
| * *s'* | *0-10* | *Spacer complementary domain:* *Extend the input toehold length* |
| *Rz (R3,Rg,Rh,Rm)* | *60-100* | *Self-cleaving ribozyme:* *Produce dsRNA gates* |
| *L* | *3* | *Ribozyme linker:* *Facilitate efficient cleavage; or a hairpin for ouput stability* |
| * *c* | *0-6* | *Clamp:* *Bury the rbs* |
| *c'* | *6* | *rbs to AUG linker:* *Optimal spacing for translation initiation* |
| *n* | *18* | *N-terminal linker:* *Provide ssRNA footprint for ribosome* |
| *CDS* | *Variable* | *Protein coding sequence:* *Encodes for protein expression after strand exchange* |

**c**

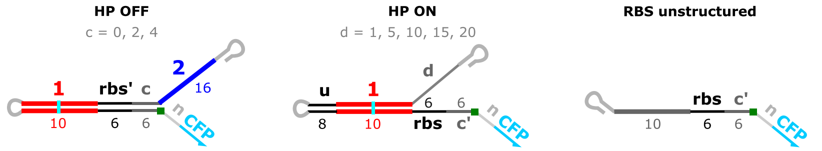

**d**

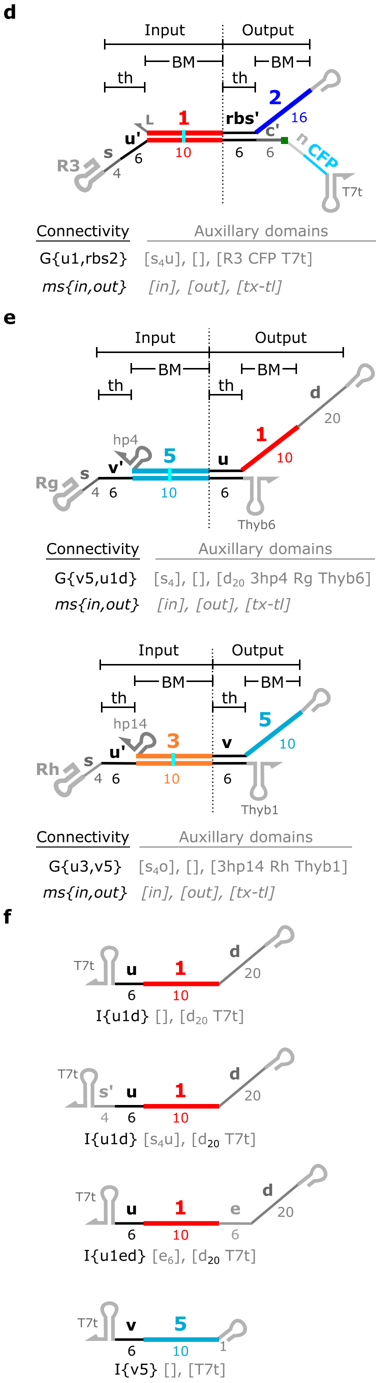

**Supplementary Figure 1:** Definitions and functions of the domains of toehold exchange riboregulators and their inputs used in this study. (**a**) Schematics of an input and THE riboregulator with all of the relevant domains from this study shown. The domains with dashed lines are optional and not necessary or not conventionally used compared to the 'canonical' THE riboregulator design. (**b**) Domain definitions and desired functions, along with their typical lengths in bases. The domain names with * to their left correspond to the non-necessary domains illustrated with dashed lines in panel a. (**c**) Control constructs used in this study.

*… Continued on next page …*

(**d**,**e**,**f**) Schematics of a toehold exchange THE riboregulator gate (panel c), upstream ctRSD gates (panel d), and inputs (panel e) along with their expanded nomenclature. THE riboregulators and ctRSD gates are composed of input and output domains, which each possess two subdomains: a toehold (th) subdomain for initiating strand displacement reactions and a branch migration (BM) subdomain for strand displacement specificity. Bold numbers and letters above the gate represent domain sequence identity and nonbold numbers below the gate represent domain length in bases. Connectivity names specify the molecular species (ms: G{ } for gate, I{ } for input, O{ } for output, *etc.*) and how it is connected to other species with input and output domains specified within the first and second position inside the curly brackets, separated by a comma. Auxiliary domains are specified after the connectivity name. These are domains that are not related to how the species connects to another species (*via* base pairing) or are not explicitly defined in the connectivity name. The convention is to group different types of auxiliary domains together and separate these groupings by a comma. The first grouping is for domains related to the input portion of the species (input extensions (*e*) and spacers / extended toeholds (s)), the second grouping is for domains related to the output portion of the species (output extensions (*e*)), and the last grouping is for domains related to transcriptional encoding or translation properties of the species. Note we specified the *d* domain in the connectivity name as part of the output because it *implies* a connection to a THE riboregulator given its role in protein translation. However, we specify the identity of the *d* domain in transcription-translation [ ] of the auxiliary domains because it influences translation and isn't involved in base pairing with other species or modulating the kinetics of interactions between species. Within the auxiliary groupings [ ] the convention is to list domains in the 5′ to 3′ order they appear within the species. For auxiliary domains that can be different lengths *e.g.*, *s*, *e*, *d*, *etc*. the convention is to specify the domain type first, the length in bases second as a subscript, and any sequence identifier last. For example, $s_4u$ is a spacer that is 4 bases in length and an extension of the *u* toehold. Across connectivity and auxiliary domain names, numbers in subscripts refer to the length of the domain in bases and all numbers not in subscripts are part of the sequence identifier. Specific subdomain sequences are part of the ctRSD sequence compiler and listed in the `ctRSD_domains_list_210.xlsx` file.

### 2 Toehold exchange riboregulator design notes

#### 2.1 Riboregulator design notes

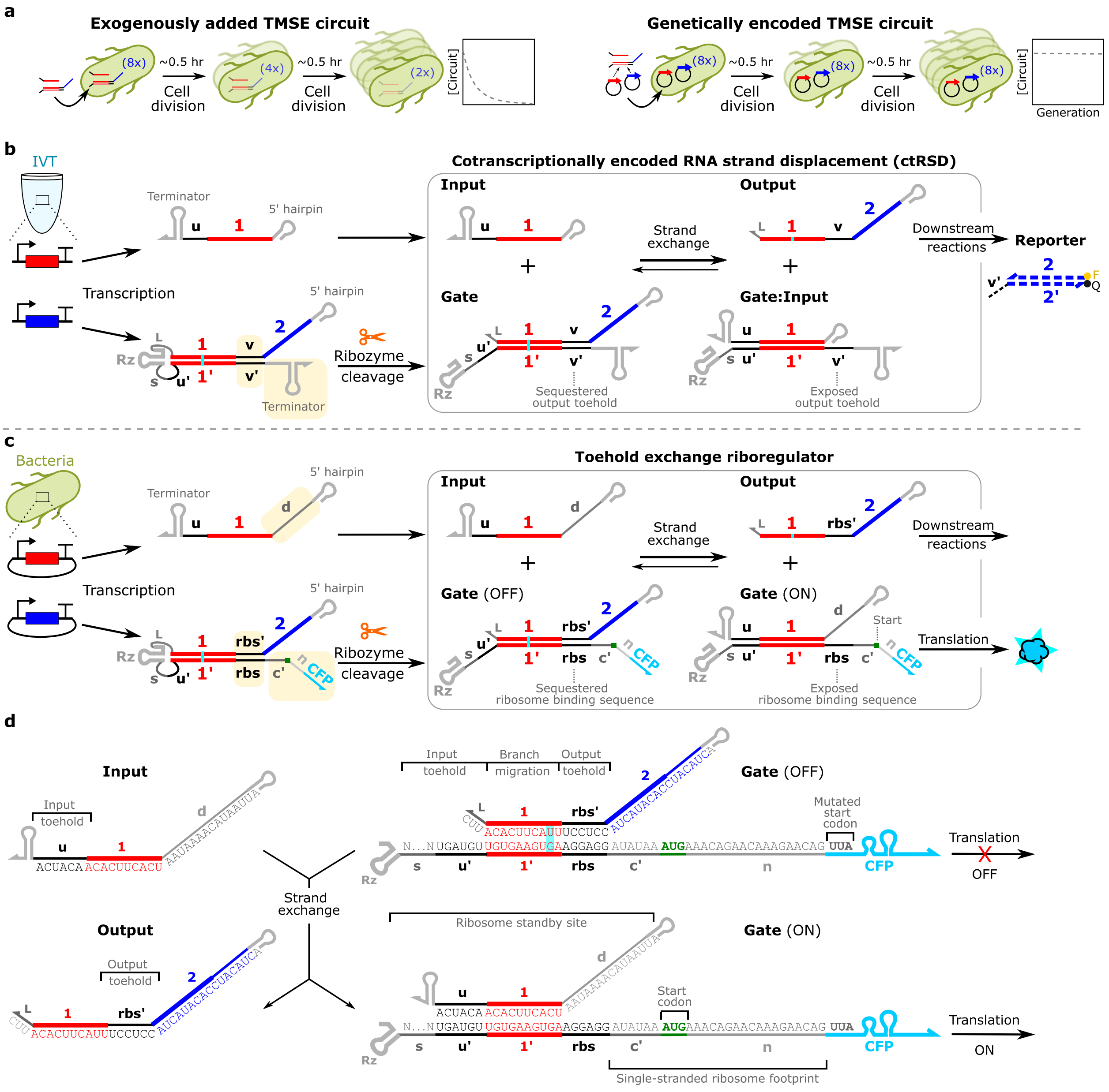

**Supplementary Figure 2:** Comparison of TMSE circuits, genetically encoded TMSE circuits (ctRSD circuits) and toehold exchange riboregulators. (**a**) Adding pre-prepared TMSE components at fixed concentrations to cells will result in decreasing concentrations as cells divide, while encoding components on plasmids allows circuit instructions to be propagated across generations. (**b**) ctRSD toolkit schematic showing the design as typically employed *in vitro* transcription. The three domains highlighted in yellow are changed to convert from the ctRSD toolkit to a toehold exchange riboregulator. (**c**) Toehold exchange riboregulator schematic. (**d**) Sequence schematics of the components in (panel b). The *s* domain is shown as Ns as different lengths and sequences are used.

To modify gates from the ctRSD toolkit[1,2] to be riboregulators *i.e.*, regulate translation of a protein coding sequence, two gate domains were changed (Supplementary Figure 2).

First, the output toehold of the gate was changed from a semi-arbitrary sequence to a strong *E. coli* ribosome binding sequence (*rbs*) (Supplementary Figure 2). Ribosome binding sequences, also known as Shine-Dalgarno sequences, hybridize to a portion of the 16S ribosomal RNA in *E. coli* to facilitate initiation of translation. Because ribosome binding sequences in *E. coli* (and many other prokaryotes) are purine-rich[3,4], we were able to maintain the 3-base sequence constraint typically employed for the output strand of ctRSD gates *i.e.*, the output strand is still only composed of C, A, U bases to prevent misfolding[1,2]. There are 9 bases in *E. coli* 16S rRNA that drive translation initiation through hybridization with a ribosome binding sequence on an mRNA[5], we designed the *rbs* domain to be 6 bases complementary to the central stretch of the relevant 16S rRNA. This keeps the *rbs'* output toehold the same length as the *u*, *v*, *w* toeholds commonly used in ctRSD circuits.

Second, the sequence downstream of the output toehold of the gate, which is typically a terminator sequence, was changed to a protein coding sequence (CDS) preceded by a 27-base linker sequence (Supplementary Figure 2). The linker sequence includes two domains: *c'*, which correctly positions the start codon downstream of the *rbs*[6], and *n*, which is designed to prevent secondary structure in the footprint of the bound ribosome to promote efficient translation initiation[7].

With this design the gate should cotranscriptionally fold into a stable hairpin that sequesters the ribosome binding sequence, preventing translation of the downstream protein coding sequence. After ribozyme cleavage, the dsRNA duplex of the gate should continue to sequester the *rbs* unless a complementary input strand is present. A complementary ssRNA input can hybridize to the input toehold of the gate to initiate a strand displacement process that results in dissociation of the output strand and exposure of the *rbs*, enabling translation. In this process the hybridization of the input toehold is exchanged for the dissociation of the *rbs'* toehold, so we term these gates “toehold exchange (THE) riboregulators”.

The translation strength of a given mRNA depends not only on the degree of complementarity of the *rbs* to the 16S rRNA, but also the structure of the RNA surrounding the *rbs*.

As discussed above, the structure of the region downstream of the *rbs* is important, with unstructured regions that can accommodate the footprint of the bound ribosome (approximately 16 bases) producing the highest translation initiation rates[7]. This is why we included the *n* domain in our design, to ensure there was minimal secondary structure in the ribosome footprint across different protein coding sequences. We used NUPACK[8] to confirm that there was little to no predicted RNA secondary structure in the first 15 bases to 20 bases downstream of the start codon for all CDSs used in this study.

Additionally, the structure of the RNA upstream of the *rbs* influences translation strength. In our case there is a 16-base dsRNA helix directly upstream of the *rbs*, which should severely inhibit translation initiation. Prior to hybridizing to the *rbs* domain of an mRNA, ribosomes often bind non-specifically to unstructured regions upstream of the *rbs* known as standby sites[9].

Unstructured standby sites of 16 to 20 bases can promote high translation rates, even in the presence of strong secondary structure around the downstream *rbs*[10,11]. Based on this information, we hypothesized that extending the 5′ end of the input in our system should increase the translation initiation rate by serving as an unstructured standby site in the ON gate (*d* domain in Supplementary Figure 2 and Figure 2). We designed this *d* domain with a C, A, U restricted sequence to ensure minimal secondary structure. It is also possible that some of the input RNAs could be captured by a ribosome[12] before initiating strand exchange; after completing strand exchange they would then be localized near the *rbs* on the gate.

Before testing our designs experimentally, we assessed their performance with the RBS Calculator[13], software designed to predict translation initiation rates from RNA sequence using the sequence and structure considerations discussed above. To simplify the system for computational exploration, we input sequences designed to be hairpin equivalents of the OFF and ON gates, rather than dsRNAs (Supplementary Figure 2).

We explored 3 different HP OFF designs, one with only the 6 bases of the *rbs* sequestered and two with an additional 2 or 4 bases paired downstream of the *rbs* to 'clamp' the *rbs* helix shut. As expected, increasing this clamping region was predicted to reduce translation initiation of the OFF state. We explored 5 different HP ON designs with *d* domains spanning 1 to 20 bases. As we hypothesized, increasing the length of the *d* domain was predicted to increase the translation initiation rate (Supplementary Table 1).

The translation initiation rates from the RBS Calculator have arbitrary units, so to put them in context we compared the results to predictions of a toehold switch (THS) design[14] regulating CFP. Our THE riboregulators were predicted to have higher OFF expression than the THS riboregulator, but with $d > 10$ our designs were predicted to exceed the ON expression of the THS designs. We did not test this comparison experimentally, but a rigorous comparison would be an interesting future study.

Comparing these predictions to our experimental results in Figure 2b of the main text, we found all our HP OFF constructs had similar expression to the background fluorescence of blank cells and the HP ON results followed the increase in expression predicted by the RBS Calculator.

**Supplementary Table 1:** Predicted standby free energy and translation rates of RNA components (Supplementary Figure 2d) in *E. coli* BL21 (DE3) from the RBS Calculator[13] (Version 2.1). The first 120 bases of the sCFP3A CDS were included in the prediction. RBS Calculator predictions are for the intended start codon, not any spurious start sites. $\Delta G_{standby}$ indicates a free energy penalty for ribosome binding to a structured standby site. Note this penalty is predicted to go away once *d* reaches 20 bases. HP OFF (-*rbs*) is a component that replaced the *rbs* domain to the *w* domain, which has very little complementarity with *E. coli* 16S rRNA. UTR unstructured is a component without secondary structure upstream of the *rbs*. The toehold switch (THS) constructs are taken from Ref[14]. sCFP3A molecules of equivalent fluorophore (MEF) indicates the average geometric mean of flow cytometry distributions from three colonies ± one standard deviation.

| **Component** | **$\Delta G_{standby}$ (kcal/mol)** | **Predicted translation initiation rate (AU)** | **sCFP3A (MEF)** |
|---|---|---|---|
| HP OFF (*c*=0) | 0.00 | 1362 | 55.41 ± 0.66 |
| HP OFF (*c*=2) | 0.00 | 355 | 46.77 ± 0.60 |
| HP OFF (*c*=4) | 0.00 | 151 | 45.23 ± 0.56 |
| HP OFF (-*rbs*) | 0.00 | 72 | 50.44 ± 0.41 |
| Blank cells | N/A | N/A | 53.40 ± 0.45 |
| HP ON (*d*=1) | 7.72 | 2310 | 618.7 ± 22.84 |
| HP ON (*d*=5) | 5.65 | 5846 | 1905 ± 225.1 |
| HP ON (*d*=10) | 3.32 | 16684 | 4344 ± 320.1 |
| HP ON (*d*=15) | 0.72 | 53788 | 5540 ± 738.6 |
| HP ON (*d*=20) | 0.00 | 74452 | 9080 ± 735.6 |
| UTR unstructured | 1.09 | 262532 | 6129 ± 230.1 |
| THS OFF (ACTSII_1) | 2.81 | 41 | N/A |
| THS ON (ACTSII_1) | 2.74 | 28889 | N/A |

#### 2.2 Design considerations for protein coding sequences (CDSs)

The *n* domain we included to enhance translation initiation results in an additional 6 amino acids being added to the N-terminus of the protein encoded by the riboregulator. For the proteins we tested, fluorescent proteins and *E. coli* dihydrofolate reductase (DHFR), N-terminal extensions do not greatly influence protein function. However, these additions could cause issues for certain proteins, and this should be investigated on a case-by-case basis. The *n* domain is not strictly necessary and can be removed or reduced should it become problematic for a specific protein of interest.

Introduction of the *n* domain resulted in two start codons, the one we introduced close to the *rbs* domain and the one to start the protein CDS. To prevent any spurious translation initiation[4] at the CDS start codon, which would result in leak expression from our riboregulator, we mutated ATG (M) to TTA (L) (Supplementary Table 2). We chose this mutation because the two amino acids have similar biophysical properties and the TTA codon has a very low rate of translation initiation[15]. We did not explicitly validate whether this change was necessary, it likely depends on many factors, such as choice of the *n* domain sequence, choice of CDS, choice of organism. It is also possible some proteins cannot tolerate this mutation.

We also took precautions to reduce the chance of spurious translation initiation within the protein CDSs we tested. For example, the original sequences of mCherry and mNeonGreen (and many of their relatives) have cryptic translation initiation sites due to inclusion of an eGFP N-terminal peptide during their development[16,17]. We mutated the alternative start codon in these proteins to avoid translation initiation at these alternative sites, which would have led to substantial leak in our system (Supplementary Table 2).

Using these design principles, we tested a total of 7 different fluorescent protein sequences and found that all designs resulted in a > 40 ON/OFF ratio. Further, the dependence of expression strength on the length of the *d* domain on the input RNA held across all the sequences (Supplementary Figure 34).

The components of RNA circuits should constantly be turned over in the cell, but commonly used fluorescent proteins do not turnover as they are highly stable. Many C-terminal degradation tags[18] can be added to increase fluorescent protein turnover, which has been used in other RNA circuit work in bacteria to reduce leaky expression[14,19]. We found adding an ASV degradation tag[18] reduced the amount of leak observed across input sequences. It also produced more normal distributions, as some of the distributions without the degradation tag were skewed or had secondary peaks (Supplementary Figure 35). These types of degradation tags could be useful for studying dynamic systems and may lower toxicity some toxicity of overexpression.

Lastly, in fast growing organisms like *E. coli* the maturation time of a fluorescent protein is crucial for obtaining high signal within single cells[20], so we mainly tested proteins with short maturation times (Supplementary Table 2).

**Supplementary Table 2:** Analysis of protein CDS and N-termini for the fluorescent proteins used in this study. Green highlighted amino acid are N-terminal amino acids of eGFP and gray highlighted amino acids are a short linker. These regions were originally derived from eGFP as a linker to effectively tag the C-termini of proteins with other fluorescent proteins[16]. Green bases represent intended start codons and red bases indicate potential spurious start codons further along in the CDS. Bold amino acids and bases were changed in this study to reduce the chances translation initiation at a location other than the intended location. Underlined bases have with complementarity to *E. coli* 16S rRNA. The mat. time column indicates the reported time in minutes to reach 50 % maturation at 37 °C[20].

| | | |
|---|---|---|
| | **N-termini of proteins from FPbase**[21] | |
| eGFP_N-linker | M V S K G E E D N M A - FP TAG<br>ATG GTC AGT AAG GGT GAG GAG GAT AAC ATG GCC | |
| | | Mat. time (min) |
| GFPmut3_ASV | M R K G E E L F T G V V P I L V<br>ATG CGT AAA GGA GAA GAA CTT TTC ACT GGA GTT GTC CCA ATT CTT GTT | 4.1 |
| sfGFP | M S K G E E L F T G V V P I L V<br>ATG AGC AAA GGA GAA GAA CTT TTC ACT GGA GTT GTC CCA ATT CTT GTT | 13.6 |
| sCFP3A | M V S K G E E L F T G V V P I L V<br>ATG GTC AGT AAA GGT GAA GAA TTA TTT ACC GGT GTT GTA CCG ATA TTA GTC | 6.4 |
| mNeonGreen | M V S K G E E D N M A S L P A T H<br>ATG GTC AGT AAA GGT GAA GAA GAT AAC ATG GCC AGC TTG CCA GCA ACT CAT | 10.9 |
| mCherry2 | M V S K G E E D N M A I I K E F M<br>ATG GTC AGT AAA GGT GAA GAA GAT AAC ATG GCC ATC ATC AAG GAG TTC ATG | 22.8 |
| mScarlett-I | M V S K G E A V I K E F M<br>ATG GTC AGT AAA GGC GAA GCA GTT ATC AAA GAG TTC ATG | 25.7 |
| mRFP1 | M A S S E D V I K E F M<br>ATG GCC TCT TCT GAG GAC GTC ATC AAG GAG TTC ATG | 21.9 |
| | **N-termini of proteins used in this study** | |
| *n* domain | M K Q N K E Q<br>ATG AAA CAG AAC AAA GAA CAG | |
| GFPmut3_ASV | **L** R K G E E L F T G V V P I L V<br>**TTA** CGT AAA GGA GAA GAA CTT TTC ACT GGA GTT GTC CCA ATT CTT GTT | |
| sfGFP | **L** S K G E E L F T G V V P I L V<br>**TTA** AGC AAA GGA GAA GAA CTT TTC ACT GGA GTT GTC CCA ATT CTT GTT | |
| sCFP3A | **L** V S K G E E L F T G V V P I L V<br>**TTA** GTC AGT AAA GGT GAA GAA TTA TTT ACC GGT GTT GTA CCG ATA TTA GTC | |
| mNeonGreen | **L** V S K G E E D N **L** A S L P A T H<br>**TTA** GTC AGT AAA GGT GAA GAA GAT AAC **TTA** GCC AGC TTG CCA GCA ACT CAT | |
| mCherry2 | **L** V S K G E E D N **L** A I I K E F M<br>**TTA** GTC AGT AAA GGT GAA GAA GAT AAC **TTA** GCC ATC ATC AAG GAG TTC ATG | |
| mScarlett-I | **L** V S K G E A V I K E F M<br>**TTA** GTC AGT AAA GGC GAA GCA GTT ATC AAA GAG TTC ATG | |
| mRFP1 | **L** A S S E D V I K E F M<br>**TTA** GCC TCT TCT GAG GAC GTC ATC AAG GAG TTC ATG | |

#### 3 Plasmid schematics and plasmid assembly notes

The pDuet vectors (Novagen), primarily pColADuet and pETDuet – along with pACYC for experiments in Figure 5 – served as the backbones for plasmid construction in this study. The two T7 promoter sites and the T7 terminator site present in these vectors were removed *via* inverse PCR and sequences encoding for RNA circuits components were inserted with Gibson assembly. THE riboregulators were inserted into the pColA backbone and RNA inputs and upstream ctRSD gates for multi-layer cascades were encoded on the pET backbone (Supplementary Figure 4). The pDuet backbones all contain a copy of LacI, which could reduce basal expression of T7 RNAP in the BL21 Star (DE3) *E. coli* strain used in this study.

DNA encoding RNA circuit components were ordered as eBlocks from IDT. For most the THE riboregulators only the gate portion (through the *n* domain) of the sequence was ordered, and this sequence was stitched to the relevant protein CDS using overlap PCR (Supplementary Figure 4). We typically did this to obtain linear templates for rapid prototyping in cell-free systems, but 5′ UTR could be put into backbone plasmids with a protein CDS using the *c′* and *n* domains as a homology domain (27 bases), which we found worked successfully for other projects. Per the DNA provider, the sequences had to be 300 bases, so filler sequences were appended to meet this length requirement when necessary. As in previous work[1,2], all gates contain a GU wobble in their dsRNA stem to enable synthesis of the long complementary regions. Example annotated sequences for eBlocks are shown below.

**Legend:** **$U_G$**, **$U_D$**, **$U_E$**, **T7p**, **filler**, RNA COMPONENT

```
G_1 ( G{u1,rbs2} [s4u],[],[R3] ):

ataggagccgcgaagtctaacgctgctctgggctaactgtccgcatctagacttaactgagatattaccatagatgactagccatt
cctctagatactacgactagcatacTTCTAATACGACTCACTATAGGGAGATTCGTCTCCCACTACATCCACATACTACCTCCTTT
ACTTCACATTCGGGTCGGCATGGCATGTGCACCTCCTCGCGGTCCGACCTGGGCTACTTCGGTAGGCTAAGGCACAGTATGATGTT
GTGAAGTGAAGGAGGATATAAATGAAACAGAACAAAGAACAG

IN_1 ( I{u1d} [d20],[T7t]):

gaagtctaacgctgctctgggctaactgtcTTCTAATACGACTCACTATAGGGAGATTCGTCTCCCAAATACTTAATACAAAATAA
TCACTTCACAACATCAATATAACCCCTTGGGGCCTCTAAACGGGTCTTGAGGGGTTTTTTGctgaaacctcaggcatttgagaagc
acacgctgaaaggaggaactatatccggattggcgactgtgtacttgttataacatctgacagttaaagtcgggagaataggagcc
gcaatacacaatttaccgcatctagacttaactgagatatta
```

In the figures below in this section, $U_i$ indicate homology domains for Gibson assembly. $U_D$ and $U_E$ are 30-base domains derived from the backbone of the pColADuet and pETDuet vectors. $U_G$ is a 30-base domain taken from Ref[14]. $U_X$ and $U_1$ to $U_9$ are 40-base domains taken from Ref[22].

The Thyb terminators are from Ref[23]. They contain a pause site for T7 RNAP that is reported to enhance termination. T7t refers to the terminator sequence from the T7 bacteriophage genome. Readthrough transcription past the terminators was still a concern for constructs with more than one component. For plasmids encoding multiple components we placed ribozyme-based insulators[24] between elements *i.e.,* downstream of the terminator of one element and before the promoter of the next element. These ribozymes should cut readthrough transcripts into the individual RNAs. Insulator sequences came from Ref[25], but we removed the unnecessary 3′ hairpin present in these sequences, hence the ‘n’ at the end of the names in the figures below.

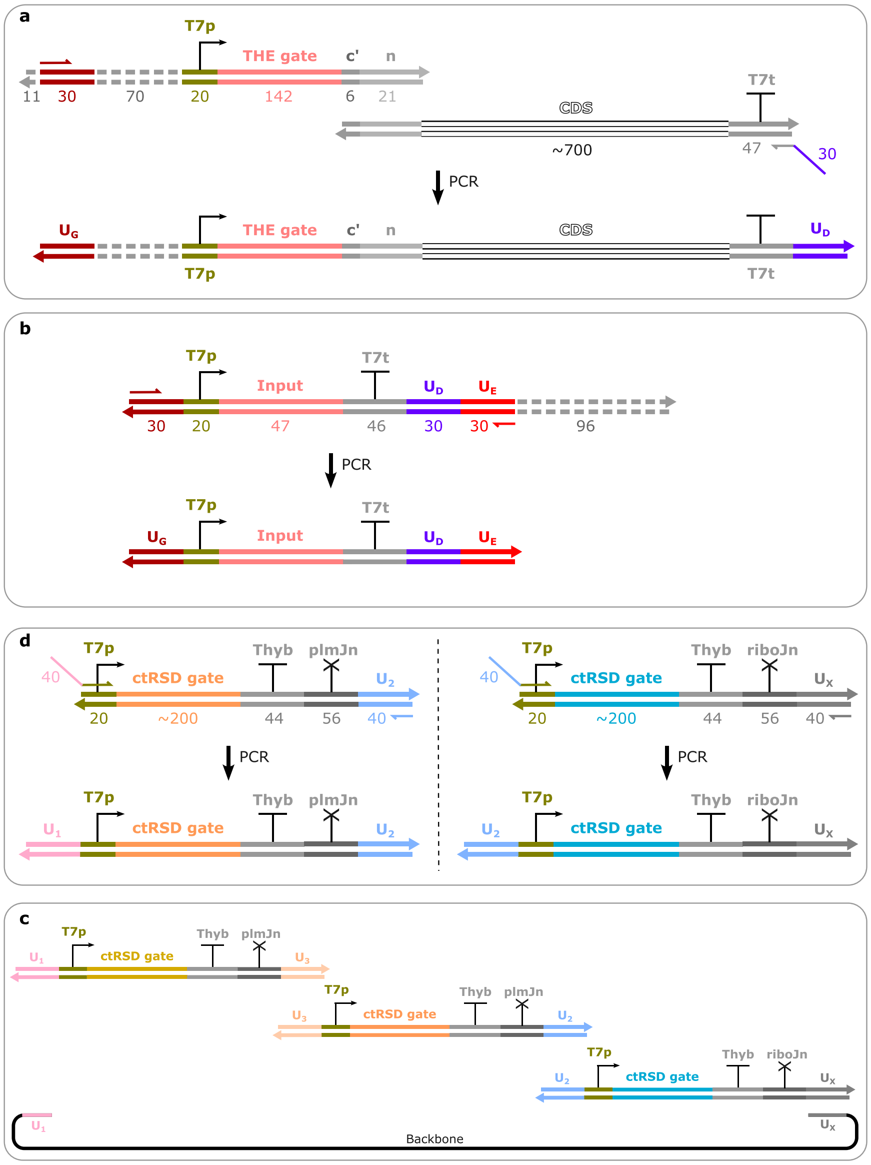

**Supplementary Figure 3**: PCRs and DNA construct assembly strategies. $U_i$ indicate homology domains for Gibson assembly. Dashed gray lines indicate filler sequences appended to get to 300-base minimum sequence length for ordering. (**a**) Schematic of overlap PCR used to assembly THE riboregulators for cloning. (**b**) Schematic of PCRs used for inputs prior to cloning. (**c**) Schematic of PCRs used to assemble upstream gates for multi-layer cascades. (**d**) Schematic of multi-component Gibson assembly. Multi-input constructs were prepared similarly. Homology domain sequences and primers for the depicted PCRs are in Supplementary File S1.

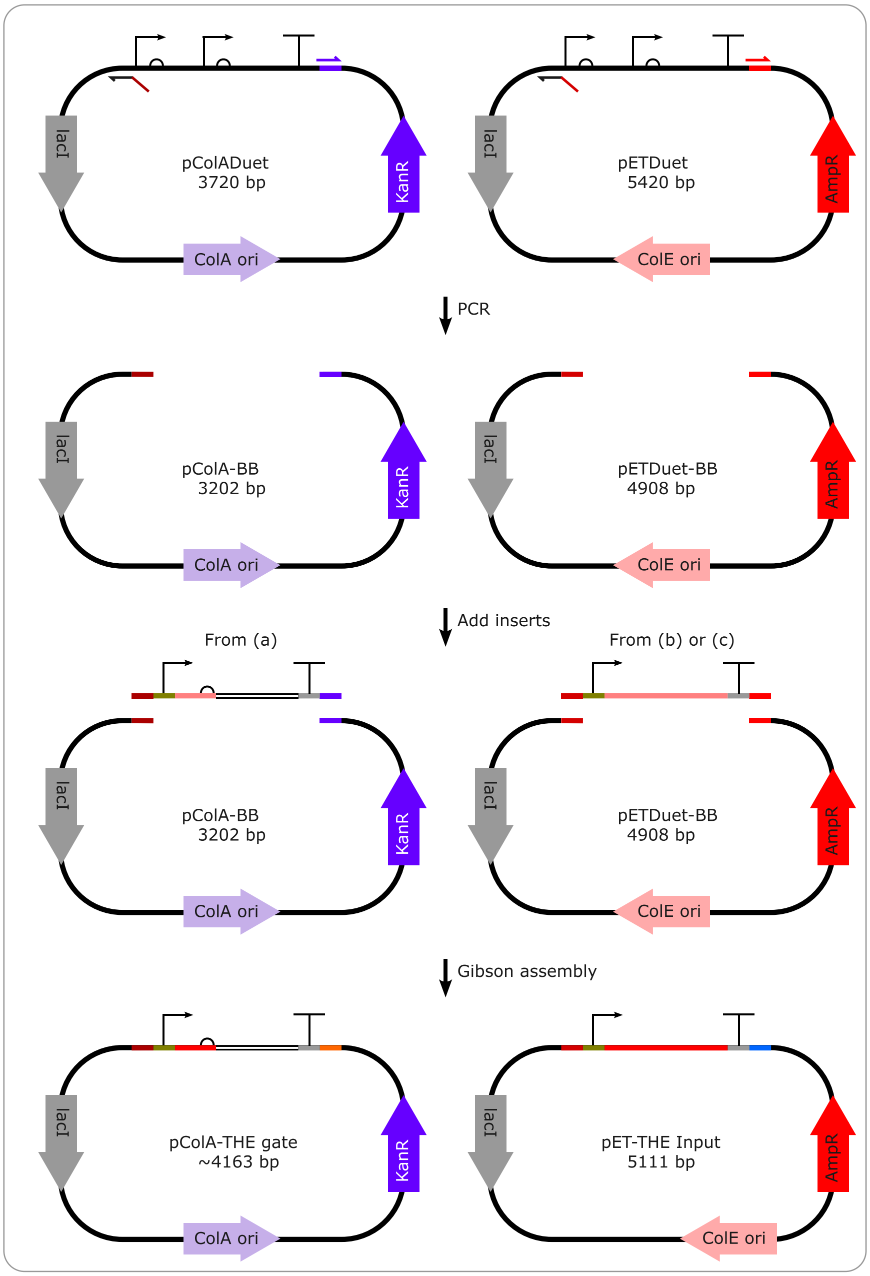

**Supplementary Figure 4**: Schematic of backbone preparation and Gibson assembly of constructs from Supplementary Figure 3. Homology domain sequences and primers for the depicted PCRs are in Supplementary File S1.

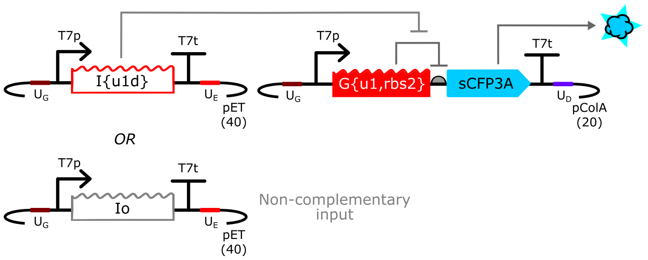

**Supplementary Figure 5**: Plasmid architecture for single input, single THE riboregulator constructs used in Figure 2. Io is a scrambled input that is not complementary to any of the riboregulator input domains. $U_i$ indicate homology domains for Gibson assembly. The number in parentheses below each backbone name indicates putative copy number in *E. coli*.

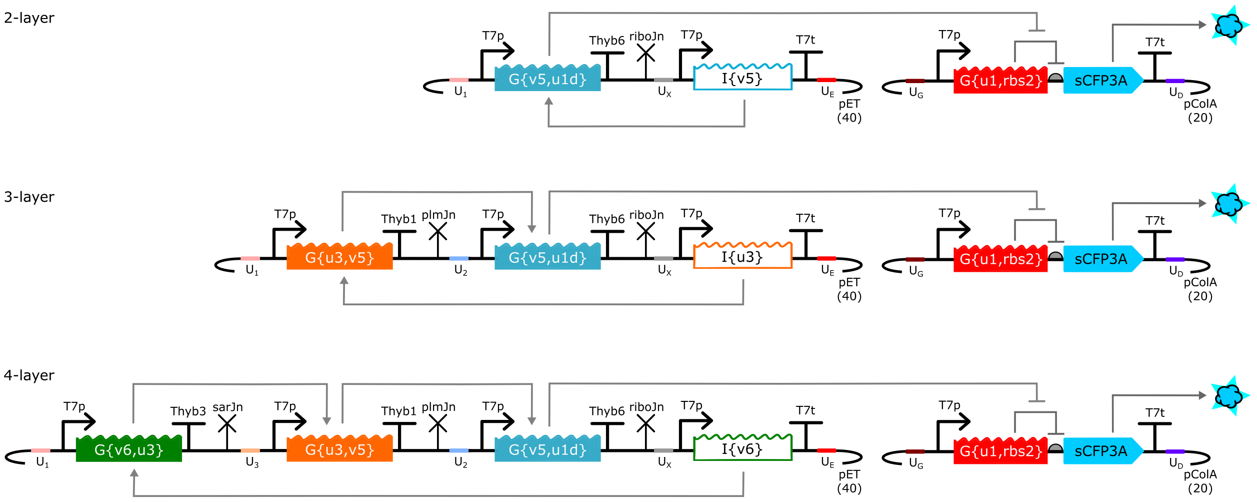

**Supplementary Figure 6**: Plasmid architectures for multi-layer RNA strand exchange constructs used in Figure 3. $U_i$ indicate homology domains for Gibson assembly. The number in parentheses below each backbone name indicates putative copy number in *E. coli*.

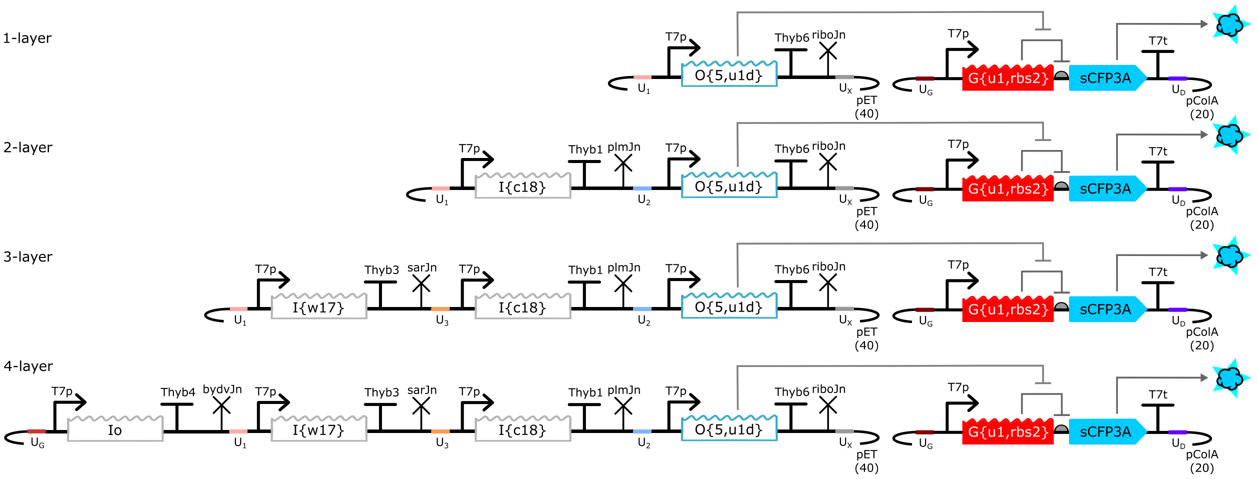

**Supplementary Figure 7**: Plasmid architectures for multi-promoter control constructs for multi-level cascades in Figure 3. The gray inputs Io, I{w17} and I{c18} do not connect to the THE riboregulator and serve as controls for increasing the number of T7 RNAP promoters in the multi-layer cascades influences reporter protein expression levels. $U_i$ indicate homology domains for Gibson assembly. The number in parentheses below each backbone name indicates putative copy number in *E. coli*.

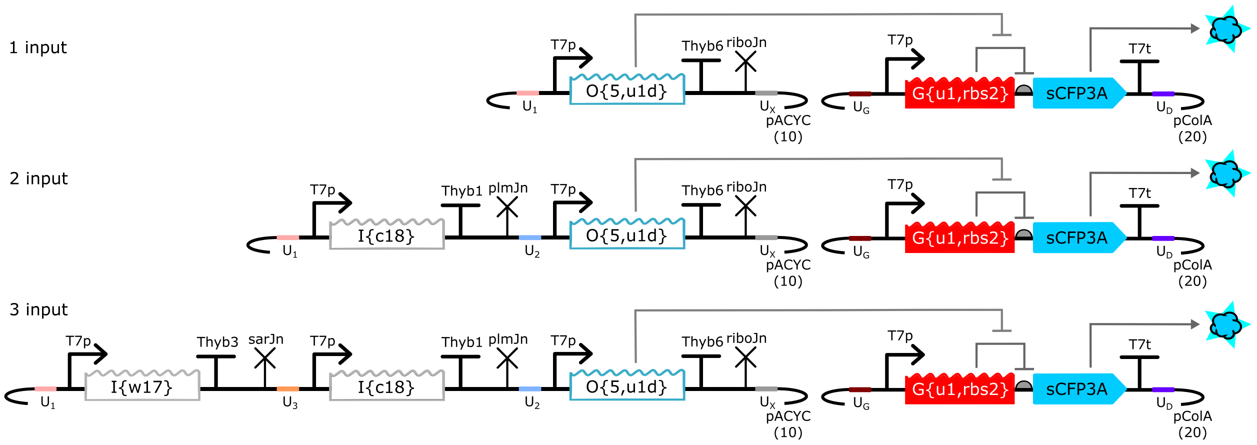

**Supplementary Figure 8**: Plasmid architectures for multi-promoter control constructs for the multi-input classifier circuit in Figure 5 (Supplementary Figure 32). The gray inputs Io, I{w17} and I{c18} do not connect to the THE riboregulator and serve as controls for increasing the number of T7 RNAP promoters in the multi-input logic element influences reporter protein expression levels. $U_i$ indicate homology domains for Gibson assembly. The number in parentheses below each backbone name indicates putative copy number in *E. coli*.

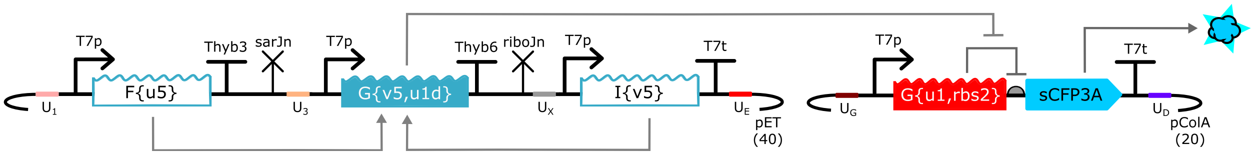

**Supplementary Figure 9**: Plasmid architecture for the fuel amplified cascade tested in Supplementary Figure 31. $U_i$ indicate homology domains for Gibson assembly. The number in parentheses below each backbone name indicates putative copy number in *E. coli*.

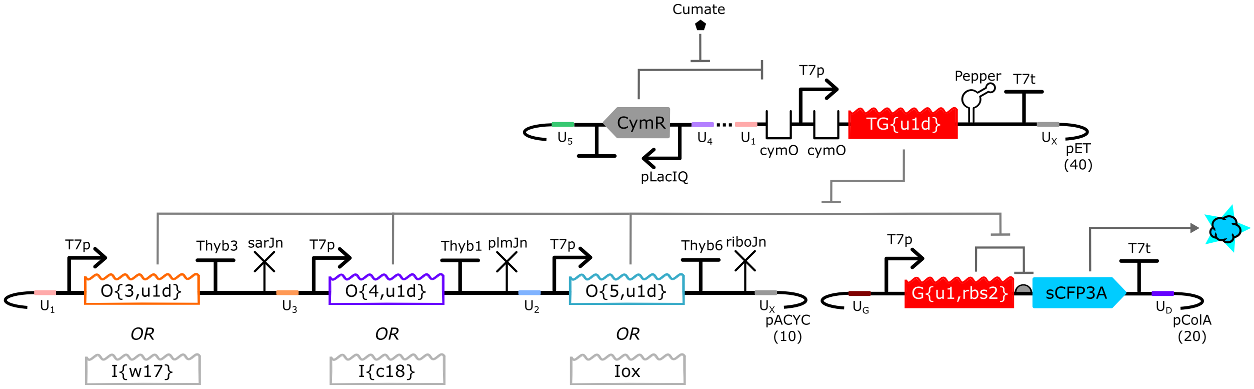

**Supplementary Figure 10**: Plasmid architectures for the thresholding-based multi-input classifier constructs used in Figure 5. Each of the three input positions can either be an input that connects to the THE riboregulator or a scrambled input (I{w17}, I{c18}, or Iox), making eight possible pACYC constructs. $U_i$ indicate homology domains for Gibson assembly. The number in parentheses below each backbone name indicates the putative copy number in *E. coli*. Note the threshold gate contains a Pepper aptamer upstream of its terminator, but fluorescent aptamer measurements were never conducted. The presence of the aptamer could have influenced RNA stability.

#### 4 Cell culturing and flow cytometry

Supplementary Figure 11 illustrates the experimental workflow for cell assays.
Supplementary Figure 12 shows how the time of induction of RNA expression alters cell growth.
Supplementary Figure 13 shows the flow cytometry plate layout and subsequent data analysis.

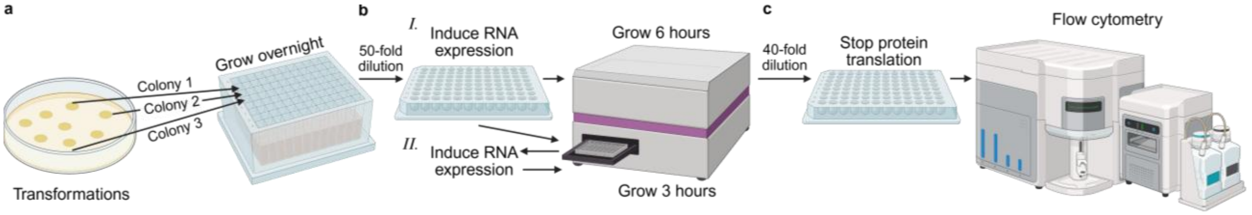

**Supplementary Figure 11**: Experimental workflow for characterizing RNA strand exchange circuits in *E. coli*. The workflow is split into three stages (**a**) Overnight cultures, (**b**) Growth and RNA expression, (**c**) Flow cytometry. As shown in Supplementary Figure 12, inducing T7 RNAP expression right after diluting overnight cultures resulted in toxicity for systems with more than a single input and single gate. So, in this study there were two variations of panel b. The first (I.), where RNA expression was induced alongside the overnight culture dilution, was used for single input-single gate systems and the second (II.), where cells were grown for 3 h after dilution before RNA expression was induced, was used for all other systems, unless otherwise stated. Created with BioRender.

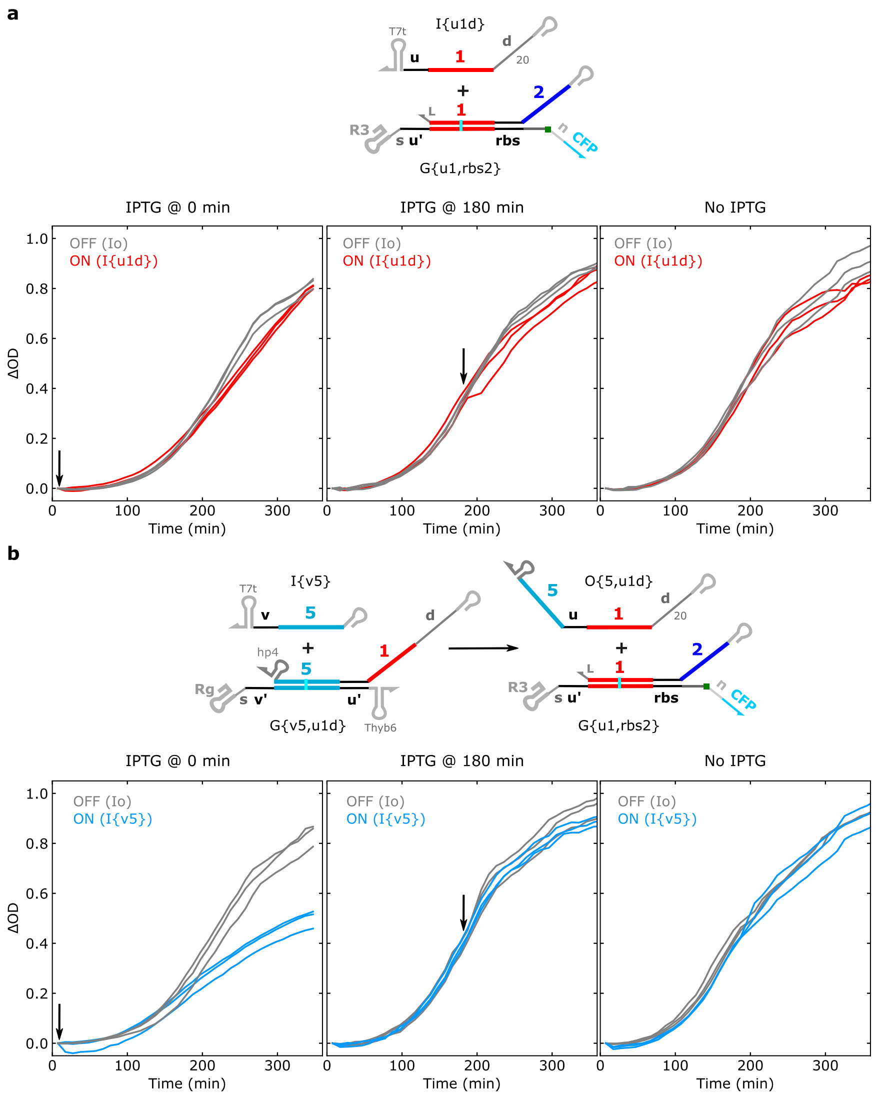

**Supplementary Figure 12**: Growth curves of cells expressing (**a**) 1-layer or (**b**) 2-layer RNA strand exchange systems with IPTG induction either at 0 h (alongside the dilution of overnight cultures) or at 3 h after dilution of overnight cultures. For a 1-layer system, growth is similar for induction of T7 RNAP with 100 µmol/L at 0 h and 3 h and for no T7 RNAP induction panel a. For a 2-layer system, growth is markedly slower for induction of T7 RNAP with 100 µmol/L at 0 h for the ON system panel b. IPTG induction of the 2-layer system at 3 h results in similar growth to conditions without IPTG induction. Unless otherwise stated, experiments with systems containing more than a single input and single gate were induced with IPTG approximately 3 h after overnight culture dilution. Increased toxicity of induction directly after dilution of overnight culture could be due to accumulation of the plasmids in the stationary phase cells[26].

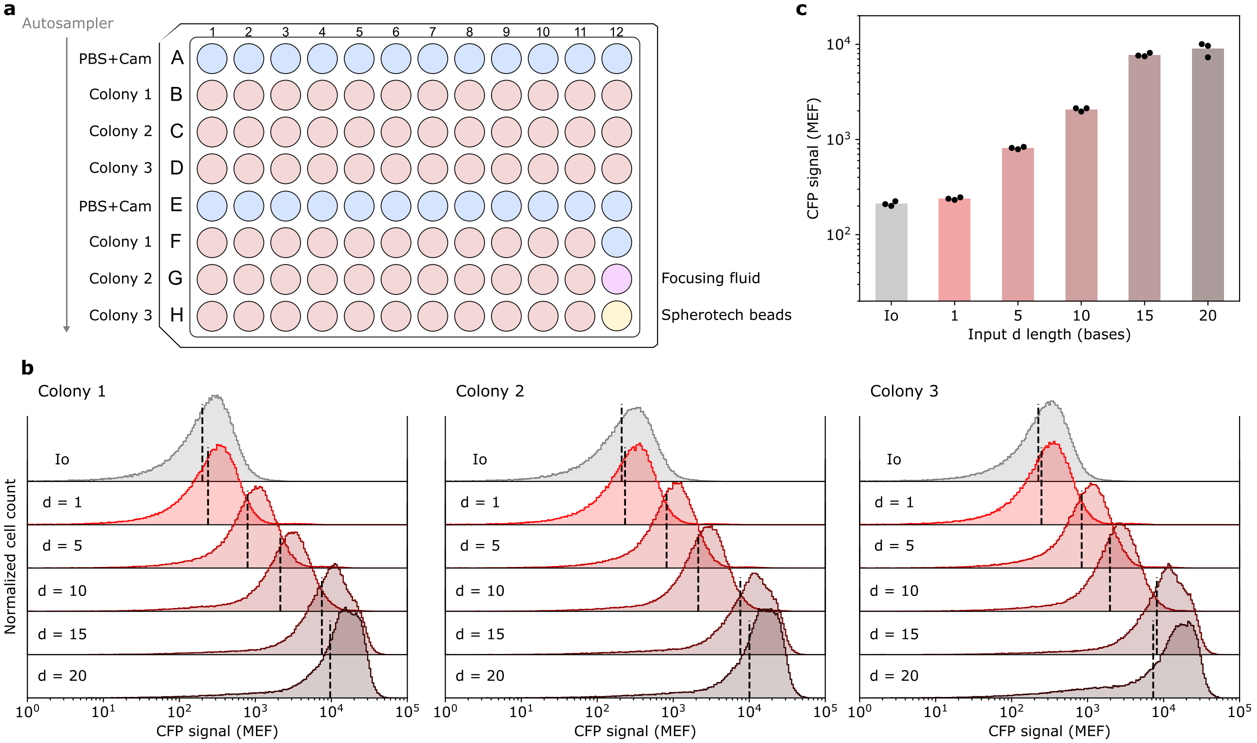

**Supplementary Figure 13**: Overview of flow cytometry analysis. (**a**) Typical plate layout for flow cytometry experiments. Each column contained cells from two different systems each originating from individual transformant colonies. For example, wells B1, C1, D1 are three replicates of one system and wells F1, G1, H1 are three replicates from another system. The samples were read starting at A1 and moving down the rows such that the three replicates of each system were read in succession followed by a PBS wash between replicates. The Spherotech beads in well H12 were used to convert fluorescence signal into molecules of equivalent fluorophore (MEF). (**b**) Flow cytometry distributions for three colonies for the systems shown in Figure 2c of the main text. The vertical dashed black lines represent the geometric means plotted as individual points in the bar plots of panel c. (**c**) Bar plots summarizing CFP signal from the distributions in panel b. Individual data points are geometric means of the flow cytometry distributions in panel b and the height of the bar is average of the geometric means of the three replicates.

#### 5 RT-qPCR measurements of ribozyme cleavage within THE riboregulators

RT-qPCR measurements of ribozyme cleavage were conducted as previously described using a blocking oligonucleotide to prevent ribozyme catalysis during sample preparation and measurement[27]. Using a relative quantification technique typically termed the ΔΔCt method[28–30], the fraction of uncleaved gate for a given sequence was calculated with Eq. 1. The ΔΔCt value is the difference between the ΔCt of U and G for PCRu and the ΔCt of U and G for PCRo (Eq. 2). $\Delta Ct_{PCRu}$ indicates the concentration difference in uncleaved RNA between the U and G samples and $\Delta Ct_{PCRo}$ accounts for any differences in the total concentration of U and G RNA added in an experiment (Supplementary Figure 14).

Eq. 1: $\text{Fraction cleaved} = 1 - 2^{-\Delta\Delta\overline{Ct}}$

Eq. 2: $\Delta\Delta\overline{Ct} = (\overline{Ct}_{U,PCRu} - \overline{Ct}_{G,PCRu}) - (\overline{Ct}_{U,PCRo} - \overline{Ct}_{G,PCRo}) = \Delta\overline{Ct}_{PCRu} - \Delta\overline{Ct}_{PCRo}$

Where $\overline{Ct}$ indicates the mean of three technical replicates.

For the above analysis to be valid, PCR amplification efficiencies should be between 90 % to 110 %[31]. Using dilution series of U control RNAs, we verified the PCR amplification efficiencies for PCRo and PCRu primers ranged between 96 % to 100 % across sequences and replicates from independent RNA extractions (Supplementary Figure 15). Thus, Eq. 1 was valid to use for analysis of ribozyme cleavage (Supplementary Figure 16).

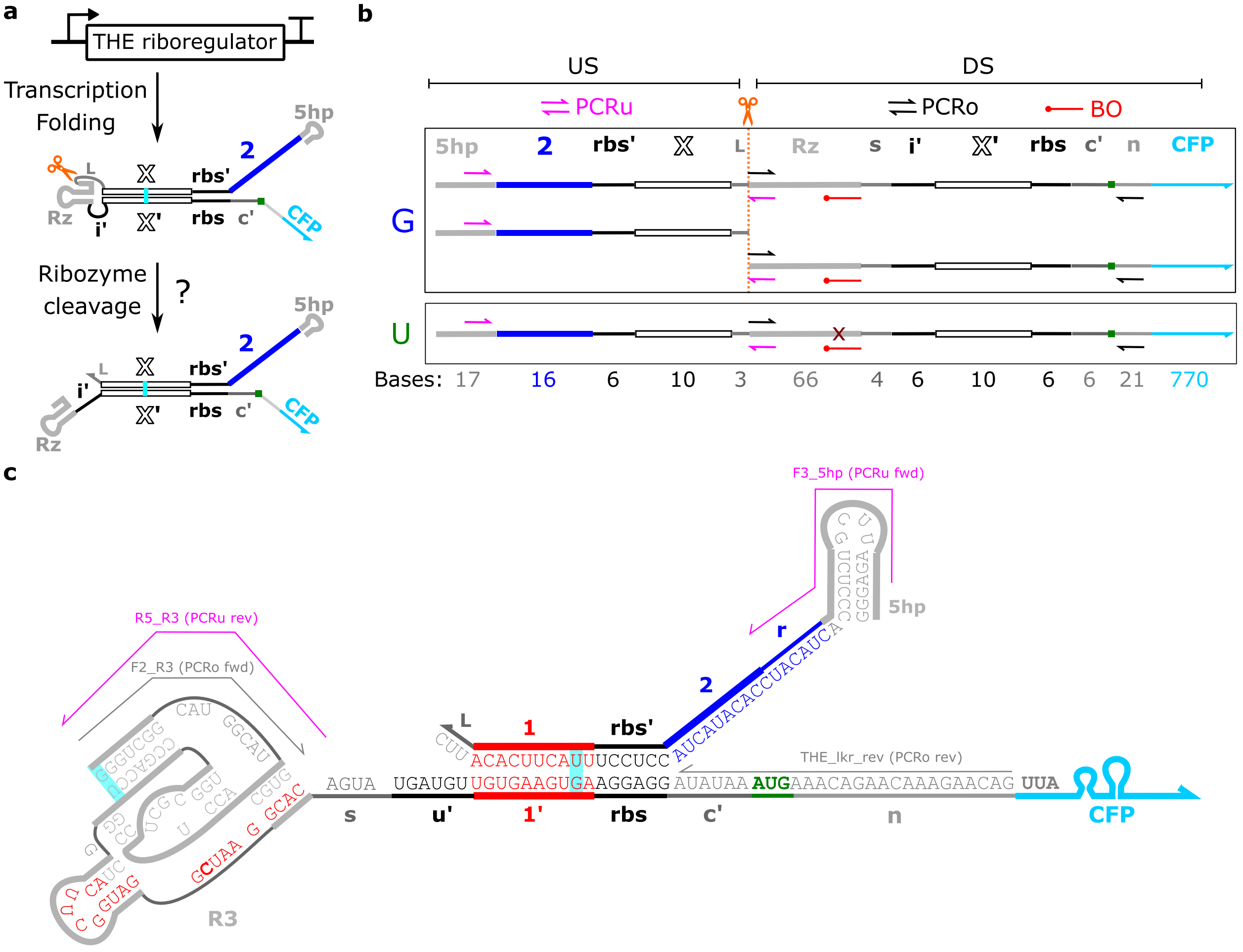

**Supplementary Figure 14**: Overview of primer design for RT-qPCR measurements of ribozyme cleavage. Schematics and caption adopted from Ref[27].(**a**) Schematic of genetically encoded THE riboregulator. Domains X and domains i′, X′ represent different sequences upstream and downstream, respectively, of the ribozyme. (**b**) Schematic of primer layout to measure ribozyme cleavage with RT-qPCR. The PCRu primers (pink) span the cleavage site (orange dashed line) and should only amplify uncleaved products. The PCRo primers amplify both cleaved and uncleaved products to amplify total RNA. U is a reference RNA which contains a single base mutation (maroon X) compared to G, which abolishes ribozyme activity. The difference in PCRu Ct between G and U transcripts, corrected for total RNA differences by PCRo, yields a measure of ribozyme cleavage (Methods). BO represents a blocking oligonucleotide that is added to prevent ribozyme cleavage during sample preparation. The red circle at the left end of BO indicates a 3′ amino modification to prevent extension during RT-qPCR. The numbers below the box indicate the length of each domain in bases. (**c**) Gate secondary structure schematics. Red bases in the ribozyme sequence indicate the sequence the blocking oligonucleotide binds to within the ribozyme. The bold C base in the ribozyme sequence indicates the base that was changed to a U base to abolish cleavage in the uncleaved control RNAs. Pink and gray lines indicate the regions the PCRu and PCRo primers bind to, respectively. Cyan highlighted bases indicate GU wobble base pairs. Primer and blocking oligonucleotide sequences are in Supplementary File 1.

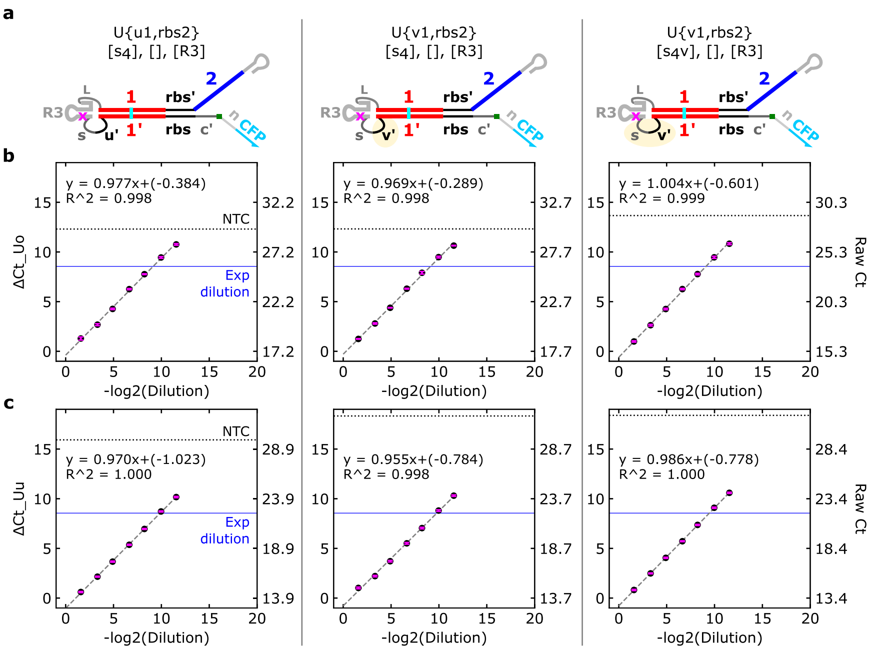

**Supplementary Figure 15**: Primer amplification efficiency analysis. (**a**) Schematics of uncleaved gates used to assess primer amplification efficiency. (**b**,**c**) RT-qPCR results for a 3000-fold dilution series of uncleaved gates shown in panel a. The ΔCt (relative to RNA at approximately 5 ng/μL) and corresponding Ct values are labeled on the right and left axes, respectively. Panel b shows results for the PCRo primer set and panel c shows results for the PRCu primer set. Pink error bars indicate ± one standard deviation of the mean from three technical replicates. Both primer sets showed > 95 % amplification efficiency (100 × slope of fit lines) across the three tested sequences. The dotted black horizontal lines represent controls in which the RNA was left out of the reactions (NTC), which indicates the upper bound of measurable Ct. The blue horizontal lines indicate the dilution used in the final experimental protocol, which is in the linear range of these control curves and multiple Ct away from NTC. For the dilution series, a sample of U RNA nominally at 5 ng/μL were serial diluted 3-fold, 10-fold, 30-fold, 100-fold, 303-fold, 1000-fold, and 3030-fold in RNA storage solution.

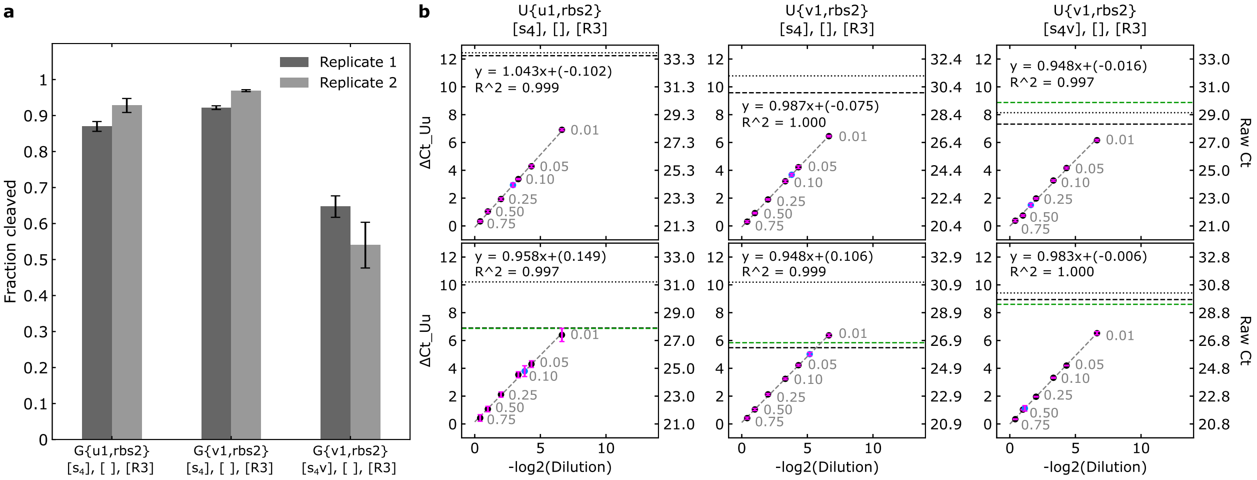

**Supplementary Figure 16**: Results of RT-qPCR measurements of ribozyme cleavage. (**a**) Fraction cleaved RNA of the gate sequences labeled on the x-axis. Error bars indicate standard deviation from three technical replicates. Replicate 1 and replicate 2 indicate replicates from two independent cell cultures and RNA purifications conducted on different days. (**b**) Amplification efficiency curves and controls for each sample. Top row are results from replicate 1 and bottom row are results from replicate 2. The black dots represent dilutions of the uncleaved (U) control RNA sample and the blue dot indicates the gate (G) RNA sample. Pink error bars indicate ± one standard deviation of the mean from three technical replicates. PCRu primer sets showed ≥ 95 % amplification efficiency (100 × slope of fit lines) across the three tested sequences over a narrower dilution range than in Supplementary Figure 15. The dotted black horizontal lines represent controls in which the RNA was left out of the reactions (NTC), which indicates the upper bound of measurable Ct. Dashed black and green horizontal lines indicate controls in which the reverse transcriptase enzyme was left out (NRTC), with green indicating the U RNA extraction sample and black indicating the G RNA extraction sample. The measured ΔCt for the gate RNA is typically much lower than the negative controls. For the dilution series, a sample of U RNA nominally at 14 pg/μL were diluted 1.33-fold (0.75) to 100-fold (0.01) in RNA storage solution.

### 6 Additional THE riboregulator design space experiments

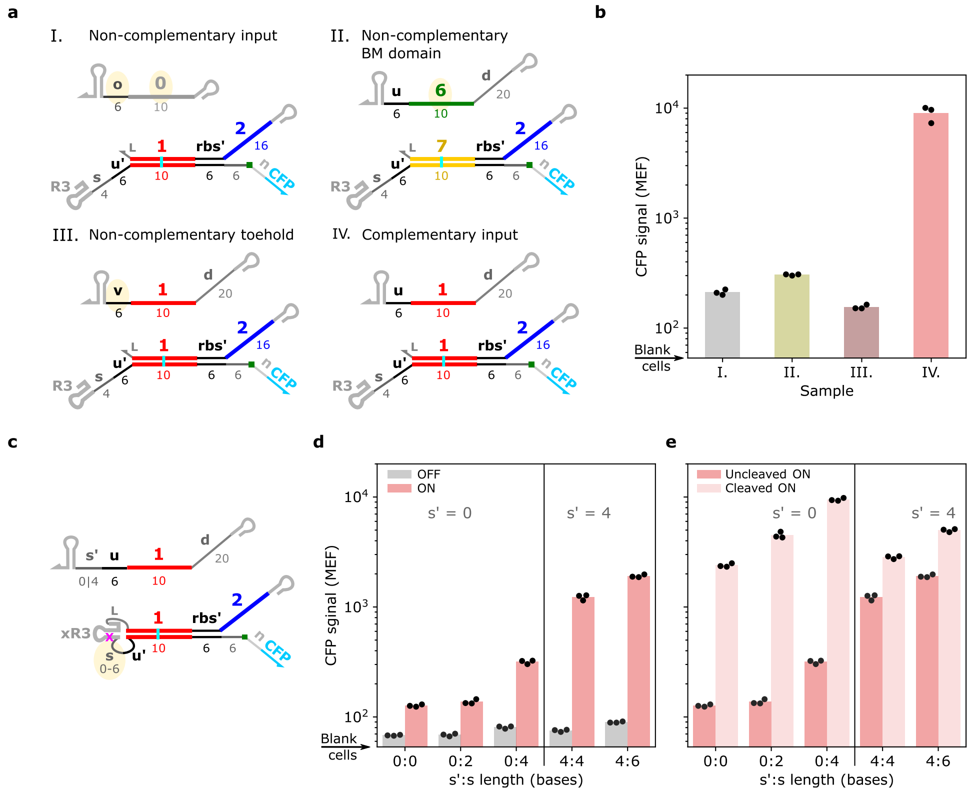

**Supplementary Figure 17**: **Mechanistic controls for toehold exchange riboregulators in cells.** (**a**,**b**) Toehold exchange riboregulators are sequence specific. (**a**) Schematics of constructs used to test sequence specificity of THE riboregulators in *E. coli*. $s=s_4$ for all gates. In I., both the toehold and the branch migration (BM) domain of the input strand are not complementary to the gate. In II., the input strand toehold is complementary to gate, but the branch migration domain is not. In III., the input strand toehold is not complementary to the gate, but the branch migration domain is. In sample IV., the toehold and branch migration domains of the input are complementary to the gate, enabling strand exchange. (**b**) Geometric means of flow cytometry distributions for components depicted in panel a. Only when both the toehold and branch migration domains of the input are complementary to the gate is substantial protein expression observed. (**c-e**) **Ribozyme cleavage is necessary for substantial protein expression with the canonical toehold exchange** riboregulator design ($s'{:}s$ = 0:4). (**c**) Schematic of the uncleaved ribozyme constructs tested. xR3 has a single mutation relative to R3 that abolishes cleavage activity[1]. The length of the spacer ($s$) domain was varied within the gate ($s = s_X u$, where x is the length in bases) and tested with input strands with either 6 bases ($s'$ = 0) or 10 bases ($s'$ = 4) of complementarity with the gate toehold. (**d**) Characterization of uncleaved constructs in panel a with complementary (ON) or non-complementary (OFF) inputs and (**e**) a comparison of the uncleaved vs cleaved riboregulator gates with complementary inputs. Increasing the spacer length of the gate and the complementarity of the input toehold to the gate toehold increases protein production for the uncleaved gates, likely from loop-mediated invasion of the hairpin[32]. These results support that ribozyme cleavage is occurring in the canonical THE riboregulator design.

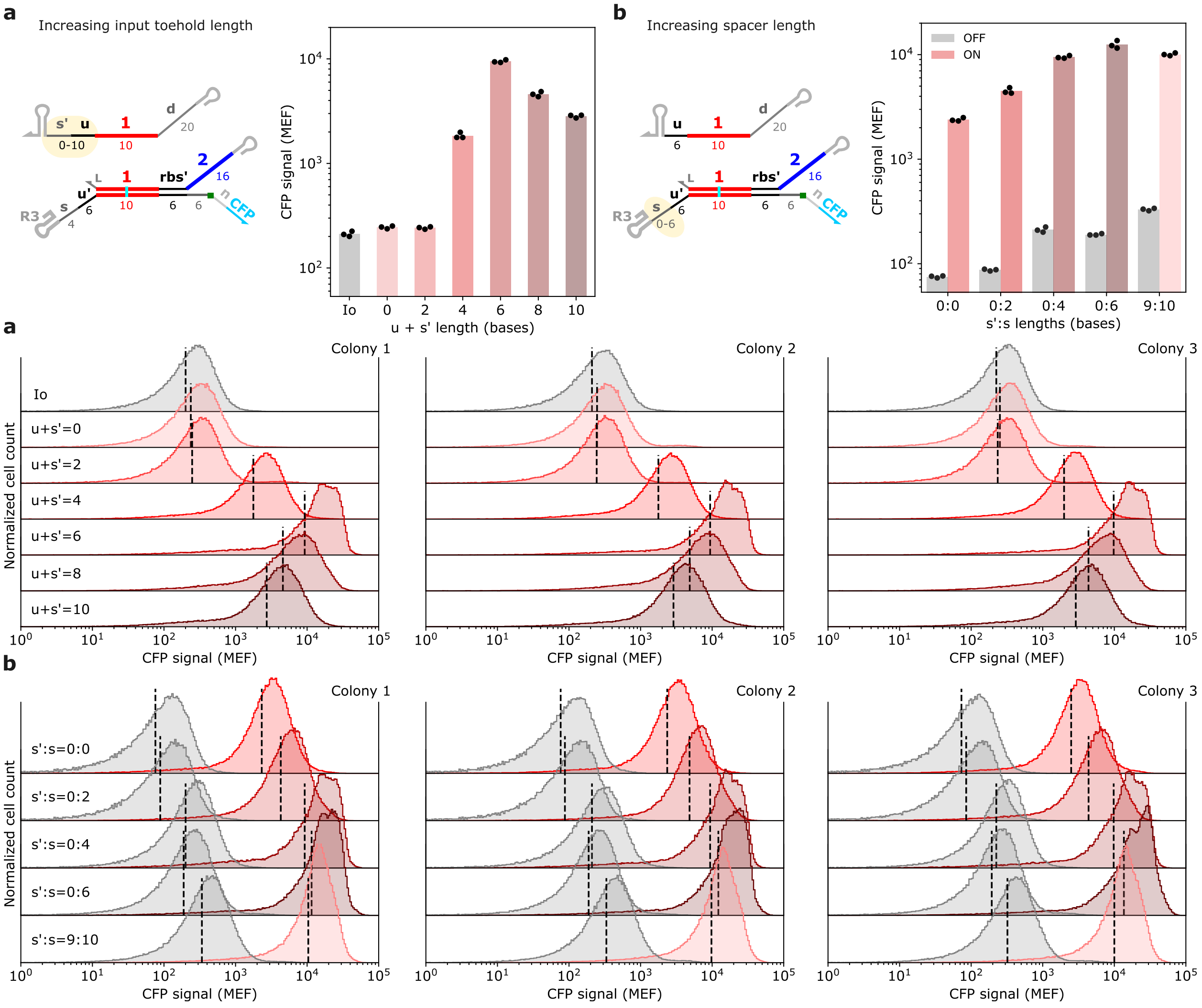

**Supplementary Figure 18: (a,b)** Flow cytometry data for THE riboregulators with different inputs and spacer lengths. Dashed black lines in cell distributions represent the geometric means plotted in the bar plots. Bar plots are reproduced from Figure 2e,f.

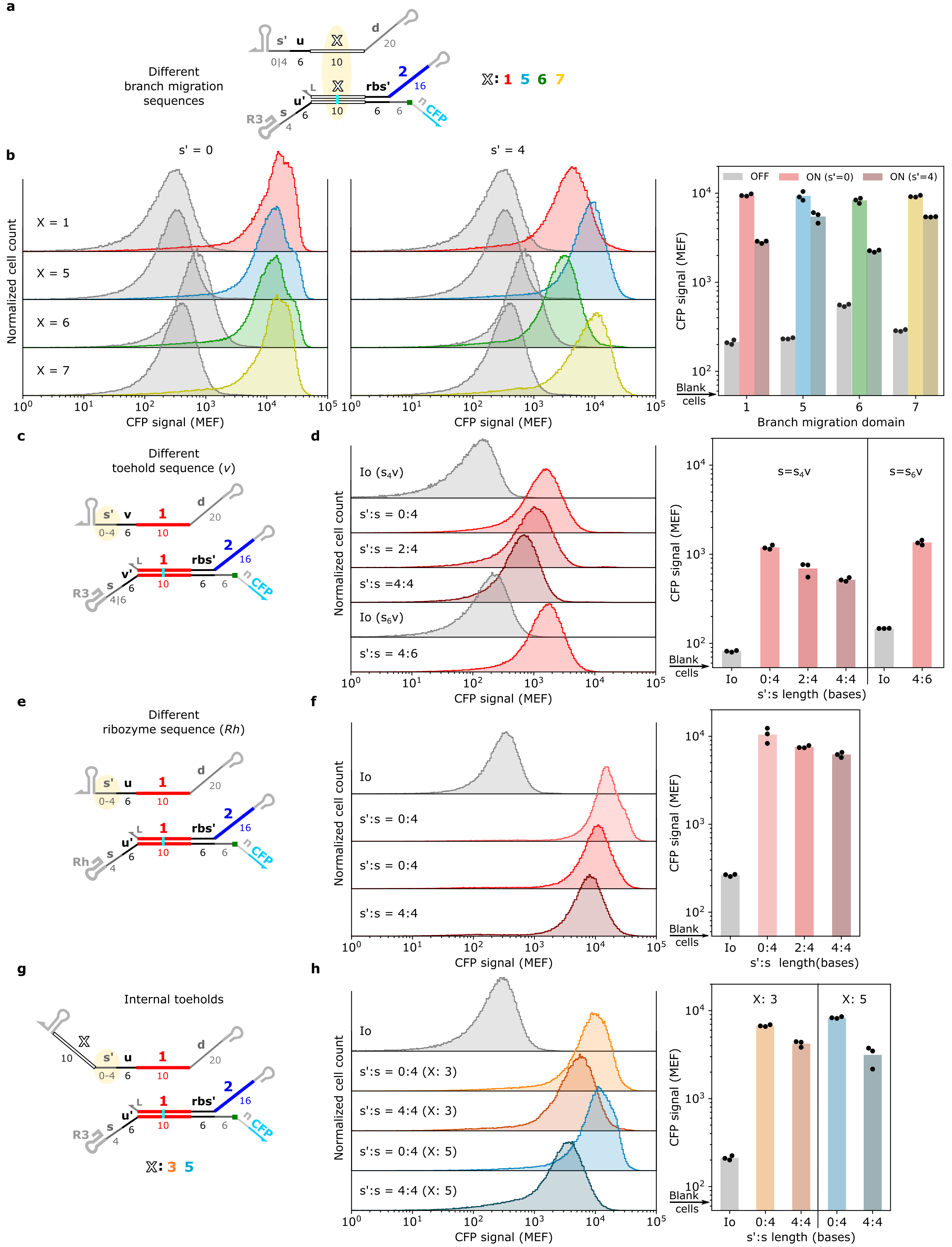
a
Different branch migration sequences
X: 1 5 6 7
b
s' = 0
s' = 4
Normalized cell count
X = 1
X = 5
X = 6
X = 7
CFP signal (MEF)
OFF
ON (s'=0)
ON (s'=4)
Blank cells
Branch migration domain
c
Different toehold sequence (v)
d
Io (s4v)
s':s = 0:4
s':s = 2:4
s':s =4:4
Io (s6v)
s':s = 4:6
s':s length (bases)
e
Different ribozyme sequence (Rh)
f
Io
s':s = 0:4
s':s = 4:4
g
Internal toeholds
X: 3 5
h
s':s = 0:4 (X: 3)
s':s = 4:4 (X: 3)
s':s = 0:4 (X: 5)
s':s = 4:4 (X: 5)

**Supplementary Figure 19**: **RNA inputs with toeholds longer than six bases result in less protein expression across different branch migration and toehold sequences**. (**a**,**c**,**e**,**g**) Input and gate schematics with domain variations highlighted. In panel a, the *s* domain on the gate is $s_4u$ and either 6-base (s′=0) or 10-base (s′=4) toeholds were used on the inputs. (**b**,**d**,**f**,**h**) Flow cytometry results for input and gate combinations shown in in panels a, c, e, g, respectively. The trend of 10-base toeholds producing lower signal than 6-base toeholds was also observed for inputs and gates with different terminator sequences (Supplementary Figure 40).

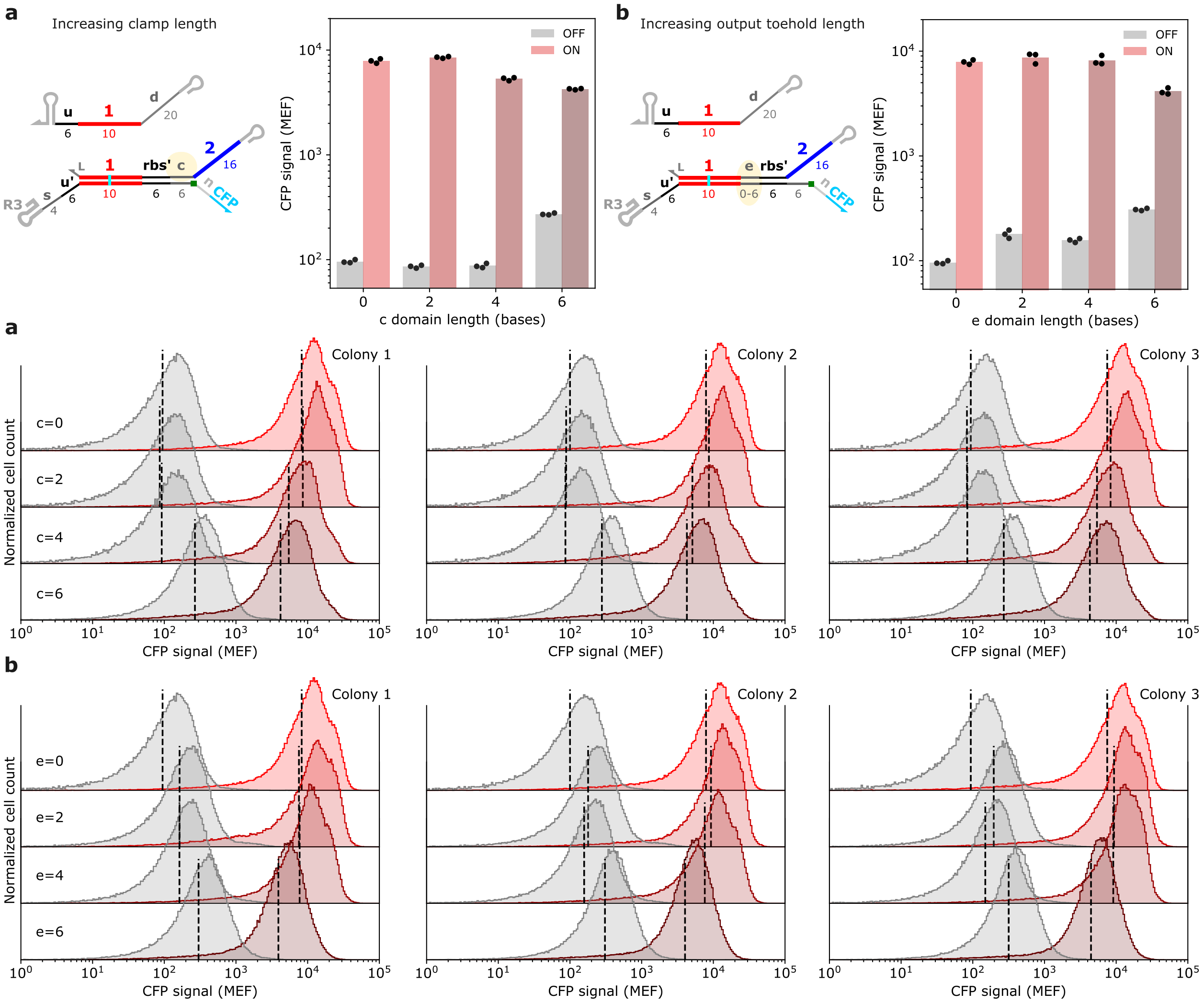

**Supplementary Figure 20:** (**a,b**) Flow cytometry data for THE riboregulators with different output toehold lengths. Vertical dashed black lines in cell distributions represent the geometric means plotted in the bar plots. Bar plots are reproduced from Figure 2g and Supplementary Figure 21, respectively.

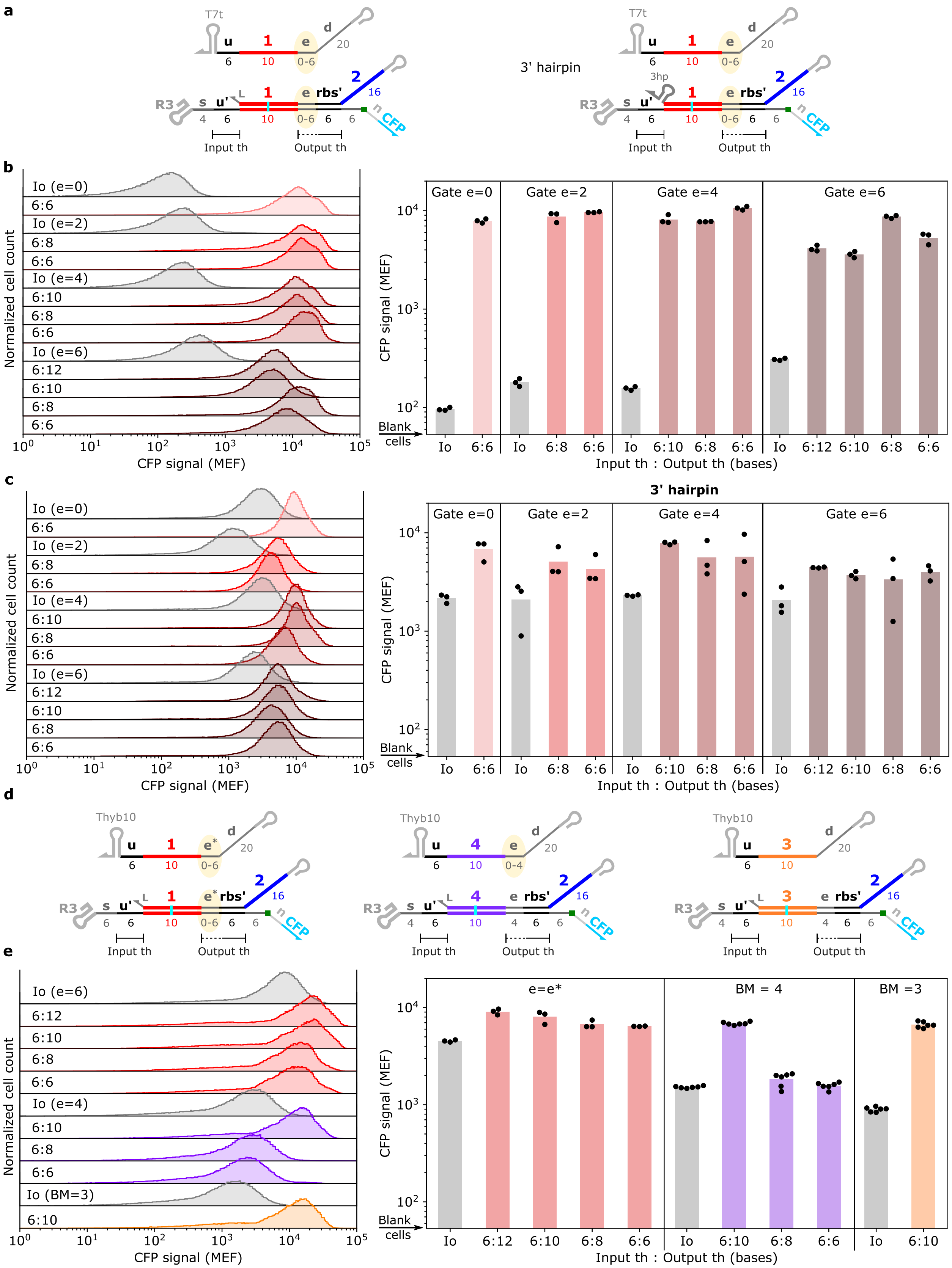
a
T7t
u
1
e
d
20
6
10
0-6
R3
s
u'
L
1
e
rbs'
2
16
4
6
10
0-6
6
6
CFP
Input th
Output th
3' hairpin
3hp
b
Io (e=0)
6:6
Io (e=2)
6:8
6:6
Io (e=4)
6:10
6:8
6:6
Io (e=6)
6:12
6:10
6:8
6:6
Normalized cell count
CFP signal (MEF)
Gate e=0
Gate e=2
Gate e=4
Gate e=6
CFP signal (MEF)
Blank cells
Io
Input th : Output th (bases)
c
3' hairpin
d
Thyb10
e*
4
0-4
3
e
Io (e=6)
6:12
6:10
6:8
6:6
Io (e=4)
6:10
6:8
6:6
Io (BM=3)
6:10
e=e*
BM = 4
BM =3

**Supplementary Figure 21**: **Extending the output toehold relative to the input toehold on a THE riboregulator does not substantially alter protein expression in cells.** (**a**) Schematics of THE riboregulators with longer output toeholds. Output toeholds were extended by an *e* domain upstream of the *rbs*. The input strands tested all had 6-base toeholds and various input/output toehold ratios were obtained by using inputs with varying degrees of complementarity with the *e* domain on the gate. Note that gates activated by inputs with shorter *e* domains will have less secondary structure upstream of the *rbs* and could thus translate more efficiently. Gates without and with hairpins used the $s_4$ and $s_4u$ spacer sequences, respectively. (**b**,**c**) Flow cytometry results for the designs in panel a with representative distributions shown on the left and geometric means of distributions on the right. For both designs, increasing the length of the output toehold relative to the input toehold did not substantially change protein expression, even with a 6-base imbalance. (**d**) Schematics of THE riboregulators with longer output toeholds using an alternative *e* domain (*e**) or different input branch migration domains (*3* or *4*). The gate with the *e** domain used the $s_6u$ spacer and the other two gates use the $s_4$ spacer sequence. (**e**) Flow cytometry results of the components in panel d. The insensitivity to expression levels for longer output toeholds was also observed for gates with different 3 untranslated regions (Supplementary Figure 40).

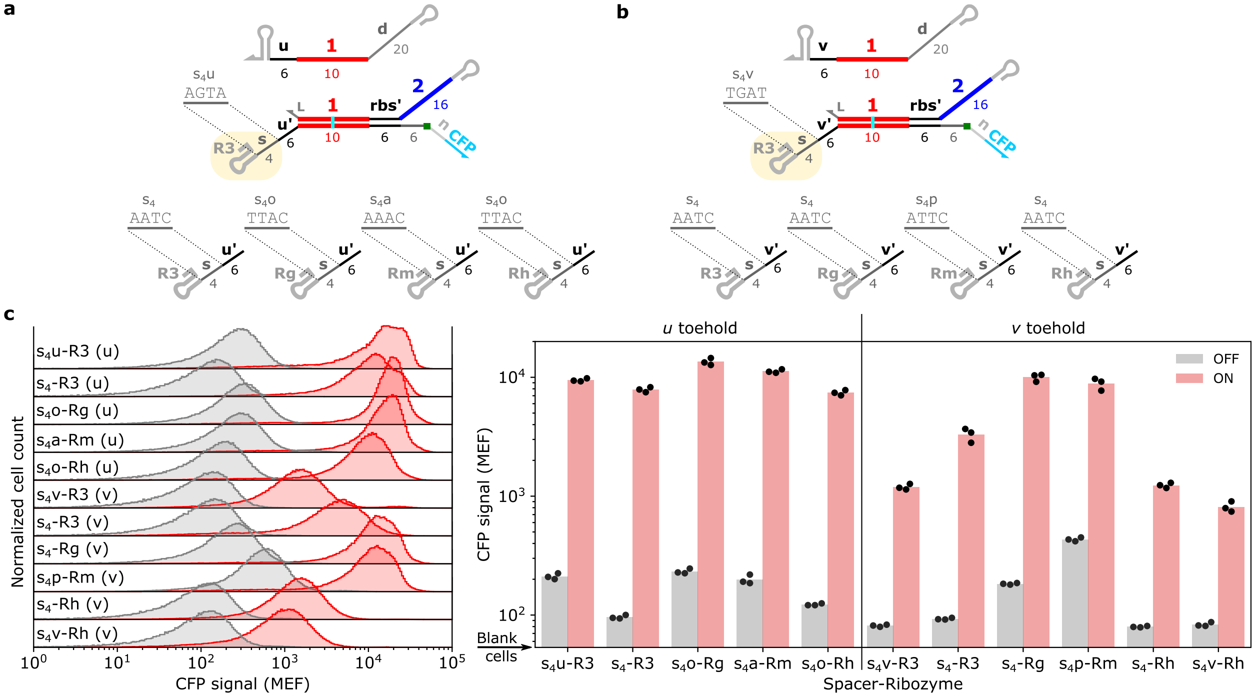

**Supplementary Figure 22**: **Orthogonal ribozymes and toeholds are functional in cells.** Schematics of THE riboregulators with different HDV-like ribozymes and spacer sequences and either (**a**) the *u* input toehold or (**b**) the *v* input toehold. The *u* and *v* toeholds and ribozymes sequences were previously characterized *in vitro* as part of the ctRSD toolkit[2]. The spacer (*s*) domains were designed with slightly different sequences depending on the ribozyme and input toehold combination to prevent undesired secondary structure predicted by NUPACK[8]. Spacer domains $s_4u$ and $s_4v$ indicate spacers designed to have additional complementarity with $s'$ domains on the inputs. All other spacer domains ($s_4x$) are designed with a C base adjacent to the toehold to prevent any additional pairing with inputs or outputs from upstream gates. (**c**) Flow cytometry results for the designs shown in panel a and panel b. The lower signal for $s_4v$-*R3* appears to be due lower ribozyme cleavage efficiency (Supplementary Section 5). This is also likely the reason for the low signal with the *Rh* ribozyme with the *v* toeholds, as both spacers had undesired predicted secondary structure.

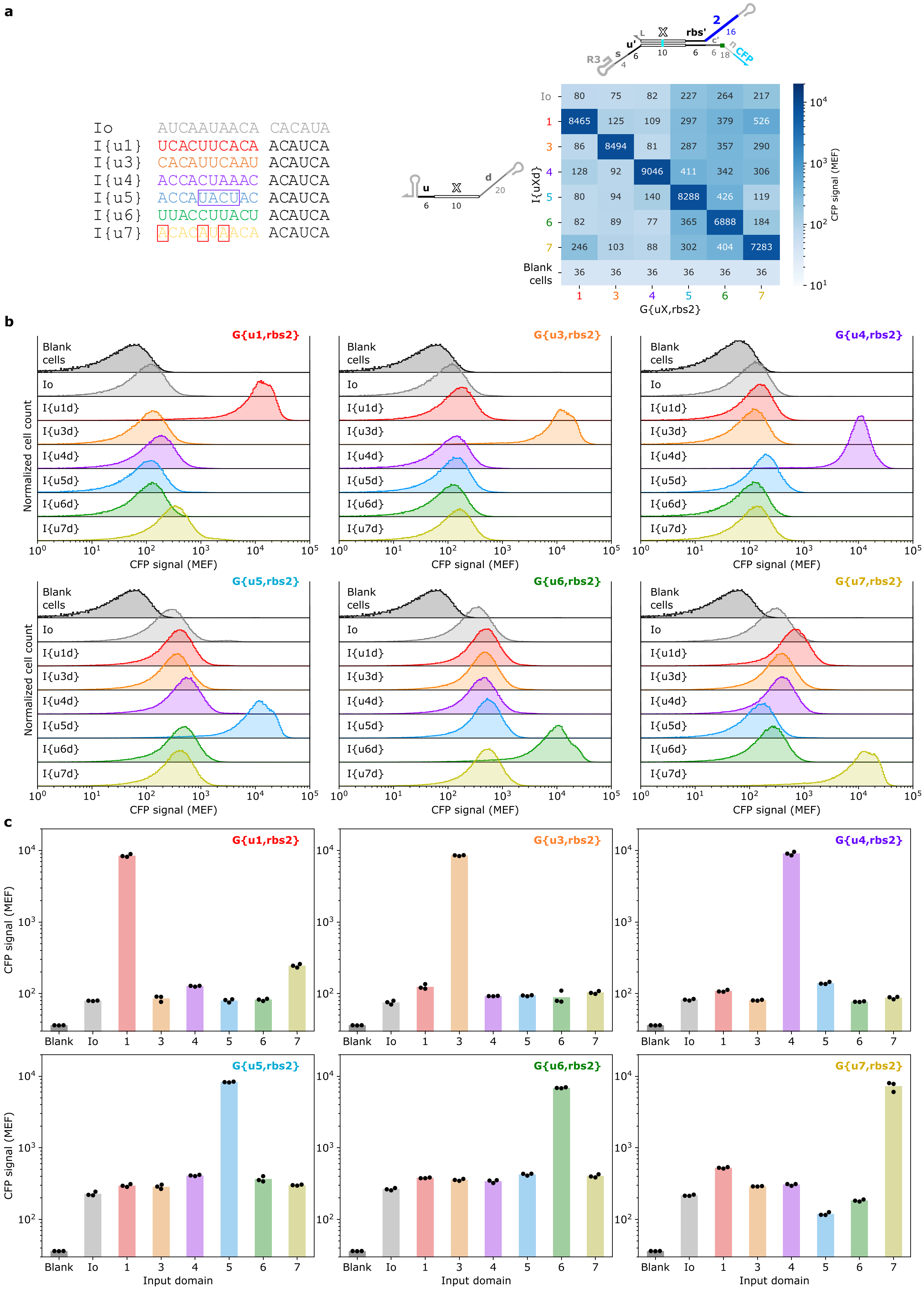
a
Io AUCAAUAACA CACAUA
I{u1} UCACUUCACA ACAUCA
I{u3} CACAUUCAAU ACAUCA
I{u4} ACCACUAAAC ACAUCA
I{u5} ACCAUACUAC ACAUCA
I{u6} UUACCUUACU ACAUCA
I{u7} ACACAUAACA ACAUCA
u X d
6 10 20
R3 s u' L X rbs' 2 c' CFP
4 6 10 6 16 6 18
I{uXd}
Io 80 75 82 227 264 217
1 8465 125 109 297 379 526
3 86 8494 81 287 357 290
4 128 92 9046 411 342 306
5 80 94 140 8288 426 119
6 82 89 77 365 6888 184
7 246 103 88 302 404 7283
Blank cells 36 36 36 36 36 36
1 3 4 5 6 7
G{uX,rbs2}
CFP signal (MEF)
b
G{u1,rbs2}
G{u3,rbs2}
G{u4,rbs2}
G{u5,rbs2}
G{u6,rbs2}
G{u7,rbs2}
Blank cells
Io
I{u1d}
I{u3d}
I{u4d}
I{u5d}
I{u6d}
I{u7d}
Normalized cell count
CFP signal (MEF)
c
CFP signal (MEF)
Blank Io 1 3 4 5 6 7
Input domain

**Supplementary Figure 23**: **Detailed orthogonal input domain results from Figure 3 of the main text.** (**a**) Left: The branch migration and toehold domain sequences of the tested inputs, shown 5′ to 3′. The colored boxes over bases highlight differences between select inputs, with the color of the box matching the color of the input being compared. For example, I{u7} differs by just three bases from I{u1} (red boxes). Right: The heatmap from Figure 3 with mean values from panel c. (**b**) Flow cytometry distributions for the samples in the panel a heatmap. (**c**) Geometric means of flow cytometry distributions in panel b. Each dot represents an independent colony.

#### 7 Testing THE riboregulators with mismatches introduced during strand exchange

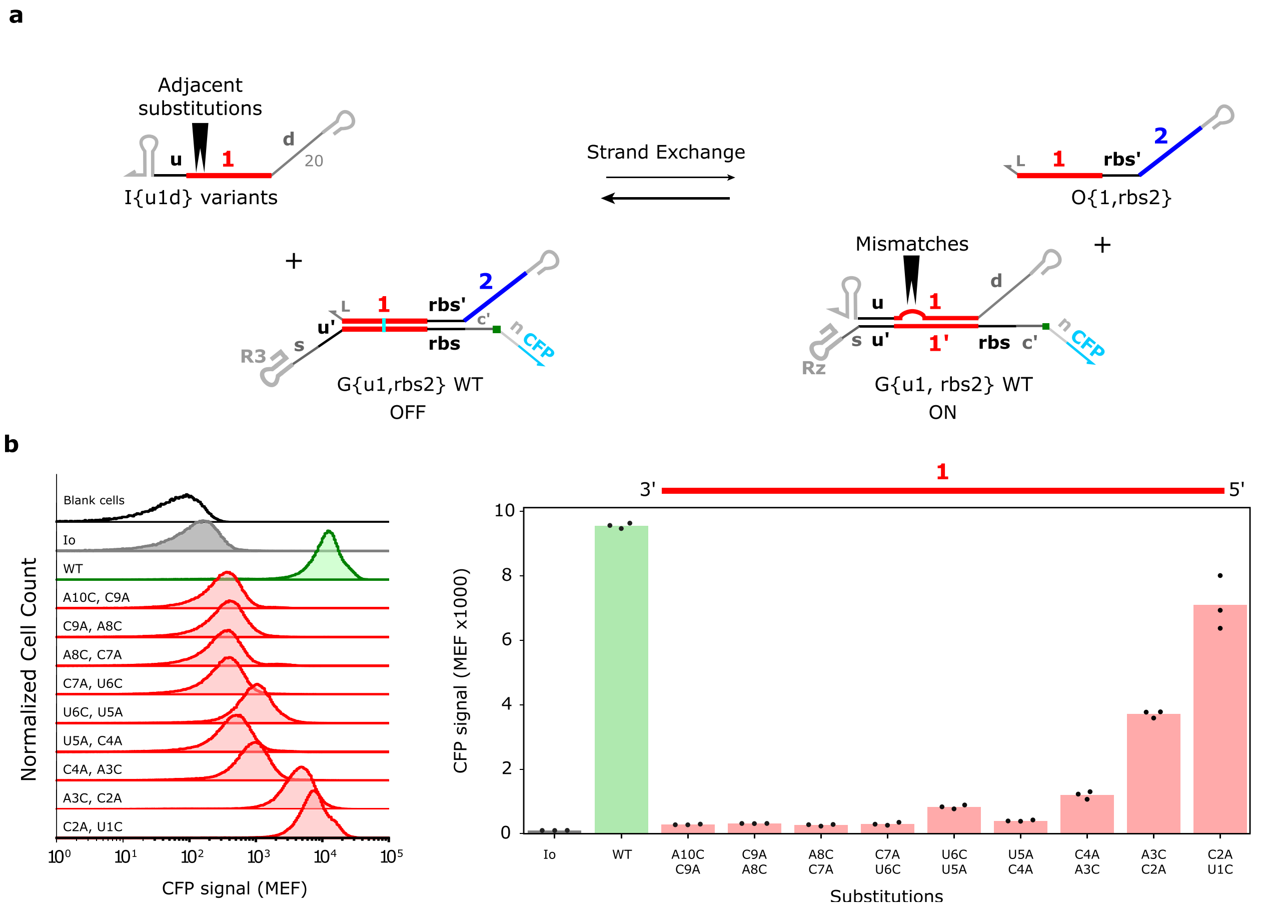

**Supplementary Figure 24**: **Using inputs that introduce two adjacent mismatches with the wild-type THE riboregulator.** (**a**) Schematic depicting an input in which two adjacent nucleotides in the branch migration domain of I{u1d} have been substituted with different nucleotides, resulting in two adjacent mismatches with the corresponding THE riboregulator. (**b**) Flow cytometry measurements of inputs with two mutations activating G{u1, rbs2}. Mutations are numbered 5′ to 3′ based on their position in the branch migration (BM) domain of the input RNA. Left: histograms showing distribution of flow cytometry values for a single representative measurement for each input. Right: Bars indicate the mean CFP signal from n = 3 biological replicates shown as dots. Io: non-complementary (decoy) input, WT: wild-type (*i.e.*, canonical I{u1d} input sequence).

As expected, substitutions introduced near the 3′ end of the input lead to dramatic decreases in CFP signal relative to WT, reflecting poor THE riboregulator activation. However, mutations introduced near the 5′ end of the input (positions 1-4) produce increasingly large CFP signals, with adjacent mismatches at the extreme 5′ end (positions 1 and 2) producing a signal that is approximately 74 % of the WT.

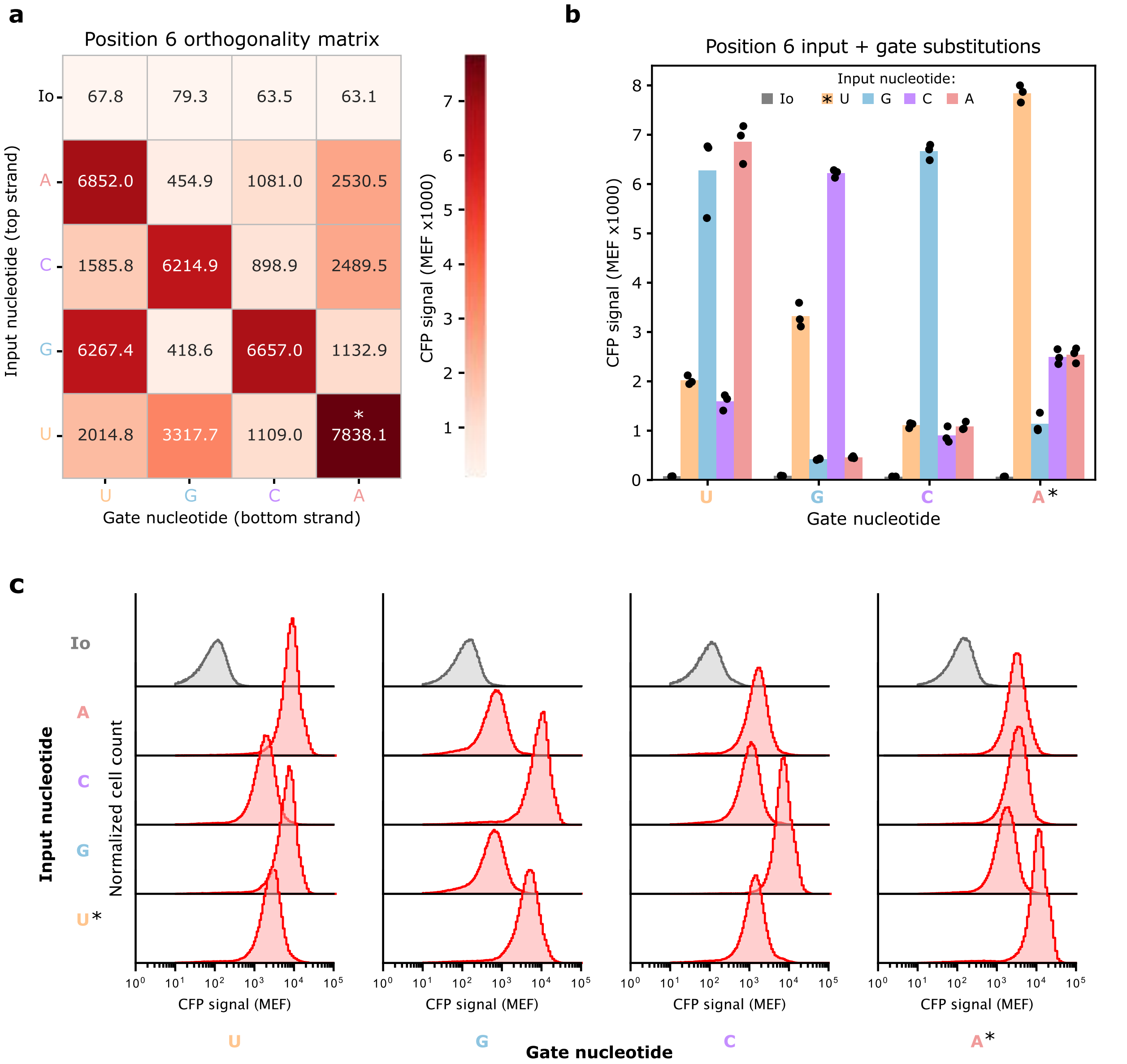

**Supplementary Figure 25**: **Detailed single nucleotide mutation results from Figure 3 from main text.** (**a**) Orthogonality matrix showing flow cytometry results of input + gate combinations with nucleotide substitutions at position 6. Values in the boxes indicate the mean CFP expression from n = 3 biological replicates. Data are also shown in Figure 3g in main text. (**b**) Data from (panel a) shown as a bar plot. Bars represent average geometric mean CFP fluorescence of n = 3 biological replicates shown as dots. (**c**) Representative flow cytometry distributions from a single representative replicate for the samples shown in panel a and panel b. Samples are arranged in the same grid layout as in panel a. Io: non-complementary (decoy) input. The * indicates the WT sequence (Figure 1c).

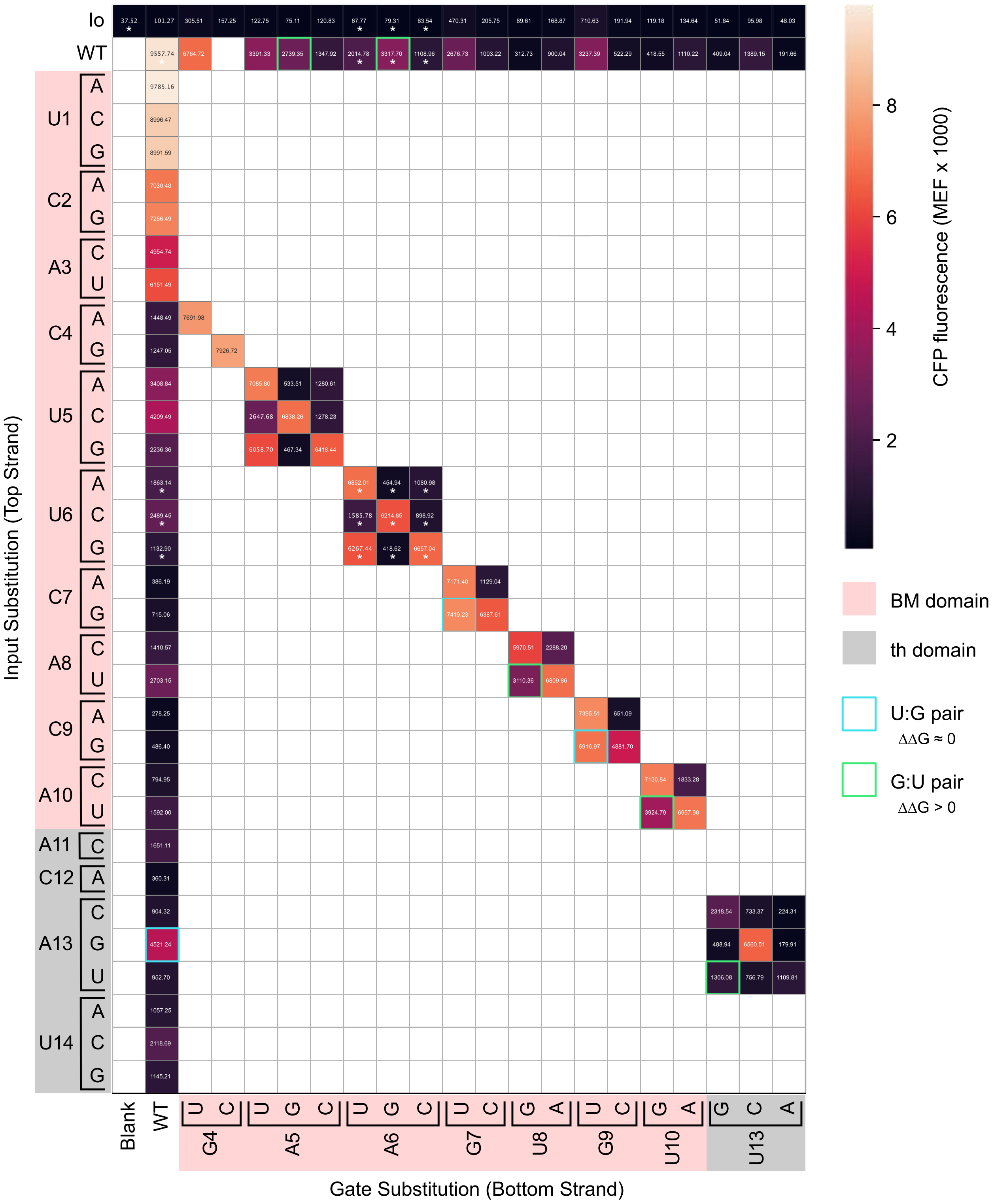

**Supplementary Figure 26**: All measurements of input-gate variant pairs in which either or both components have one substitution from the wild-type (WT) sequence (see main text Figure 1c).

*…Caption continued on the next page…*

Gate variants are shown on the x-axis, while input variants are shown on the y-axis. Substitution variants are labeled with the WT nucleotide at a given position outside the bracket and all measured nucleotide substitutions on the inside of the bracket. For example, the WT input has a U in position 1 and three variants were measured in which the U was substituted with either an A, C or G. The heatmap indicates the CFP signal produced by the corresponding input-gate pair (mean of n ≥ 3 independent replicates, measured by flow cytometry). The data in the 'WT' column, with the exception of A13G, are also shown in Figure 3e in the main text. Data from cells marked with an asterisk are also shown in Figure 3g in the main text. WT: wild-type (*i.e.*, canonical THE riboregulator or input sequence), Io: non-complementary (decoy) input, Blank: empty vector

Strong expression is observed for input-gate pairs with compensatory substitutions that produce a canonical base pair at the substitution position – this is seen along the diagonal of the matrix starting at row 9, column 3. For example, the gate in which the G nucleotide at position four has been substituted for a U (gate substitution G4U) paired with the input in which the C at position four has been mutated to an A (input substitution C4A) produces a U:A base pair after strand exchange, and consequently produces a strong CFP signal. Gate substitutions U13G and U13A are two exceptions to this pattern; these gates produce relatively little signal even when paired with input containing compensatory substitutions. As discussed in the main text, we hypothesize that this is due to new interactions between the toehold domain (where position 13 substitutions are introduced) and the ribozyme leading to misfolding that could prevent ribozyme cleavage or impede initiation of strand exchange (Supplementary Figure 27).

In contrast, input-gate pairs with substitutions that do not form a canonical base pair (shown in all squares that do not fall on the diagonal starting at row 9, columm3) generally produce a much weaker signal. Again, there are two notable exceptions to this pattern. The first, as discussed in the main text (Figure 3e), is the observation that inputs that introduce mismatches near the 5′ end of their sequence (position 1 to position 3) – near the end of branch migration route on the gate – still produce a strong signal. This aligns with measurements of DNA TMSE in the presence of mismatches in enzyme-free conditions[33,34]. The second exception is that input-gate pairs with substitutions that result in U:G or G:U wobble pairs at the substitution position also have a high degree of activation. This is expected, as these wobble pairs are known to be favorable in RNA duplexes. Notably, pairs in which there is a U on the gate and a G on the input (squares with a light blue outline) generally produce a greater CFP signal close to WT signal (6285 MEF on average across positions) than pairs in which there is a G on the gate and a U on the input (squares with a green outline, 1668 MEF on average across positions). We hypothesize that this is because, in the former case, prior to strand exchange the U on the gate is participating in a U:A base pair with the incumbent output strand. After strand exchange occurs, this base-pair is then exchanged for a U:G wobble base pair between the gate and input strand, which has a free energy comparable to that of the U:A pair it replaced. Thus, strand exchange in this case is approximately energetically neutral ($\Delta\Delta G \approx 0$ kcal/mol). In contrast, when a G is present on the bottom strand of the gate, prior to strand exchange it participates in a G:C pair. After strand exchange, it participates in a G:U pair, which is less thermodynamically favorable than a G:C pair ($\Delta\Delta G > 0$ kcal/mol), albeit still more favorable than most other non-canonical base pairs[35]. Thus, strand exchange in this case imposes a small energetic cost and therefore produces only a moderate signal.

In addition to highlighting the thermodynamic predictability of the effects of nucleotide substitutions on THE riboregulator performance, these measurements also provide further evidence that the decrease in signal observed when introducing a mismatch between input and gate are in fact a consequence of changes in strand exchange kinetics. The observation that the decrease in signal resulting from mismatch introductions between the gate and input (WT row/column) could consistently be recovered by introducing a compensatory substitution at the corresponding position of the complementary gate (*i.e.*, a substitution in the input could be compensated for by a complementary substitution in the gate and vice versa) strongly supports the model that mismatches attenuate strand exchange because of reduced strand exchange rates or altered equilibria rather than because of idiosyncratic effects introduced by changing input sequences such as altered transcription rates, premature termination, or RNA stability.

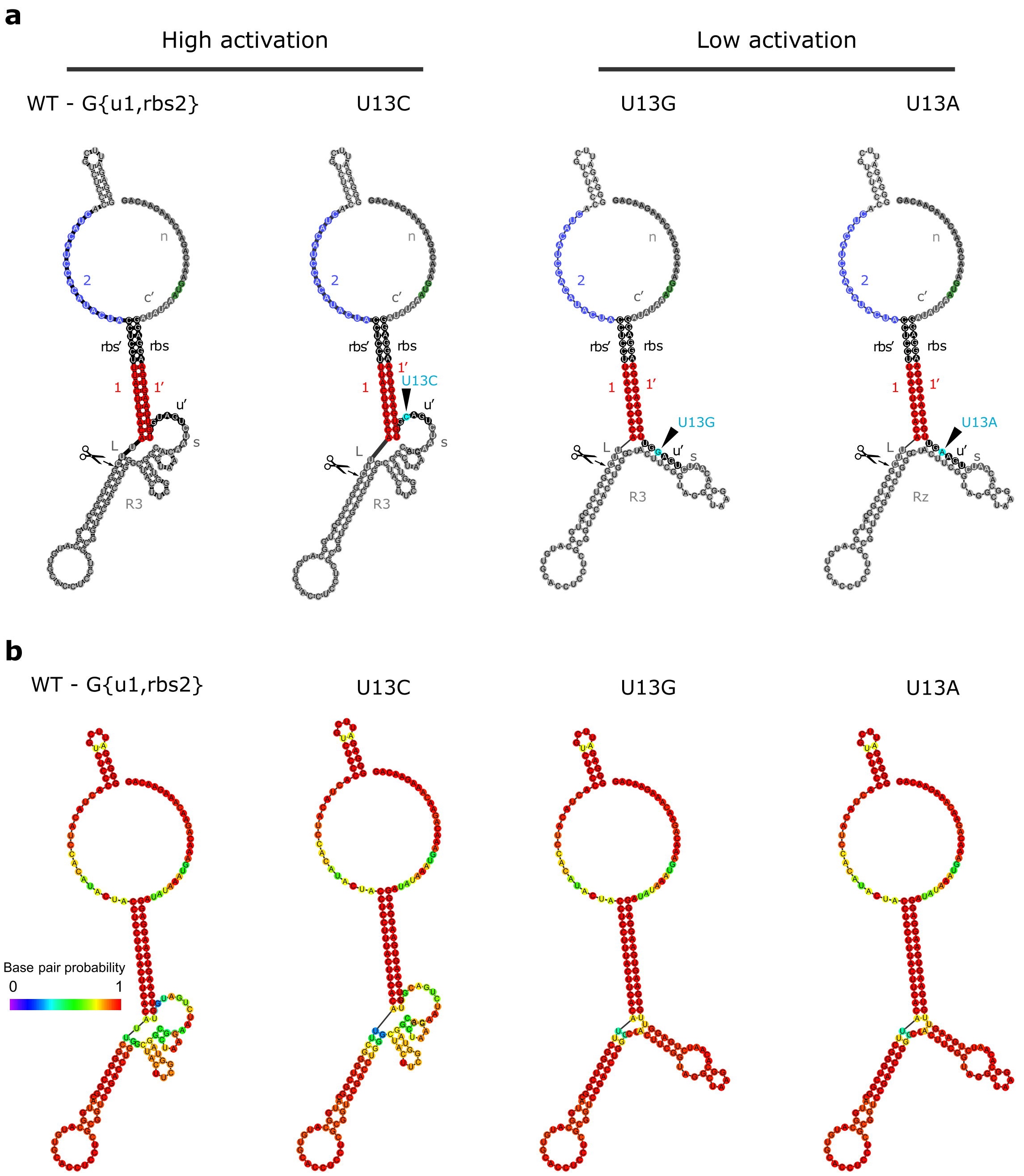

**Supplementary Figure 27**: **Substitutions in the input toehold of the gate could cause misfolding.** (**a**) Predicted secondary structures of the G{u1,rbs2} THE riboregulator gate and variants with a substitution in the toehold domain (position 13). Structures are colored by domain. Substituting the nucleotide U13 with a C (variant U13C) leads to a predicted structure that is similar to the WT structure. In contrast, substituting U13 with a G or and A introduces a favorable G:U or A:U pair to form between the substituted nucleotide and the ribozyme. This single base-pair is sufficient to form a dramatically different structure relative to wild type, in which the entirety of the input toehold domain (*u′*) forms the stem of a hairpin with 3′ end of the ribozyme. This structure may reduce the ability of the riboregulator to participate in toehold exchange by (i) occupying the toehold and slowing initiation of strand exchange and (ii) interfering with ribozyme self-cleavage. Indeed, variant U13C, when coexpressed with a complementary input, produced a signal comparable to WT (6561 and 8712 MFU, respectively), while variants U13G and U13A produced highly attenuated signals (2318 and 1100 MFU, respectively). (**b**) Same predicted structures as in panel a, but colored according to the predicted base pair probabilities. Predicted structures were generated by Vienna RNAfold[36].

### 8 Additional multi-component RNA circuit experiments

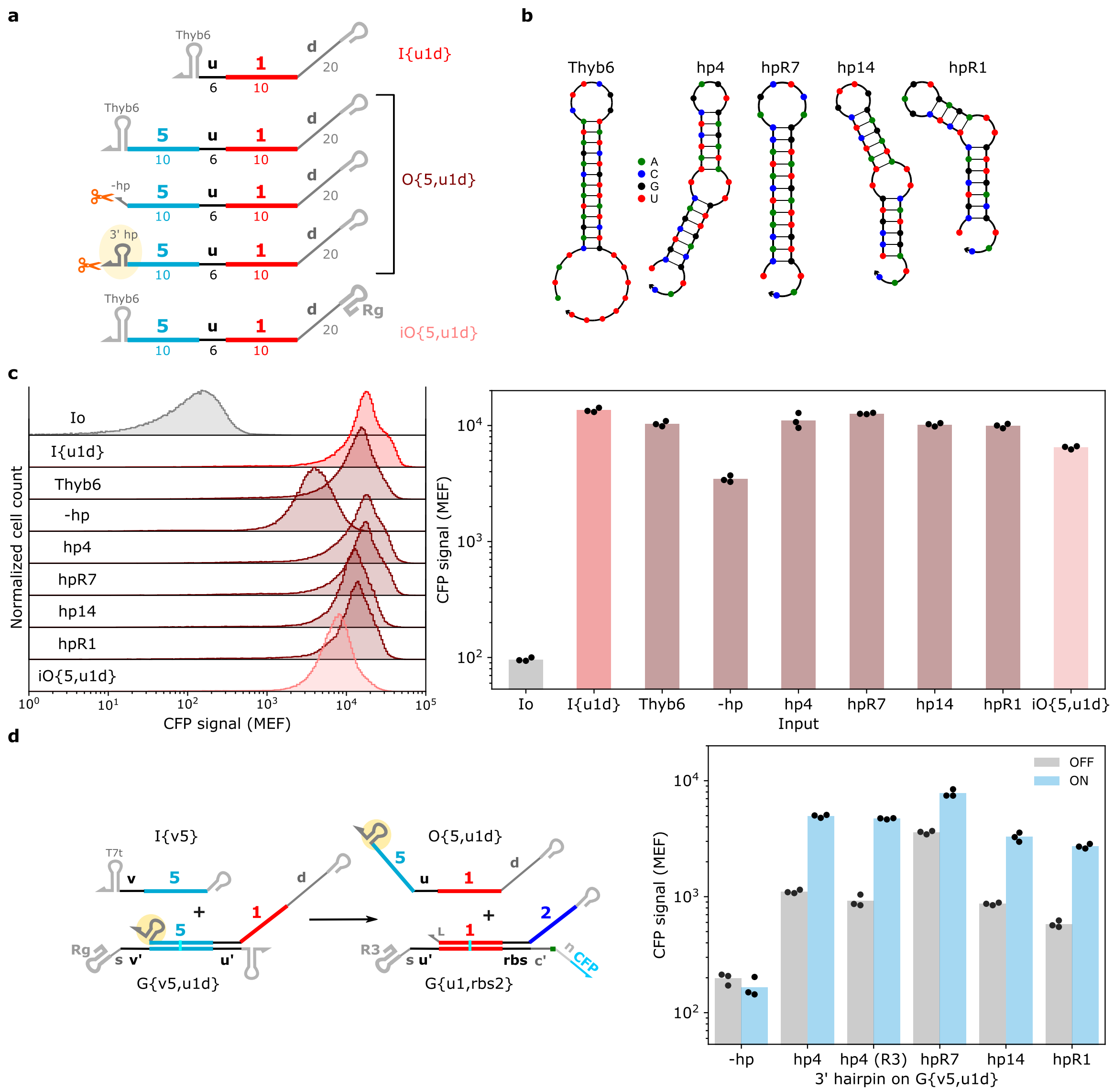

**Supplementary Figure 28**: **3′ hairpins on upstream gates increase protein expression in 2-layer cascades.** (**a**) Schematics of different output variants tested. The orange scissors indicate inclusion of the Rg ribozyme downstream to produce a 3′ end without a terminator hairpin. iO{5,u1d} represents the output of a gate with an inverted transcription order, *i.e.*, where the bottom strand of the gate is produced before the output strand[2]. (**b**) NUPACK-predicted secondary structures of the different 3′ hairpins tested. The hairpins hp4 and hp14 are from Ref[37] and hpR7 and hpR1 are from Ref[38]. (**c**) Flow cytometry results for the designs shown in panel a. The outputs were encoded on a pET backbone and the THE riboregulator was encoded on a pColA backbone (Supplementary Figure 7). (**d**) 2-layer cascades tested with different 3′ hairpins. The hp4 (R3) variant swapped in the R3 ribozyme sequence for the Rg sequence on G{v5,u1d}. I{v5} and G{v5,u1d} variants were encoded on a pET backbone) and the G{u1,rbs2} was encoded on a pColA backbone (Supplementary Figure 6). The hairpins hp4, hp14, and hpR1 were used to build multi-layer cascades.

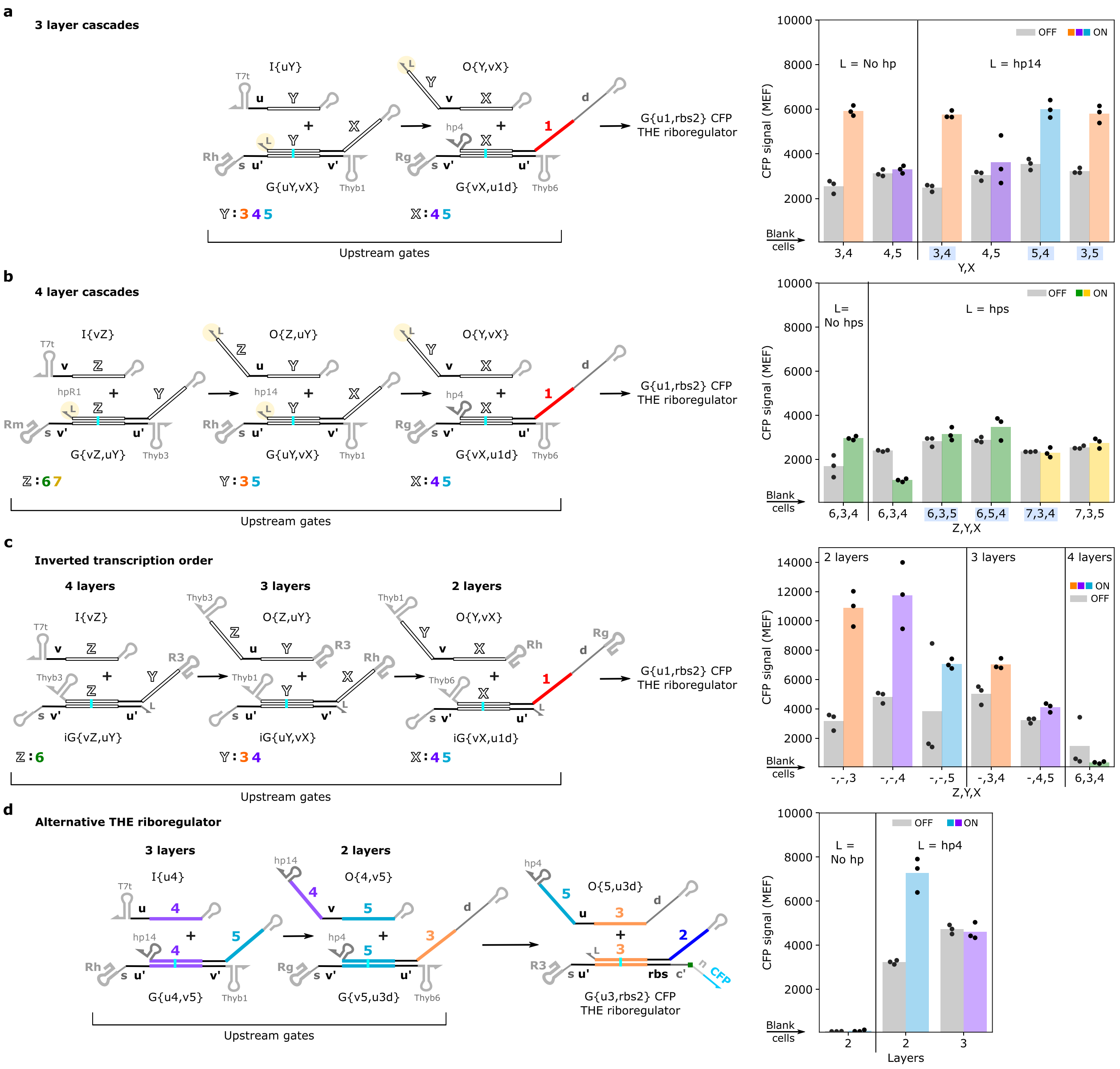

**Supplementary Figure 29**: **Multi-layer RNA strand exchange cascades**. (**a**) 2-layer, (**b**) 3-layer, and (**c**) 4-layer cascades using upstream gates with or without 3′ hairpins on their outputs. Cascades highlighted in blue on the x-axis of plots are also shown in Figure 3h of the main text. (**d**) Multi-layer cascades with upstream gates that have an inverted transcription order, *i.e.*, the bottom strand of the gate is produced before the output strand of the gate[2]. (**e**) Multi-layer cascades with a THE riboregulator that takes domain *3* as an input rather than domain *1*. Together, these results suggest a 3′ hairpin on the output is necessary for gates leading to the THE riboregulator, as all combinations of gates without this hairpin had low protein expression. However, the 3′ hairpin on the output does not appear to be strictly necessary for gates further upstream, as one 3-layer cascade (*Y*,*X*=*3*,*4*) had similar results with a gate in the third layer with and without the hairpin. G{u4,v5} appears to be broken as it does not work in any context tested.

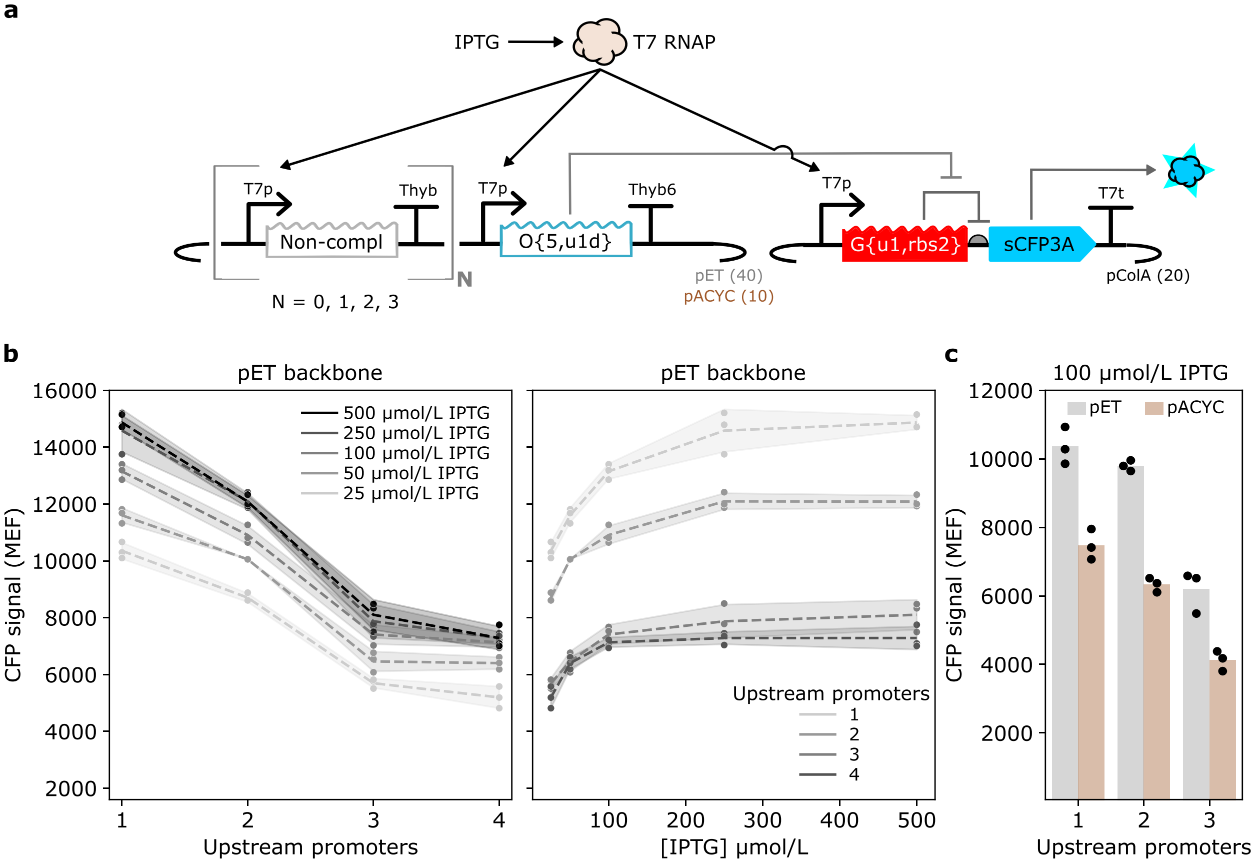

**Supplementary Figure 30**: **Increasing T7 RNAP expression does not alleviate the loss in signal as more promoters are added**. (**a**) Plasmid constructs for multi-promoter constructs, where N refers to the number of upstream promoters on the pET or pACYC plasmids. N=1, 2, 3 cassettes contain different non-complementary input sequences. G{u1,rbs2} has the *s4* spacer sequence in all experiments. See Supplementary Figure 7 for detailed schematics of each plasmid. (**b**) Flow cytometry results for the multi-promoter constructs shown in panel a with different concentrations of IPTG. IPTG was added to growing cell cultures 200 min after they were diluted from overnight cultures. Data points represent three independent replicates, lines represent means of the replicates, and shaded regions represent one standard deviation of the mean. *The left plot is also shown in Figure 4 of the main text, and the right plot shows a different visualization of the data in the left plot.* The sample 50 μmol/L IPTG, promoters = 2 sample only has one replicate. All other samples have three replicates. Increasing IPTG does increase CFP expression from each multi-promoter construct, but CFP expression still decreases as the number of promoters increases for each IPTG concentration. (**c**) The same trend of decreasing signal from a single layer cascade as upstream promoters are added was observed on a plasmid backbone with a lower copy number (pACYC).

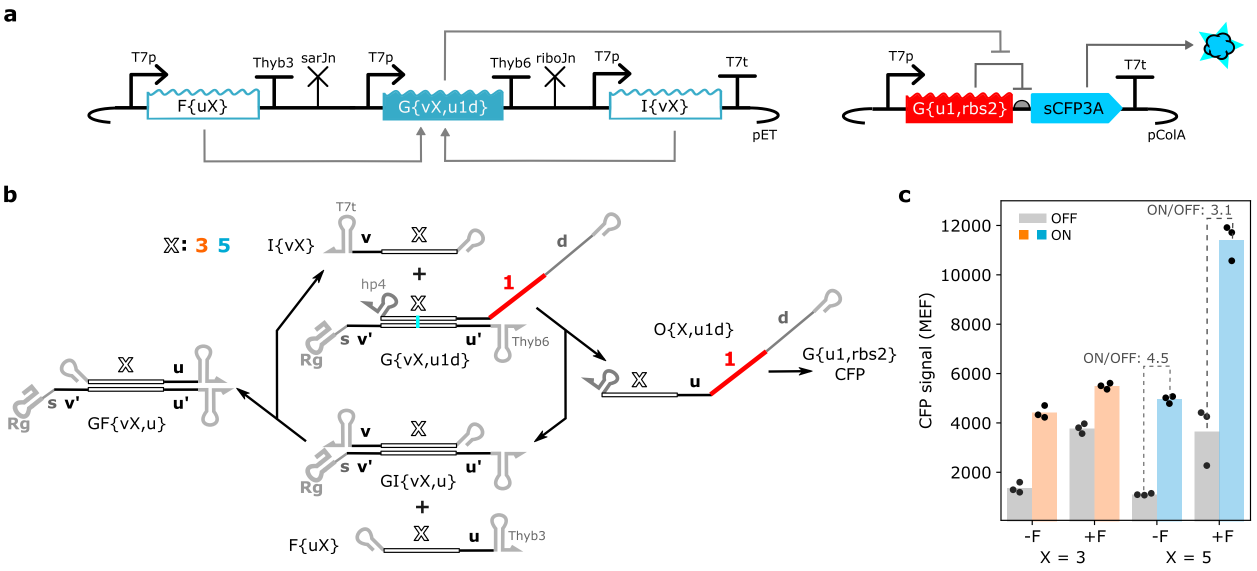

**Supplementary Figure 31**: **Amplifying second layer outputs with a fuel strand**. (**a**) Plasmid constructs for the fueled 2-layer cascade. (**b**) RNA interactions of the fueled 2-layer cascade. Note that the reaction with input and gate is reversible, likewise the reaction of fuel and the gate:input complex is also reversible. However, the output is consumed by the G{u1,rbs2}, which drives the reaction cycle in the direction shown by the reaction arrows[39]. (**c**) Flow cytometry results for two cascades without (-F) and with (+F) the fuel strand. Dashed gray lines in the plot with numbers above correspond to the ON/OFF ratios with and without fuel. Note that the fuel strand increases protein expression in both the ON and OFF states, resulting in a lower ON/OFF ratio than without the fuel strand.

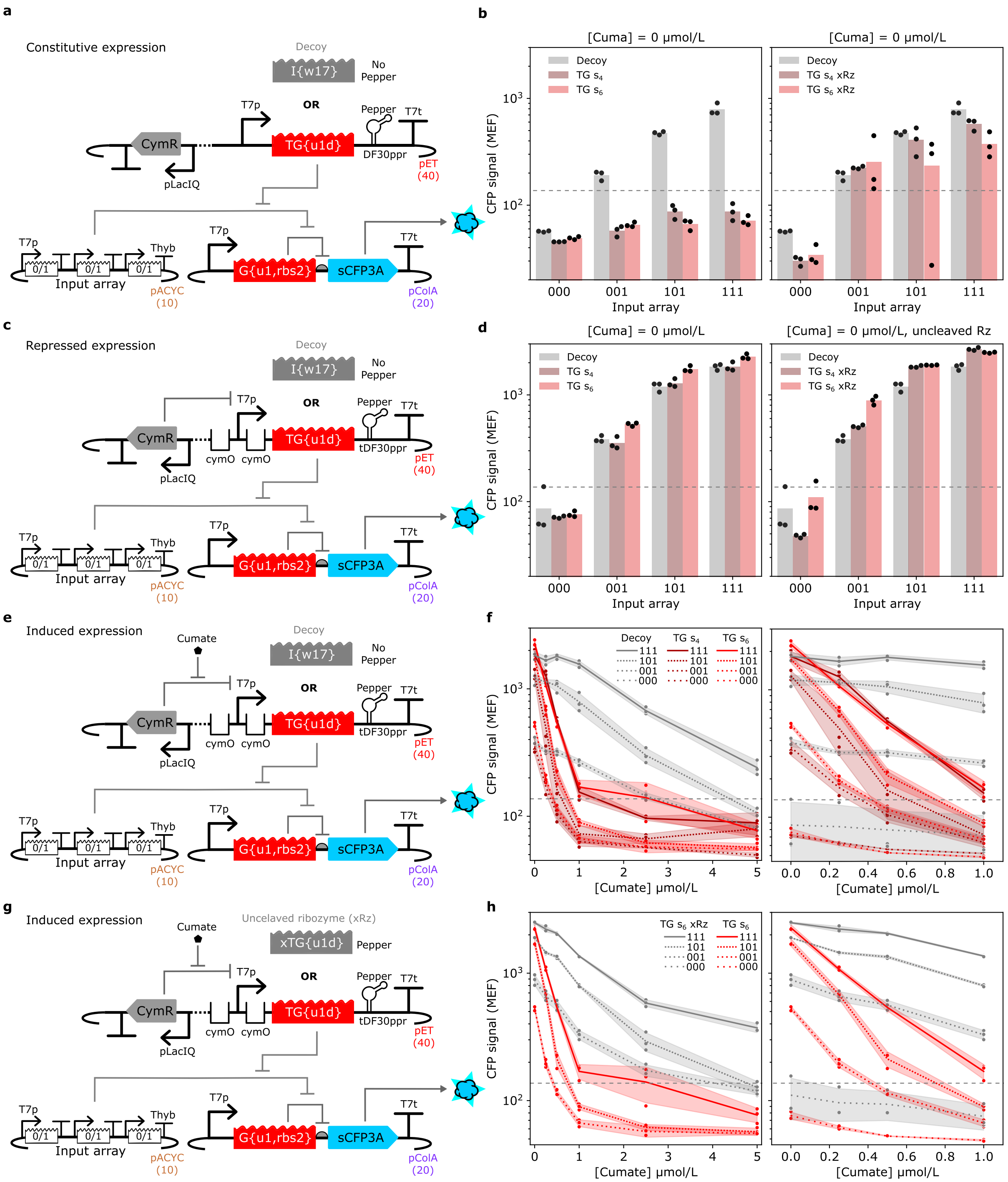

**Supplementary Figure 32**: **Validating threshold gate designs and the implementation of the classifier circuit.** Constitutive expression of threshold gates (panel a) sequesters inputs from reacting with the reporting gate (panel b, left). This sequestration is much less when threshold gates with ribozymes that cannot cleave (xRz) are used (panel b, right). CymR represses threshold gate expression (panels c,d). Cumate induces threshold gate expression (panel e), which sequesters inputs in a concentration dependent manner (panel f). Decoy expression at cumate concentrations > 1 µmol/L reduces CFP expression, indicating that increases in transcriptional load influence the system. Similar results are observed when the decoy sequence is replaced with a threshold gate that cannot cleave (panels g,h). However, these load-based reductions in expression are much less pronounced than when the active threshold gates are expressed. For cumate concentrations between (0 and 1) µmol/L, reduction in CFP signal is specific to threshold gate expression and is consistent with rapid input sequestration as threshold gates with ribozymes that cleave are necessary.

(**a**) Plasmid schematic for a 3-input logic element with constitutive expression of threshold variants. Four TG variants were tested: one with a 4-base spacer ($s_4$) and one with a 6-base spacer ($s_6$) as well as two with the same two spacers but with a mutant ribozyme incapable of cutting (xRz). (**b**) Flow cytometry results for the system shown in panel a with 0, 1, 2, or 3 inputs. The three number codes on the x-axis indicate which positions of the input array contained sequence-complementary inputs (1) vs non-complementary inputs (0). Horizontal gray dashed lines indicate the cutoff distinguishing ON vs OFF. Constitutive expression of thresholds effectively annihilates inputs while expression of a decoy RNA does not. (**c**) Plasmid schematic for a 3-input logic element with CymR repression of threshold variants. (**d**) Flow cytometry results for the system shown in panel c. CymR sufficiently represses a T7 promoter flanked by CymR operator sequences. (**e**) Plasmid schematic for a 3-input logic element with cumate-inducible expression of threshold variants. (**f**) Flow cytometry results for the system shown in panel. Data points represent three independent replicates, lines represent means of the replicates, and shaded regions represent one standard deviation of the mean. (**g**) Plasmid schematic for a 3-input logic element with cumate-inducible expression of the cleaved and uncleaved (xRz) threshold with a 6-base spacer. (**h**) Flow cytometry results for the system shown in panel g. The data for TG $s_6$ is replotted from panel f for comparison. Note the threshold gates contained a dimeric Pepper aptamer construct upstream of their terminator, but fluorescent aptamer measurements were not conducted. However, the presence of the aptamer may influence RNA stability. The scaffold for the Pepper aptamers was different for constitutively expressed threshold gates compared to threshold gates under CymR control (Supplementary File S1).

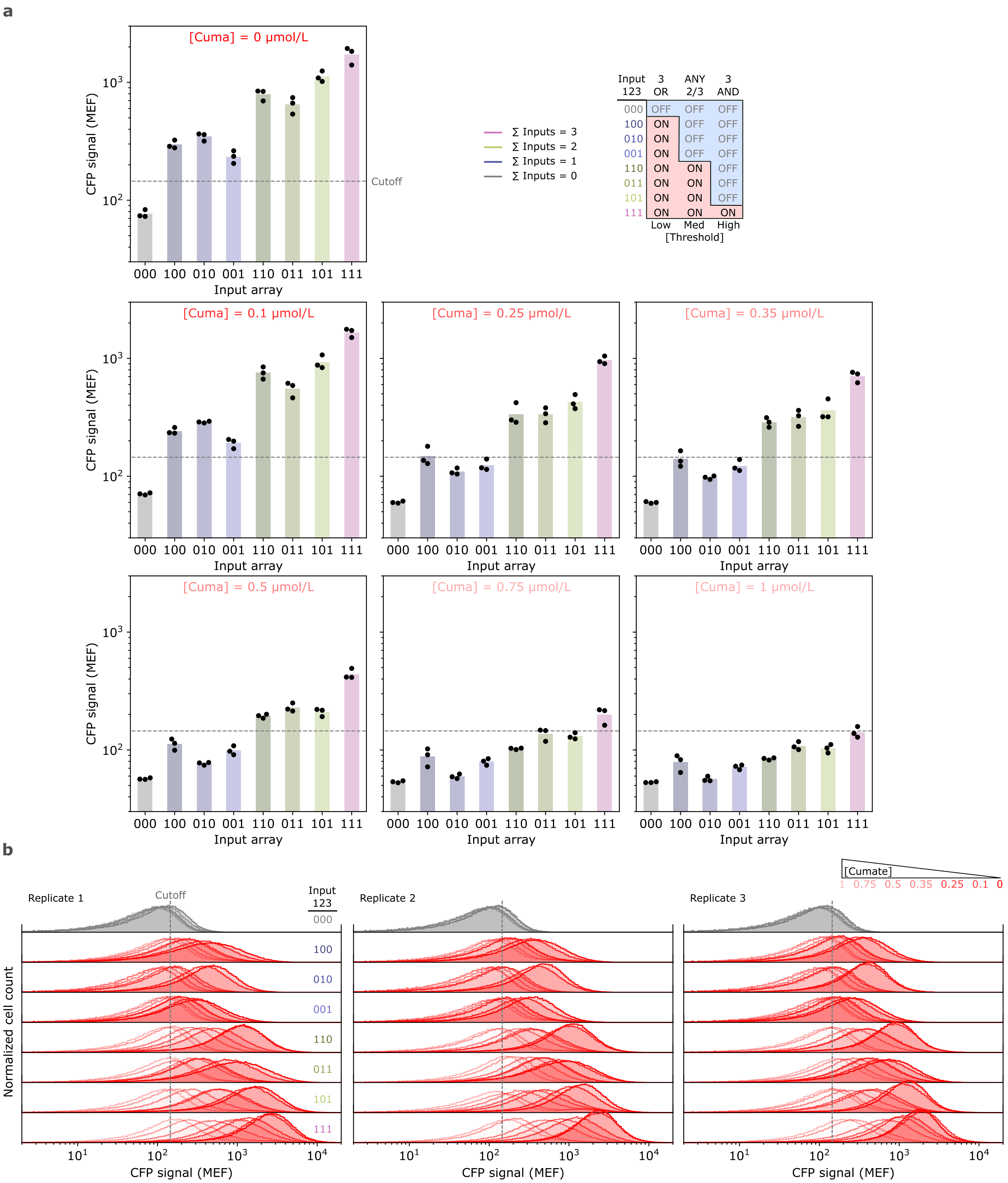

**Supplementary Figure 33**: **Flow cytometry histograms for the classifier circuit.** (**a**) Bar plots of the results shown in Figure 5 of the main text. (**b**) Flow cytometry histograms of the results shown in panel a and Figure 5 of the main text. The geometric means of these histograms represent the data points in panel a for three biological replicates. The gray dashed lines in both panel a and panel b indicate the cutoff value between the ON and OFF states.

#### 9 Experiments with different fluorescent protein coding sequences

See Supplementary Section 2.2 for design considerations and details for how these protein coding sequences were designed.

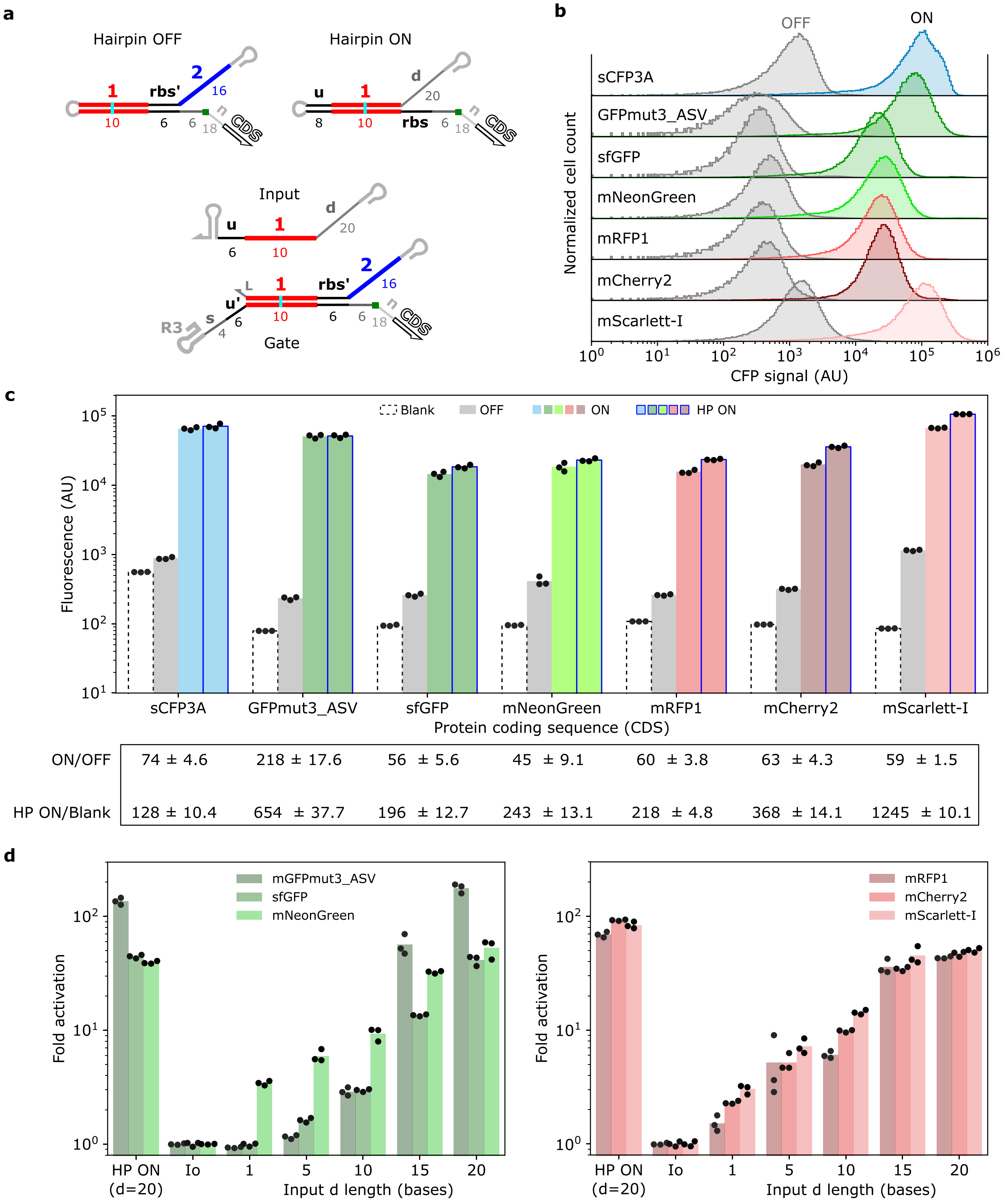

**Supplementary Figure 34**: **Regulating expression of additional fluorescent protein outputs with THE riboregulators.** … *Caption continued on next page…*

(**a**) Schematics of RNA components used to test additional fluorescent proteins. The *s* domain was $s_4$ for THE riboregulators encoding all proteins other than GFPmut3_ASV, which used $s_4u$. Additionally, the GFPmut3_ASV sequence encodes for a C-terminal degradation tag[18]. ASV degradation tags were also tested with sCFP3A in Supplementary Figure 35. (**b**) Representative flow cytometry distributions for THE riboregulators expressing different fluorescent proteins. (**c**) Geometric means of flow cytometry distributions for the designs depicted in panel a. The excitation laser, emission filters, and channel gains differed across proteins (Methods) so the absolute values across proteins are not strictly comparable. The table below the bar plot show the mean ON/OFF ratios from the bar plot (± one standard deviation) for each fluorescent protein, indicating the dynamic range for each riboregulator. The mean HP ON/Blank cell ratios are also shown, indicating the maximum possible dynamic range for each system under our measurement conditions. (**d**) Flow cytometry results for inputs with increasing *d* domain lengths and THE riboregulators controlling a variety of fluorescent proteins indicate the precise control of protein expression using strand exchange transfers to different coding sequences. These results are normalized by dividing by the mean value of the Io sample for each fluorescent protein, which represents the fold activation of the samples above the OFF.

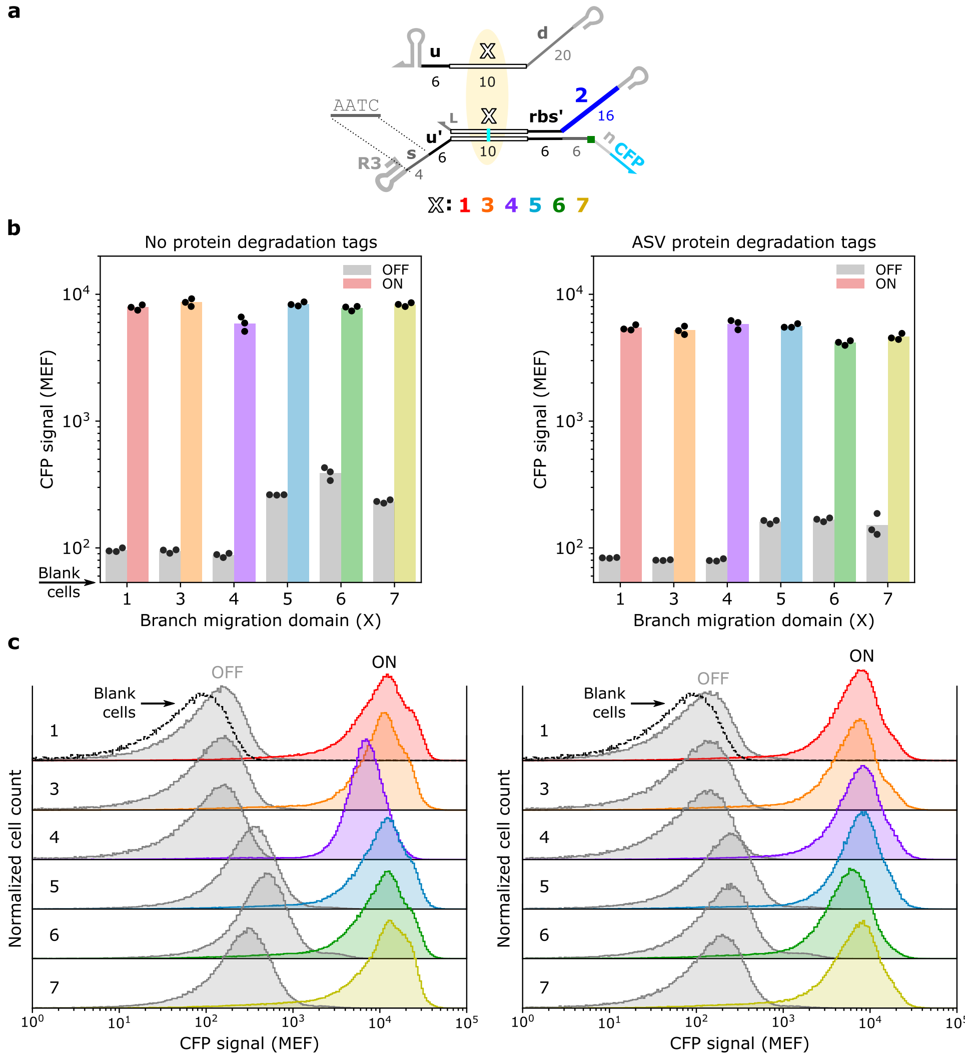

**Supplementary Figure 35**: C-terminal ASV degradation tags[18] moderately reduce protein expression with and without the complementary RNA input. The results without the ASV tag are also shown in Figure 3 of the main text and are repeated here for comparison.

#### 10 DHFR growth assay

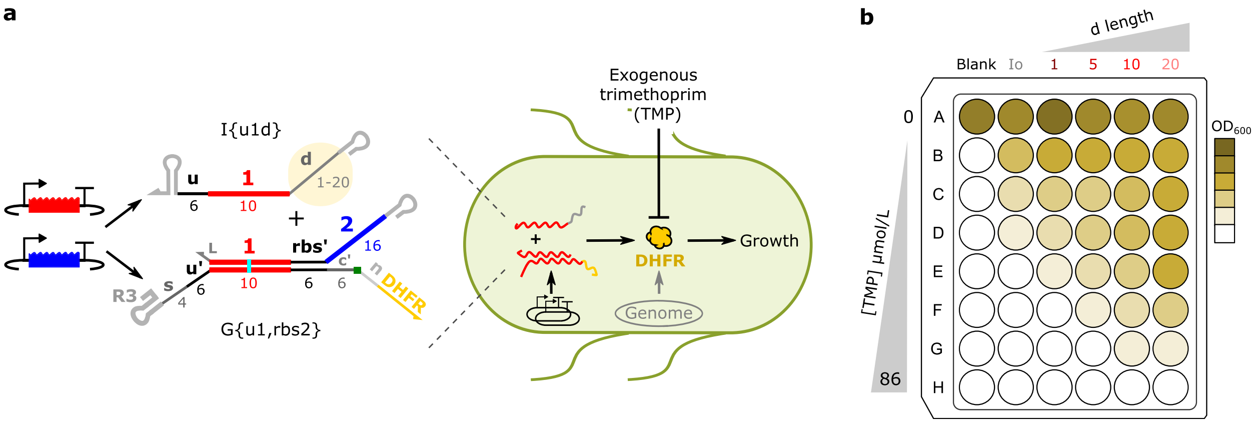

**Supplementary Figure 36**: Experimental set up of the DHFR growth assay. (**a**) Schematic of the system. Controlling expression of a copy of *E. coli* dihydrofolate reductase (DHFR) with RNA strand exchange enables control of cell fitness across antibiotic concentrations. DHFR is an essential gene required for DNA replication and protein production – and thus bacterial growth. *E. coli* DHFR is inhibited by the antibiotic trimethoprim (TMP), preventing growth. (**b**) Illustration of the experimental plate layout along with expected results. Increasing the length of the *d* domain on the input strand should increase the translation rate of DHFR enabling cells to grow at higher TMP concentrations, indicated by a higher optical density at 600 nm (darker $OD_{600}$).

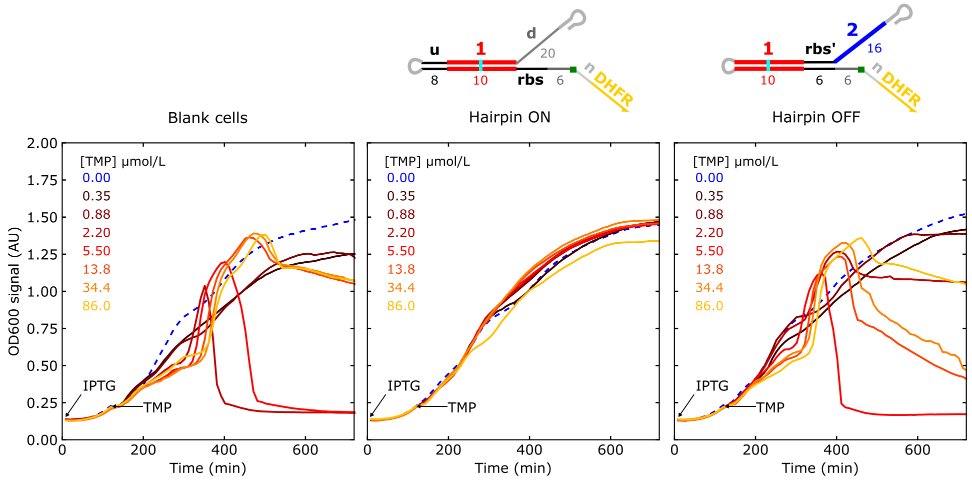

**Supplementary Figure 37**: The DHFR growth assay produced strange growth curves in rich LB medium. Other than the different growth medium these experiments were conducted as described in the methods and Supplementary Figure 36.

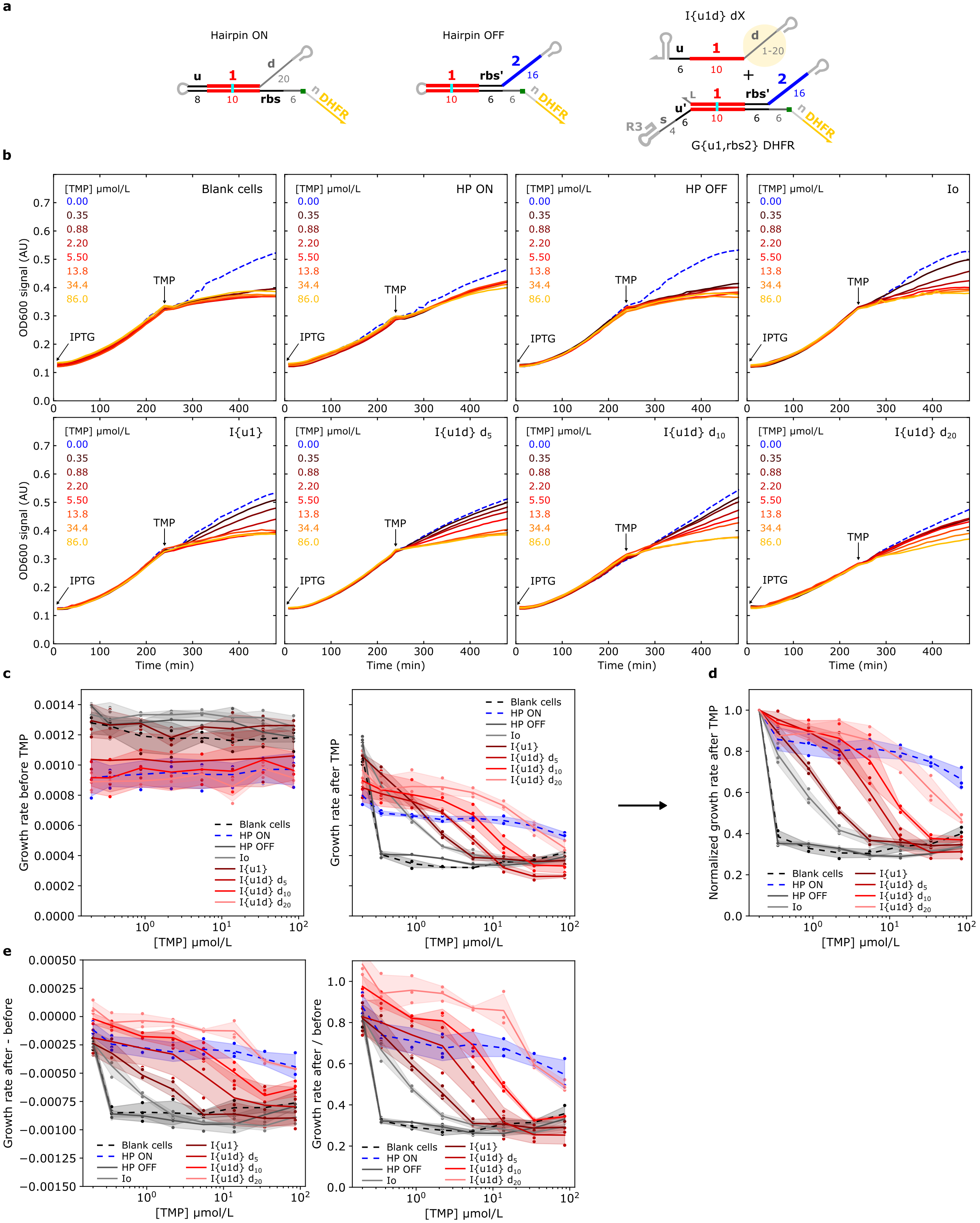

**Supporting Figure 38**: Growth curves and analysis for the DHFR growth assay. (**a**) Schematics of RNA components used in the assay. Hairpin ON and hairpin OFF constructs serve as controls for maximum and minimum DHFR expression, respectively. These samples also included a pET plasmid encoding Io to mimic the two-plasmid system used for the THE riboregulators. … *Caption continued on next page…*

(**b**) Optical density measurements at 600 nm over the course of the growth assay for the designs in panel a. At time = 0 min, overnight cultures of cells were diluted 50-fold into M9 medium supplemented with 100 µmol/L of IPTG to induce RNA expression. At time = 240 min, TMP was added to the cultures to the concentrations specified in the legend of the plots. The optical density measurements over the next 90 min (240 min to 330 min) were used to fit a line to obtain a growth rate at each TMP concentration. (**c**) Growth rate of strains before (right) and after (left) TMP addition. Growth rates before and after were determined by fitting a line to OD600 measurements 90 min before and 90 min after TMP addition, respectively. (**d**) Growth rate after TMP addition was normalized by dividing each curve by the growth rate of each strain without any TMP present. This data is presented in Figure 6 of the main text. (**e**) Alternative analysis approaches that yield qualitatively similar results. Using the growth rates in panel c, the left plot was produced by subtracting the growth rates before and after TMP addition and the right plot was produced by dividing the growth rates before and after TMP addition.

#### 11 THE riboregulator expression in different genetic contexts

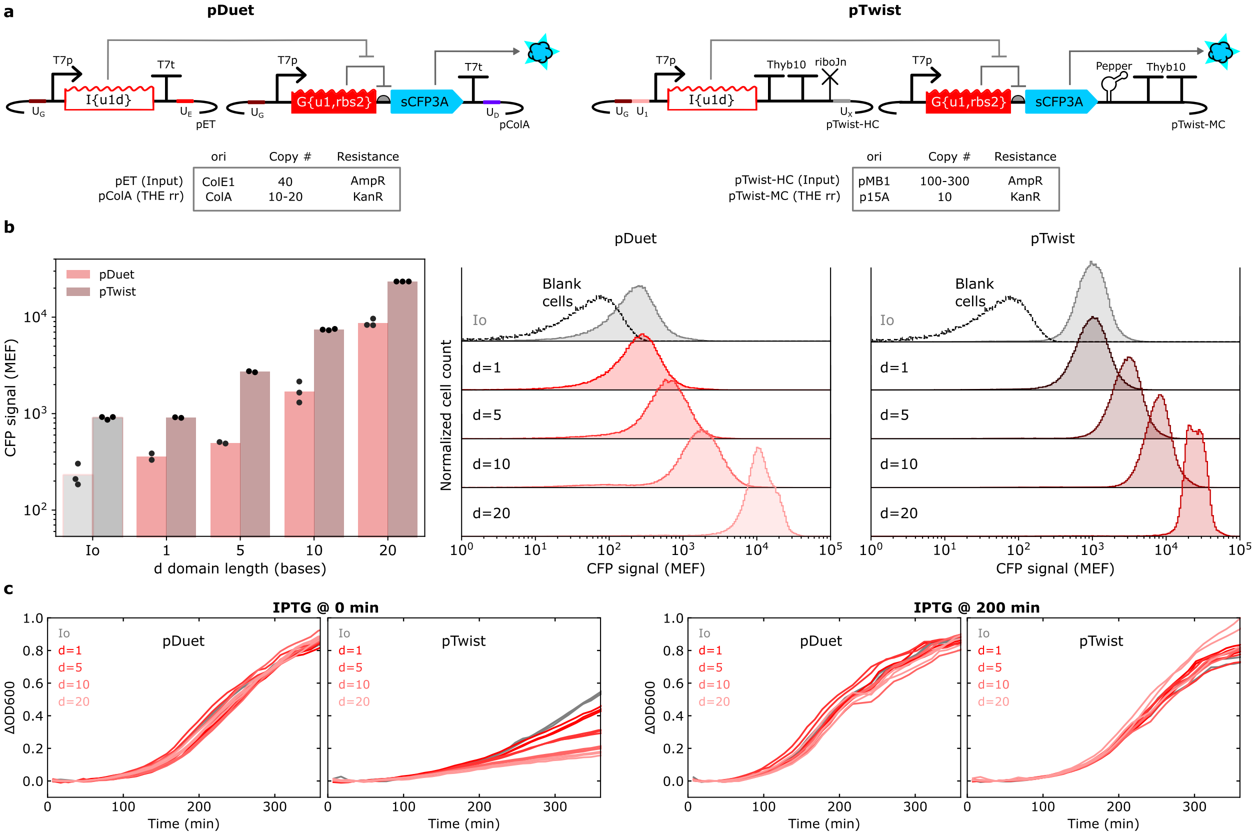

**Supplementary Figure 39**: Testing THE riboregulators in different plasmid backbones. (**a**) Schematics of THE riboregulators encoded on pDuet backbones or pTwist backbones. In addition to the origin of replication and antibiotic resistance genes, both pDuet plasmids encode a copy of LacI (Supplementary Figure 4). The pTwist THE riboregulator also included a dimeric Pepper RNA aptamer on a tRNA scaffold[23]. (**b**) Flow cytometry data comparing CFP expression levels from the two plasmid sets in panel a. The pTwist constructs produce more signal across all d domain lengths, as well as with the non-complementary input. The results in Supplementary Figure 40 suggest this is not entirely due to the difference in the 3′ UTRs between the pDuet RNAs and the pTwist RNAs. It is possible that the additional copies of LacI on the pDuet backbones suppress basal T7 RNAP expression levels and reduce the amount of T7 RNAP expressed at 100 µmol/L IPTG compared to the pTwist backbones. (**c**) Growth curves of cells containing the different plasmids sets in panel a. Lines of the same color are biological replicates. Plots on the left represent cells that were induced with IPTG at t = 0 min (immediately after a 50-fold dilution of overnight cultures). Plots on the right represent cells that were induced with IPTG at t = 200 min after dilution of the overnight cultures. The results in panel b were from cells induced with IPTG at 200 min after dilution of the overnight cultures. The pTwist constructs induce more cellular stress when inducing directly after overnight culture dilution. This could be due to an accumulation of the high copy plasmid in stationary phase cells[26].

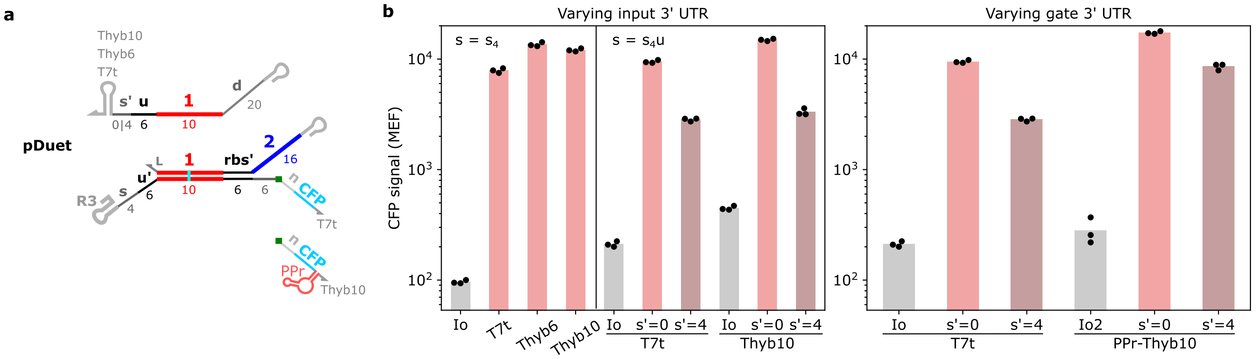

**Supplementary Figure 40**: Testing THE riboregulators with different 3′ untranslated regions (UTRs). (**a**) Schematic of THE riboregulator components tested with different 3′ UTR shown. (**b**) Flow cytometry results for inputs with different terminators and a THE riboregulator with the native T7 RNAP terminator (T7t) (left) and THE riboregulators with different 3′ UTRs and an input with T7t (right). In both cases, inputs with either 0-base or 4-base complementarity with the *s* domain on the THE riboregulator were tested. The same trend as in Supplementary Figure 19 hold with different 3′ UTRs – a 6-base toehold on the input RNA is better than a 10-base toehold.
